# Decoding RNA–RNA Interactions: The Role of Low-Complexity Repeats and a Deep Learning Framework for Sequence-Based Prediction

**DOI:** 10.1101/2025.02.16.638500

**Authors:** Adriano Setti, Giorgio Bini, Flaminia Pellegrini, Valentino Maiorca, Gabriele Proietti, Dimitrios-Miltiadis Vrachnos, Angelo D’Angelo, Alexandros Armaos, Julie Martone, Michele Monti, Giancarlo Ruocco, Emanuele Rodolà, Irene Bozzoni, Alessio Colantoni, Gian Gaetano Tartaglia

## Abstract

RNA–RNA interactions (RRIs) are fundamental to gene regulation and RNA processing, yet their molecular determinants remain unclear. In this work, we analyze several large-scale RRI datasets and identify low-complexity repeats (LCRs), including simple tandem repeats, as key drivers of RRIs. Our findings reveal that LCRs enable thermodynamically stable interactions with multiple partners, positioning them as key hubs in RNA–RNA interaction networks. These RRIs appear to be important for several aspects of RNA metabolism. Sequencing-based analysis of the lncRNA Lhx1os interactors validates the importance of LCRs in shaping contacts potentially involved in neuronal development. Recognizing the pivotal role of sequence determinants, we develop RIME, a deep learning model that predicts RRIs by leveraging embeddings from a nucleic acid language model. RIME outperforms traditional thermodynamics-based tools, successfully captures the role of LCRs and prioritizes high-confidence interactions, including those established by lncRNAs. RIME is freely available at https://tools.tartaglialab.com/rna_rna.

## Introduction

RNA molecules play key roles in cellular function, not only as carriers of genetic information but also as regulators of gene expression, structural components of molecular complexes, and guides for enzymatic processes. These functions often rely on RNA–RNA interactions (RRIs), which are central to diverse processes such as transcriptional and translational regulation, RNA splicing, stability control, and the formation of ribonucleoprotein complexes^1–5^.

In recent years, technological advances have enabled systematic mapping of RRIs, unveiling their prevalence and complexity. High-throughput methods such as PARIS^6,7^, SPLASH^8^, and LIGR-Seq^9^ combine Psoralen-based chemical crosslinking with sequencing to capture base-paired regions in RNAs. These techniques allow the detection of interactions at near-base resolution, revealing structural motifs and interaction networks within and between RNA molecules. Additionally, emerging methods such as KARR-Seq provide a complementary approach by capturing RNA interactions with the aid of click chemistry reactions^10^. Protein-based approaches, including RIC-seq^11^ and MARIO^12^, broaden the scope of RNA interactome studies by focusing on protein-facilitated RNA–RNA interactions. Together, these approaches have provided unprecedented insights into RNA interactomes, highlighting the pervasive role of RNA–RNA interactions in cellular regulation.

Despite these technological advances, the molecular determinants governing RRIs remain poorly understood. Sequence complementarity and RNA secondary structure are known to play fundamental roles^13,14^, but recent findings suggest that specific sequence motifs and repeat elements contribute significantly to interaction specificity^15–17^. For instance, the A-repeat region of Xist long non-coding RNA (lncRNA) forms a series of inter- and intra-repeat bindings, suggesting that these interactions are vital for the multimerization necessary for Xist function^18,19^. Similarly, satellite repeats in a subset of the mRNAs targeted by miR-181 provide binding platforms for the miRNA; while miR-181 typically induces mRNA degradation, its interaction with these repeats is instead associated with translational repression^20^. Tandem repeats in the TERT gene allow to establish base pairing interactions between distal regions of the pre-mRNA, which are necessary for exons 7 and 8 skipping^21^. Interspersed repeat elements, such as SINEs, also play a significant role in RNA–RNA interactions. For example, Staufen 1 (STAU1) protein has been shown to recognize imperfect duplexes formed between pairs of Alu elements (primate-specific SINEs) embedded within a lncRNA and the 3’UTR of a target mRNA, leading to mRNA decay^22^. Collectively, these examples suggest a conserved mechanism by which repeat elements enhance RNA–RNA interactions, contributing to their specificity, stability, and functional versatility. However, a systematic study on the involvement of repeated elements in the establishment of RNA–RNA interactions is still lacking.

To complement experimental approaches, computational tools have emerged as powerful resources for predicting RRIs. Algorithms relying on thermodynamic models (e.g., RNAduplex and IntaRNA^23–25^) are effective for short RNA interactions but face limitations in capturing the full complexity of long RNA molecules and their in vivo dynamics^26^. Recent advances in neural networks^27^, particularly deep learning^28^, have created opportunities to enhance the prediction of both intra-and inter-RNA interactions. These developments harness large-scale datasets and diverse sequence encoding strategies to improve accuracy and broaden predictive capabilities. In particular, sequence representations generated by nucleic acid language models have demonstrated remarkable effectiveness in encoding intricate and informative sequence features.^29,30^. By combining high-throughput training data with advanced sequence representations, it becomes possible to develop predictive models that effectively capture the complexity of long RNA molecules and their interactions.

In this study, we investigated sequence determinants underlying the interactions between long RNA molecules, retrieved from several high-throughput datasets. Strikingly, we discovered that the interacting regions are enriched in simple and low-complexity repeats. These repetitive elements are associated with thermodynamically stable interactions and are often found in regions with a high degree of connectivity, suggesting a potential role as interaction hubs. Notably, low-complexity repeats co-occur with interactions between transcripts involved in neuronal development. Recent research has highlighted the widespread upregulation of lncRNAs containing simple repeats during neuronal differentiation^31^. To explore this aspect further, we sequenced the RNA interactors of a lncRNA critical for motor neuron development. Our results showed that these interactors are enriched in low-complexity repeats that are likely to be involved in forming energetically favorable interactions with the lncRNA.

The discovery of these sequence features prompted us to model RNA–RNA interactions using a sequence-encoding strategy based on Language Models (LMs). To this end, we developed RIME, a deep learning-based predictor trained on Psoralen-based datasets. RIME outperforms thermodynamics-based tools, even when evaluated against interactions identified using protein-based methods. Notably, RIME achieves excellent performance on high-quality interaction datasets and demonstrates a clear ability to discern the importance of low-complexity repeats in RNA–RNA interactions. Our study provides the first systematic benchmarking of thermodynamics-based tools against RNA–RNA interactomes generated by high-throughput technologies.

## Results

### Preparation of a transcriptome-wide RNA duplexes dataset

We derived the main RRI dataset used in this study by collecting the results of five transcriptome-wide techniques that were developed to massively identify RNA-duplexes through RNA-sequencing: PARIS1, PARIS2, and SPLASH identify intramolecular and intermolecular helices crosslinked with Psoralen or its derivatives (Psoralen-based techniques), while RIC-Seq and MARIO detect RNA–RNA interactions mediated by proteins (protein-based techniques, **Supplementary Fig. 1a**). Overall, we collected a total of 7,813,978 RNA duplexes derived from both mouse (mESC and mouse brain samples) and human systems (HEK293T, HeLa, Lymphoblastoid and ES cells treated or not with Retinoic Acid) (**Supplementary Data 1**). Since our aim was the characterization of interactions between long RNA molecules that are potentially involved in post-transcriptional gene expression regulation, we focused on duplexes derived from exonic RNA regions that are more likely to arise from processed transcripts (**Supplementary Fig. 1b** and c), and we discarded duplexes formed within the same transcripts, probably resulting from intramolecular RNA–RNA pairing (**Supplementary Fig. 1b** and c). Moreover, we removed interactions involving particularly abundant RNA species, which are most likely driven by their stoichiometric imbalance rather than by the specificity of RNA–RNA contact. These species include the ribosomal RNA, which directly contacts the mRNAs during translation. We also filtered out interactions carried by small RNAs, such as miRNAs and snoRNAs (**Supplementary Fig. 1b** and c). After these filters, we reduced our pool of RNA duplexes to 117,948 inter-molecular interactions (**Fig. 1a**, **Supplementary Data 1**), which account for 1.5% of the initial set (**Supplementary Fig. 1d**). Among these duplexes, the majority consist of mRNA pairs, with a significant proportion represented by 3’UTR-CDS and CDS-CDS interactions. Interactions between non-coding RNAs and mRNAs, particularly those involving 3’UTRs, are also well represented, with the exception of the SPLASH RRI sets, which are almost exclusively composed of interactions between protein-coding transcripts (**Supplementary Fig. 1e**).

**Fig. 1.**
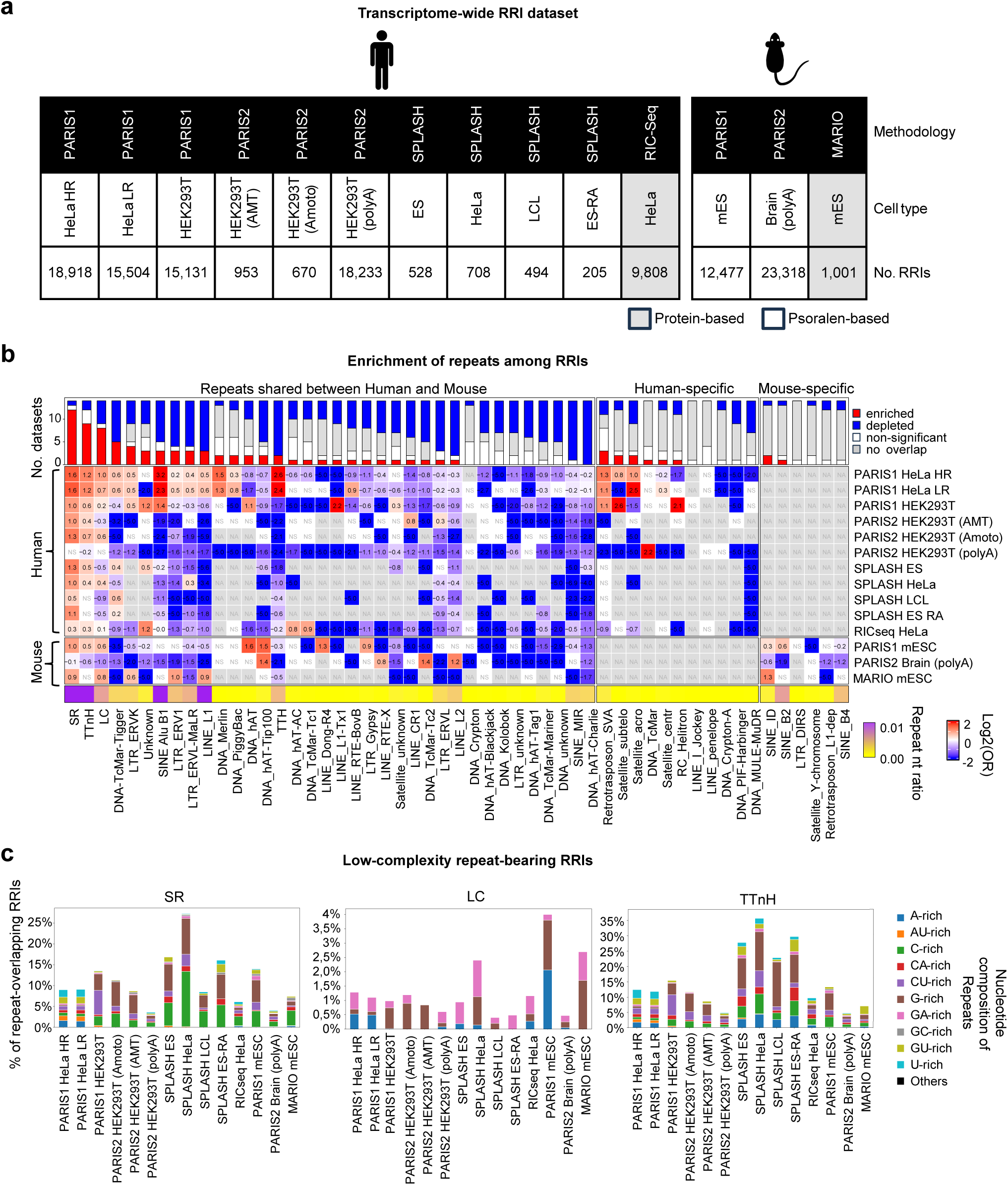
RNA>–RNA interaction landscapes reveal a predominance of low-complexity repeats. **a**, Schematic representation of the RRIs derived from transcriptome-wide experiments that were employed in this work. The first row describes the RRI detection methodology; the second row describes the organism and the third one reports the cellular system; the fourth row reports the amount of intramolecular RNA–RNA interactions used in the present study. Columns are colored based on the type of the RRI detection methodology: Psoralen-based or protein-based. **b**, Heatmap depicting, for each transcriptome-wide experiment (rows), the log_2_(Odds Ratio) representing the nucleotide enrichment of repeat families (columns) in RRIs. Significant enrichment or depletion of repeats is indicated by red or blue colors, respectively; white (NS) denotes the absence of statistical significance, while gray (NA) indicates no nucleotide overlap. Statistical significance was assessed using two-sided Fisher’s exact test with Benjamini-Hochberg correction for multiple testing. The bar plot above the heatmap shows, for each tested repeat family, the number of transcriptome-wide experiments where a statistically significant association between repeats and RRIs was found (red for enrichment, blue for depletion) or not (gray). The color bar below the heatmap indicates the proportion of repeat nucleotides involved in RRIs relative to the total RRI nucleotides, averaged across all experiments, using a color scale from yellow (low frequency) to violet (high frequency). **c**, Stacked bar plot reporting, for each transcriptome-wide experiment, the percentage of RRIs overlapping with SR (left), LC (middle) or TTnH (right), categorized by their nucleotide composition. The colors in the legend describe the nucleotide composition categories used to classify repeated elements.

### RNA–RNA interactions detected by transcriptome-wide experiments are enriched in low-complexity repeats

After dataset preparation, we investigated the relationship between RRIs and repeated elements. We used RepeatMasker software^32^ to identify known repeated elements in the 28,393 and 16,559 interacting RNAs detected in human and mouse samples, respectively. Human interacting RNAs contain 76,799 repeats belonging to 45 different families, while the murine ones harbor 43,825 repeats from 40 families (**Supplementary Fig. 1f**). To assess whether some repeat families are preferentially involved in RRIs, we performed a nucleotide-level repeat enrichment analysis (**Supplementary Fig. 1g**, see **Methods**), testing for the enrichment or depletion of repeat nucleotides in the interacting RNA regions, while controlling for their overall abundance in transcripts. We found that, while most of the repeat classes displayed depletion or sporadic enrichment in RNA–RNA interacting regions, two classes stood out with widespread enrichments among datasets: Simple Repeats (SRs) and Low Complexity (LC) sequences. Indeed, the RRI identified in 12 and 8 (out of 14) experiments were significantly enriched (log_2_[Odds Ratio] > 0 and FDR < 0.05) in SRs and LCs, respectively (**Fig. 1b**). Moreover, to confirm that the enrichment of these repeat families was not dependent on the software used to annotate repeated elements, we also employed the TanTan algorithm^33^, observing that tandem repeats (not homopolymeric, TTnotHomo, see **Methods**) are enriched in RNA–RNA interactions from 9 experiments (**Fig. 1a**). We grouped the repeats belonging to the classes of SR, LC and TTnotHomo under the definition of Low-Complexity Repeats (LCRs)^34^.

To further validate the robustness of our results, we repeated the analyses on a subset of RNA–RNA interactions with more stringent characteristics, such as longer interaction patches, that are more robust to mapping errors (**Supplementary Fig. 1h**, upper panel), or higher number of supporting reads (**Supplementary Fig. 1h**, lower panel), confirming the LCR enrichment in most of the sets. Moreover, we followed an orthogonal strategy by performing a permutation-based repeat enrichment analysis (see **Methods**), which estimates how much the observed overlap between interactions and repeats deviates from what would be expected by chance (**Supplementary Fig. 1i**). Overall, the results confirmed the strong enrichment of LCRs within the RNA–RNA interaction regions (**Supplementary Fig. 1j** and k), with even stronger enrichment observed when requiring the presence of repeats on both RNA interaction patches (**Supplementary Fig. 1j** and l). By analyzing the sequence of the LCRs involved in RNA–RNA interactions, we found that the majority of them were composed of multiple repeats of G- and C-containing units (G-rich and C-rich SR, G-rich and GA-rich LC, G-rich and CU-rich TTnotHomo, **Fig. 1c**, **Supplementary Fig. 1m** and **Supplementary Data 1**).

Finally, to obtain some potential insight on regulatory processes in which the interactions could be implicated, we performed Gene Ontology (GO) term enrichment analysis on RNAs found to interact through LCRs. Interestingly, most of these RNAs encode for proteins involved in developmental processes and that act as transcription factors (**Supplementary Fig. 1n** and **Supplementary Data 1**), even HeLa and HEK293T cells, which do not themselves undergo development or differentiation.

### LCR-bearing RRI patches are preferentially bound by proteins regulating stability and translation, and edited by ADAR

To gain additional insight into the functional role of LCRs in RRIs, we examined the overlap between RNA-binding protein (RBP) target sites, identified by the ENCODE consortium through eCLIP experiments in human cell lines^35^, and RRI patches detected by transcriptome-wide techniques. Across all examined human RRI sets, RRI patches were enriched for protein binding relative to matched controls (**Supplementary Fig. 2a** and b), with LCR-bearing patches consistently exhibiting higher overlap compared to those without LCRs. The co-occurrence between LCRs and RBP target sites within RRI patches prompted us to explore their functional interplay. First, we assessed which RBPs had a significant overlap of target sites with LCR-bearing RRIs and found a strong agreement between different RRI sets (**Supplementary Fig. 2c** and **Supplementary data 2**). As expected, the LCR-bearing RRI set derived from the RIC-seq experiment exhibited the highest number of enriched RBPs (**Supplementary Fig. 2d**). Among the proteins enriched in all the sets, we found three RNA helicases, UPF1, DDX6 and DDX55, consistent with the established role of this class of proteins in remodeling RNA duplexes. Of note, UPF1 is involved in nonsense-mediated and structure-mediated RNA decay^36,37^, while DDX6, a key component of P-bodies and modulator of stress granule (SG) formation, has been implicated in translational repression, RNA decay and miRNA-mediated gene silencing^38–42^. Furthermore, DDX6 also regulates A-to-I editing through its interaction with ADAR in the nucleus^43^, controlling neuronal differentiation. Then, we assessed which RBP functions^44^ were over-represented for each set of enriched RBPs. The results indicated that LCR-bearing RRIs tend to overlap with target sites of RBPs involved in translation regulation, viral RNA regulation (usually associated with RNA duplexes), RNA stability and decay, as well as cytoplasmic ribonucleoprotein granules (P-bodies and SG) formation (**Fig. 2a**, **Supplementary Fig. 2c** and **Supplementary data 2)**. Building on the hypothesis that LCR-mediated RRIs might influence RNA translation and stability, we examined their association with STAU1, a double-stranded RNA-binding protein known to play a role in these aspects of RNA metabolism—for example, by recruiting UPF1 to trigger mRNA decay^45,46^. To begin, we evaluated whether repetitive elements were enriched among the intermolecular RNA duplexes bound by STAU1, as detected in Flp-In T-REx 293 using the hiCLIP technique^47,48^ (**Supplementary data 2**). Nucleotide-level repeat enrichment analysis revealed a marked enrichment of LCRs among these duplexes (**Fig. 2b** and **Supplementary data 2**), supporting their potential role in modulating STAU1-dependent processes. To further explore this possibility, we re-analysed RNA-seq and ribosome profiling data produced from siRNA-resistant FLAG-STAU1 Flp-In 293 T-REx cells in three settings: UT (untransfected control), KD (siRNA-mediated STAU1 knockdown), and rescue (knockdown plus inducible siRNA-resistant STAU1 expression)^47^ (**Supplementary Fig. 2e**). Transcripts engaged in STAU1-bound heteroduplexes showed an overall increase in abundance in the KD versus UT comparison and a decreased abundance upon rescue, in contrast to those not involved in such interactions, consistent with the paradigm of STAU1-mediated decay (**Supplementary Fig. 2f,** left panel). However, while the overall magnitude of increase was independent of LCR presence in the RRI sites, the decrease in the rescue condition was stronger for transcripts with LCR-bearing STAU1-bound heteroduplexes (**Supplementary Fig. 2f,** left panel), suggesting an LCR-specific effect, possibly driven by the higher STAU1 expression observed in rescue compared to UT^47^ (**Supplementary Fig. 2e**). When examining translation efficiency, we observed an apparent overall increase upon KD specifically for the set of transcripts containing LCR-bearing STAU1-bound heteroduplexes, hinting at a potential role in STAU1-mediated translational regulation, although this trend did not reach statistical significance (**Supplementary Fig. 2f,** right panel). This effect, however, was not substantially reverted in the rescue condition (**Supplementary Fig. 2f,** right panel). Driven by our previous observation that transcripts involved in LCR-containing heteroduplexes are enriched for transcription factor-encoding ones, we investigated whether STAU1-mediated decay control acting on this class of duplexes might represent a mechanism to modulate transcriptional activity. Notably, 14% of the transcripts engaged in STAU1-bound, LCR-bearing heteroduplexes that were upregulated upon STAU1 KD were found to encode for transcription factors (**Supplementary Fig. 2g**). Among them, POU3F2 (**Supplementary Fig. 2h**), a transcriptional regulator of neural progenitor cells proliferation and differentiation that has been implicated in neuropsychiatric disorders^49^, forms a STAU1-bound heteroduplexes with a lncRNA (AL022324.4), with both interaction patches overlapping LCRs (**Supplementary Fig. 2i**). This lncRNA could transactivate STAU1-mediated mRNA decay, in partial agreement with the mechanism proposed by Gong and Maquat, which specifically involves Alu elements within the interaction patches^22^. Though detected in a non-neuronal context, this mechanism may hold functional relevance in neuronal lineages, particularly during development. Finally, since ADAR enzymes bind double stranded RNA and catalyze A-to-I editing^50^, we explored the potential of LCRs to form intermolecular RNA duplexes targeted by these enzymes. To this aim, we leveraged publicly available RNA-seq data from biological systems in which ADAR had been depleted, or its editing activity abrogated, to identify A-to-I editing events^51–53^. In every dataset, ADAR1-responsive adenosines showed stronger colocalization with RRI patches identified in the same system compared with unedited adenosines, and this enrichment was more pronounced for LCR-bearing RRI patches (**Fig. 2c**), highlighting a potential role for LCRs in guiding A-to-I modifications through the formation of intermolecular RNA duplexes. Interestingly, the connection with editing may not be limited to the A-to-I type, since we found, in all the LCR-bearing RRI datasets tested for colocalization with eCLIP-derived binding sites, a strong enrichment of sites bound by APOBEC3C (**Supplementary Fig. 2c**), a cytidine deaminase belonging to a family whose members have been shown to recognize stem-loop structures^54^.

**Fig. 2.**
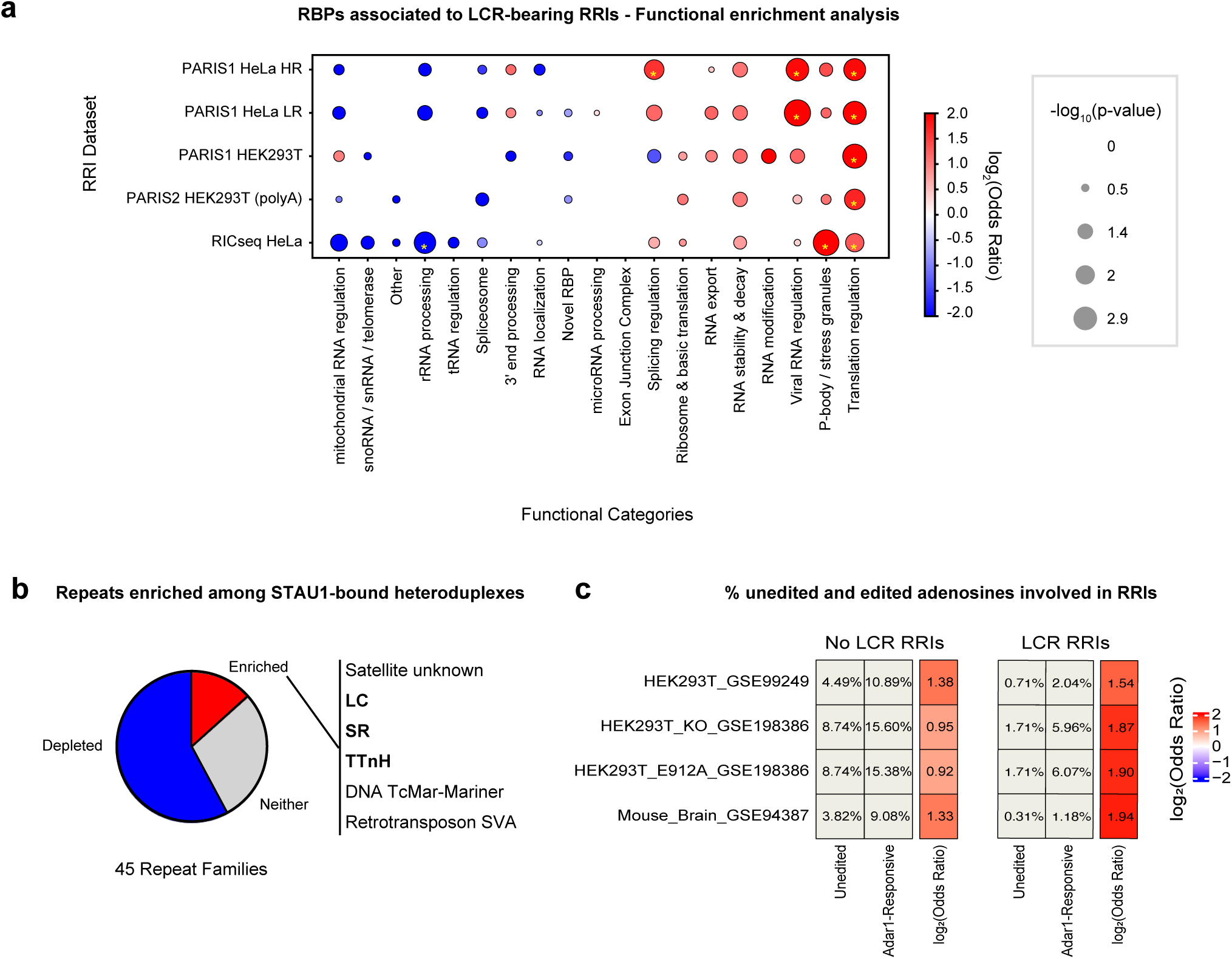
Translation, decay and editing factors recognize LCR-bearing RNA–RNA interaction sites. **a**, Bubble plot summarizing enrichment of RBP functional categories among RBPs associated with LCR-bearing RNA–RNA interactions across RRI transcriptome-wide experiments (y-axis). Columns report curated functional categories (x-axis). Color encodes the logC(Odds Ratio) (red = enrichment, blue = depletion), while bubble size reflects -log□□(p-value). Asterisks mark categories significant at p < 0.05 according to two-sided Fisher’s exact test. **b**, Pie chart showing the fraction of repeat families (out of 45 detected) that are significantly enriched (red), depleted (blue), or neither enriched nor depleted (gray) among STAU1-bound heteroduplex patches according to nucleotide-level enrichment analysis. The list on the right reports the enriched families, with LCR-related groups in bold (LC, SR, TTnH). Statistical significance was evaluated with two-sided Fisher’s exact test followed by Benjamini-Hochberg correction. **c**, Heatmaps showing the percentages of Unedited adenosines and ADAR1-responsive edited adenosines, identified across different biological systems (rows), that overlap with non-LCR-bearing (left) and LCR-bearing (right) RNA–RNA interaction patches detected in the same system. The rightmost columns display the log_2_(Odds Ratio) comparing the fractions of ADAR1-responsive versus Unedited adenosines involved in RRIs, with positive values indicating enrichment of RRIs among ADAR1-responsive sites. All differences are significant according to two-sided Fisher’s exact test with FDR correction, with the highest adjusted p-value equal to 0.0028.

### Low-complexity repeats are involved in multiple RNA**–**RNA contacts

We then assessed which mRNA regions are most involved in RNA–RNA interactions by performing a region-level RRI enrichment analysis (see **Methods**) on the transcriptome wide RNA– RNA interaction data. The results of this analysis, visualized *via* meta-transcript representation (**Fig. 3a**), revealed that, while interactions from almost all experiments are more prevalent in the 5’UTR, those identified in HeLa cells using PARIS1 are preferentially localized in the 3’UTR (**Fig. 3a**). Notably, LCRs over-representation in interactions showed a similar profile with respect to global RRIs enrichment, reinforcing the idea that the region-specific RRIs enrichment could be driven by low-complexity repeats (**Fig. 3a** and **Supplementary Fig. 3a**). Moreover, these results indicate that different biological systems could have different preferential regions for RNA–RNA interactions, albeit retaining the same properties. Indeed, the interaction sites at the 3’UTR in HeLa cells maintain the enrichment of low-complexity sequences (**Supplementary Fig. 3a**).

**Fig. 3.**
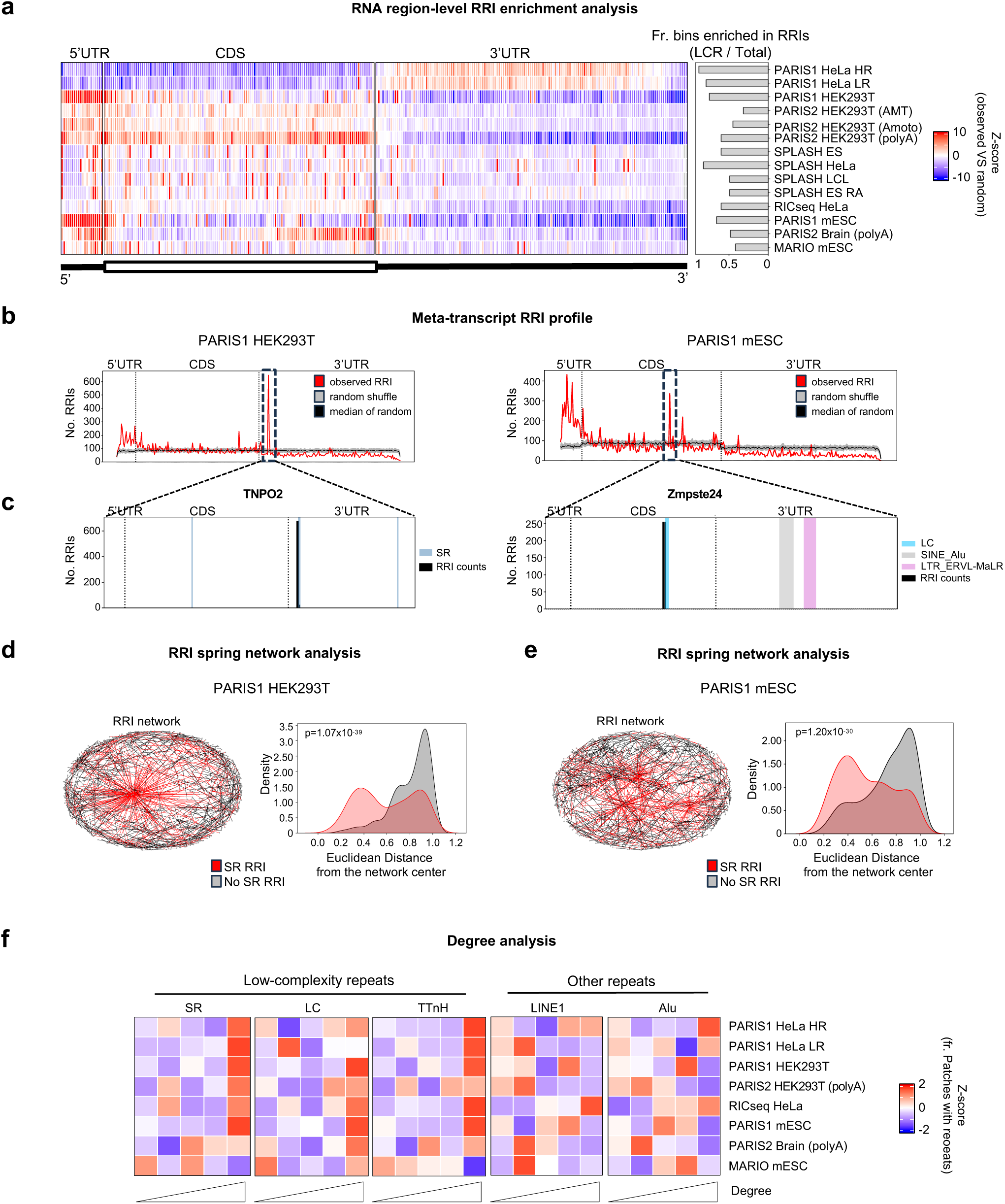
LCRs exhibit high connectivity in RNA–RNA interaction networks. **a**, Heatmap reporting, for each transcriptome-wide experiment (rows), the region-level RRI enrichment analysis Z-score calculated in each meta-transcript bin (columns). A representation of the coding and non-coding mRNA regions is depicted under the heatmap. The bar plot on the right reports the fraction of meta-transcript bins enriched in RRIs (Z-score > 2) that display LCR nucleotide enrichment in RRIs (FDR < 0.05, log_2_[OR]>0 for SR, LC or TTnH). **b**, Meta-transcript profile reporting, for PARIS1 HEK293T (left) and PARIS1 mESC (right) transcriptome-wide experiments, the amount of observed RRIs (red line) compared to the distribution of 100 random shuffles (gray area, with median value represented with a black line, region-level RRI enrichment analysis) across mRNA regions (x-axis). Dashed boxes highlight regions with the highest peak in CDS (PARIS1 HEK293T) or 3’UTR (PARIS1 mESC). **c**, Bar plot depicting the amount of RNA– RNA interactions identified in the binned sequence (x-axis) of TNPO2 (left) and Zmpste24 (right) transcripts, as well as the localization of repeated elements. **d** and **e**, Left panel: Spring network of RRIs involving LCRs (red edges) or not (black edges), detected in PARIS1 HEK293T (**d**) or PARIS1 mESC (**e**) experiments. Right panel: density plot reporting the distribution of nodes distance from the center of the network for RRIs overlapping (red) or not (gray) with LCRs. Statistical significance was assessed using two-tailed Mann-Whitney U-Test. **f**, Heatmap illustrating the row-scaled (Z-score) frequency of overlap between various repeat families and the RRIs detected by transcriptome-wide experiments with over 1000 interactions (rows). Frequencies are measured across five equally sized groups of RNA–RNA interaction patches (strata), ranked by their number of contacts (degree level).

Additionally, both the RIC-seq and the PARIS2 experiments performed upon polyA enrichment (HEK293T-polyA and mouse brain) revealed widespread enrichments in the CDS (**Fig. 3a**), suggesting some preference of these experiments in capturing contacts involving translated regions. Interestingly, we observed strong spikes in the meta-transcript profile of RNA–RNA contacts derived from PARIS1 experiments in HEK293T and mESC (**Fig. 3a** and **b**). We found that these spikes were due to single RNAs, TNPO2 for HEK293T and Zmpste24 for mESC, capable of generating large amounts of RNA–RNA interactions with multiple partners. Strikingly, we observed that 675 interactions of TNPO2 and 254 interactions of Zmpste24 were concentrated in specific regions of their transcripts, the first annotated as SR and the second as LC (**Fig. 3c**, left and right panel, respectively).

This finding prompted us to investigate whether LCRs have a natural tendency to form interactions with many different partners. To this aim, we performed a network analysis comparing one random sampling of repeats-associated RNA–RNA interactions to an equal number of interactions not associated with repeats. RRI network analysis could be robustly performed only in datasets with a suitable amount of RRIs, so, based on their size, we decided to focus on all the dataset with at least 1000 interactions. We utilized a spring network model that tends to localize nodes with more edges centrally within the network. The analysis demonstrated that LCR-containing interactions exhibit a shorter distance from the network center compared to control RRIs, confirming that they can engage more molecular partners (**Fig. 3d** and **e**). Notably, this is a common feature across the analyzed datasets. Indeed, we confirmed the same trend in all the datasets for at least one type of LCRs in at least 2 out of 5 independent samplings (**Supplementary Fig. 3b**). These findings held true also when using controls with matched expression to address expression-driven detection bias (**Supplementary Fig. 3c**). We further tested the potential of LCR-containing interacting patches to be involved in multiple interactions with an orthogonal approach. First, we stratified the RNA patches from each experiment into five bins based on the interaction degree. We then assessed the proportion of LCR-containing patches within each bin. In line with the previous result, we found that, for 7 out of 8 experiments tested, LCRs were enriched in the bins with highest degree (**Fig. 3f**). Interestingly, we did not observe the same behavior for other abundant repeats such as LINE1 or Alu/B1 SINE (**Fig. 3f**). To assess the interaction degree of RRI patches while controlling for RNA abundance, we focused on transcripts containing both LCR-bearing and non-LCR-bearing patches. For each transcript, we calculated the average degree of LCR-containing patches and of non-LCR patches (**Suppl. Fig. 3d**). We found that average degree values ≥ 2 were predominantly associated with LCR-bearing patches, confirming that these regions are frequently engaged in interactions with multiple RNA partners (**Suppl. Fig. 3e**).

### Complementary low-complexity repeats mediate lncRNA-target interactions

Looking for further evidence to support our findings, we explored the possible role of repeated elements in mediating the RNA–RNA interactions identified through targeted approaches. First, we focused on TINCR and its RNA interactome, revealed by Kretz and colleagues through RIA-Seq methodology^16^. The lncRNA hosts a UC-rich SR (**Fig. 4a**, upper panel). Therefore, we decided to investigate the repeat content of its top 300 targets, as compared to that of non-interacting RNAs with matched properties (**Supplementary Fig. 4a**). This analysis revealed that SRs and LC sequences are over-represented in TINCR interactors and, among them, the most abundant one is a GA-rich LC sequence, found in 12% of the targets, that could provide a complementary anchor for direct heteroduplex formation with TINCR’s UC-rich SR (**Fig. 4a** and **Supplementary Fig. 4b**). Since TINCR forms a complex with STAU1 that seems to stabilize target mRNAs, this would represent an alternative mechanism to the LCR-driven, STAU1-mediated RNA decay suggested by our previous results, highlighting how such interactions could function as regulatory modules whose effects are highly dependent on the context. Given the link between RNA–RNA interactions involving LCRs and regulatory genes implicated in development, we decided to experimentally investigate the RNA interactome of the lncRNA Lhx1os, recently described as a potential regulator of motor neuron (MN) development^55^. To this aim, we set up an RNA pull-down experiment, which allowed the isolation of the lncRNA Lhx1os using 20 nt long antisense biotinylated oligonucleotide probes in mESC-derived MNs. The experiment was performed employing two sets of probes, ODD and EVEN, targeting all the Lhx1os isoforms (**Fig. 4b**). A third set of probes, targeting the LacZ mRNA, was used as a negative control (LacZ) (**Supplementary Data 3**). The proper enrichment of Lhx1os was validated through qRT-PCR (**Supplementary Fig. 4c**). To identify the RNA interactors, we performed RNA sequencing of duplicated samples for each probe set, obtaining a set of 161 transcripts significantly enriched by ODD and EVEN, but not LacZ, purification (**Supplementary Fig. 4d** and **Supplementary Data 4**). We randomly selected 5 of them and validated their enrichment by qRT-PCR (**Supplementary Fig. 4e**). Analyzing the Lhx1os sequence with RepeatMasker, we observed the presence of a SINE B1 element (also annotated as MurSatRep1) in antisense orientation in the third exon of the transcript. We investigated the possible enrichment of complementary repeats in the identified RNA targets with respect to a control set with matched properties (**Supplementary Fig. 4f**). First, we focused on the SINE B1, wondering whether, among the candidates, there was an enrichment of complementary SINEs. Indeed, while the presence of SINE B1 in antisense orientation was comparable between target RNAs and controls, the SINE B1 with sense orientation, which could in principle pair with the antisense SINE B1 hosted by Lhx1os, was over-represented in the interacting RNAs (**Supplementary Fig. 4g**). Then, we carried out a global repeat enrichment analysis, revealing a significant over-representation of MurSatRep1 and GA-rich LC sequences within the targets (**Fig. 4c**). Since the GA-rich LC is the most frequently enriched repeat in the Lhx1os interactors (**Supplementary Fig. 4h**), we wondered if there was an association with the SINE B1 observed in Lhx1os. Interestingly, mapping the 7-mers complementary to the GA-rich LC on the Lhx1os sequence, we found that they are mainly located in the SINE B1 region, indicating a possible pairing of the two repeated elements (**Supplementary Fig. 4i**).

**Fig. 4.**
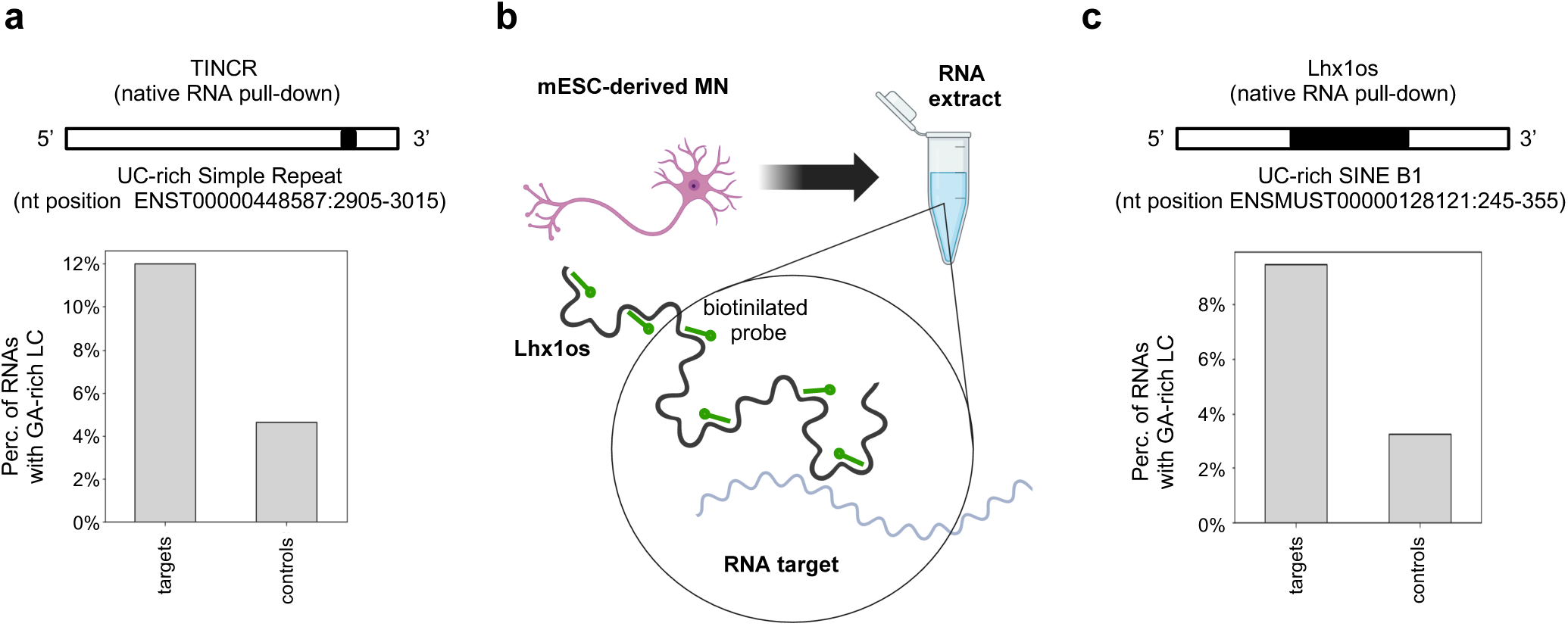
Transcripts targeted by TINCR and Lhx1os lncRNAs are enriched in LCRs. **a**, Top panel: schematic representation of the TINCR sequence, highlighting the position of the LC UC-rich element indicated. Lower panel: bar plot reporting the percentage of transcripts hosting LC GA-rich elements among the top TINCR interactors (targets) and negative controls. **b**, Schematic representation of Lhx1os RNA pull-down in mESC-derived motor-neurons (MN). Created with BioRender.com **c**, Top panel: schematic representation of the Lhx1os sequence, indicating the position of the UC-rich SINE B1 element. Lower panel: bar plot reporting the percentage of transcripts hosting LC GA-rich elements among the identified Lhx1os interactors (targets) and negative controls.

### Low-complexity repeats are involved in thermodynamically stable interactions

After determining the widespread presence of repetitive elements within the regions involved in RRIs, we proceeded to characterize the thermodynamic properties of these interactions, starting from those determined using transcriptome-wide approaches. To do this, we used the IntaRNA 2 software^25^ to predict the stability (ΔG) of each interaction. However, we noticed that, when using the default parameters, for roughly 10% of the interactions the software failed to identify any pairing in the evaluated regions (**Supplementary Fig. 5a**, left panel). This issue arose from the default seed constraint usage, which requires a perfectly complementary region of 5 to 8 nucleotides. By disabling the seed constraint, or by using IntaRNA 2 to emulate other prediction tools that do not require the presence of a seed (RNAup^56^ and RNAhybrid^57^), we were able to obtain ΔG scores for each interaction (**Supplementary Fig. 5a**, right panel, and **b**). This evidence suggests that a significant portion of the experimentally identified interactions do not rely on a seed as their core. Notably, these seedless interactions were found to be less thermodynamically stable compared to others (**Supplementary Fig. 5c**). Subsequently, we compared the ΔG distributions of experimentally identified interactions to a set of mock interactions derived from random RNA regions using multiple approaches (IntaRNA 2 with and without seed, RNAup, and RNAhybrid). While the deviation from the mock interactions was modest, all methods consistently showed that the experimentally identified interactions are more thermodynamically favorable than the random controls (**Supplementary Fig. 5d**).

Next, we focused on repeats. We stratified the interactions from each experiment into five groups, from the most thermodynamically favorable to the least favorable (**Supplementary Fig. 5e**), evaluating the enrichment of repeats in the various strata. Interestingly, LCRs showed a strong enrichment among the most thermodynamically favorable interactions, which was not observed for other repeated elements such as Alu or LINE1, used as controls (**Fig. 5a**). We then decomposed the interaction ΔG into two components: one attributable to intramolecular base pairing (reflecting the energy needed to make the two patches accessible) and the other to intermolecular base pairing (hybridization energy). Across most of the datasets analyzed, we observed that, although LCR-containing patches had higher predicted accessibility costs than patches lacking LCRs, they nevertheless engaged in more favorable intermolecular base pairing (**Supplementary Fig. 5f**). This suggests that, despite their stable local secondary structures, LCRs readily undergo rearrangement to form energetically advantageous intermolecular interactions.To validate the link between low ΔG values and LCRs, we examined the lncRNA–RNA interactions detected through targeted approaches. By performing an RNA contact enrichment analysis (see **Methods, Supplementary Fig. 5g**), we identified 5 TINCR regions which are more prone to interact with the target RNAs with respect to the control set for at least one of three algorithms (**Fig. 5b**, **Supplementary Fig. 5h** and **Supplementary Data 3**). Notably, 4 of the identified regions overlap with the TINCR motif boxes previously characterized as RNA-binding regions^16^. Of these identified regions, region 2 displayed the highest enrichment in contacts with the target RNAs, while region 4 exhibited the most favorable thermodynamics (**Fig. 5b**, **c** and **d**, and **Supplementary Fig. 5i**). First, we focused on region 2 and noticed that it does not coincide with any annotated repeat (**Fig. 5b**). However, by looking at the regions of the targets predicted to contact TINCR region 2, we observed an enrichment of SRs compared to controls, indicating that the predicted interactions with this region are probably mediated by LCRs present in the target RNAs (**Fig. 5c**). In contrast, analyzing region 4, we found that it coincides with the TINCR UC-rich SR (**Fig. 5b**) and we observed an enrichment in GA-rich LCRs in the contact region of the predicted targets (**Fig. 5d**), supporting the hypothesis that the interactions involving region 4 of TINCR are mediated by contacts between low-complexity sequences.

**Fig. 5.**
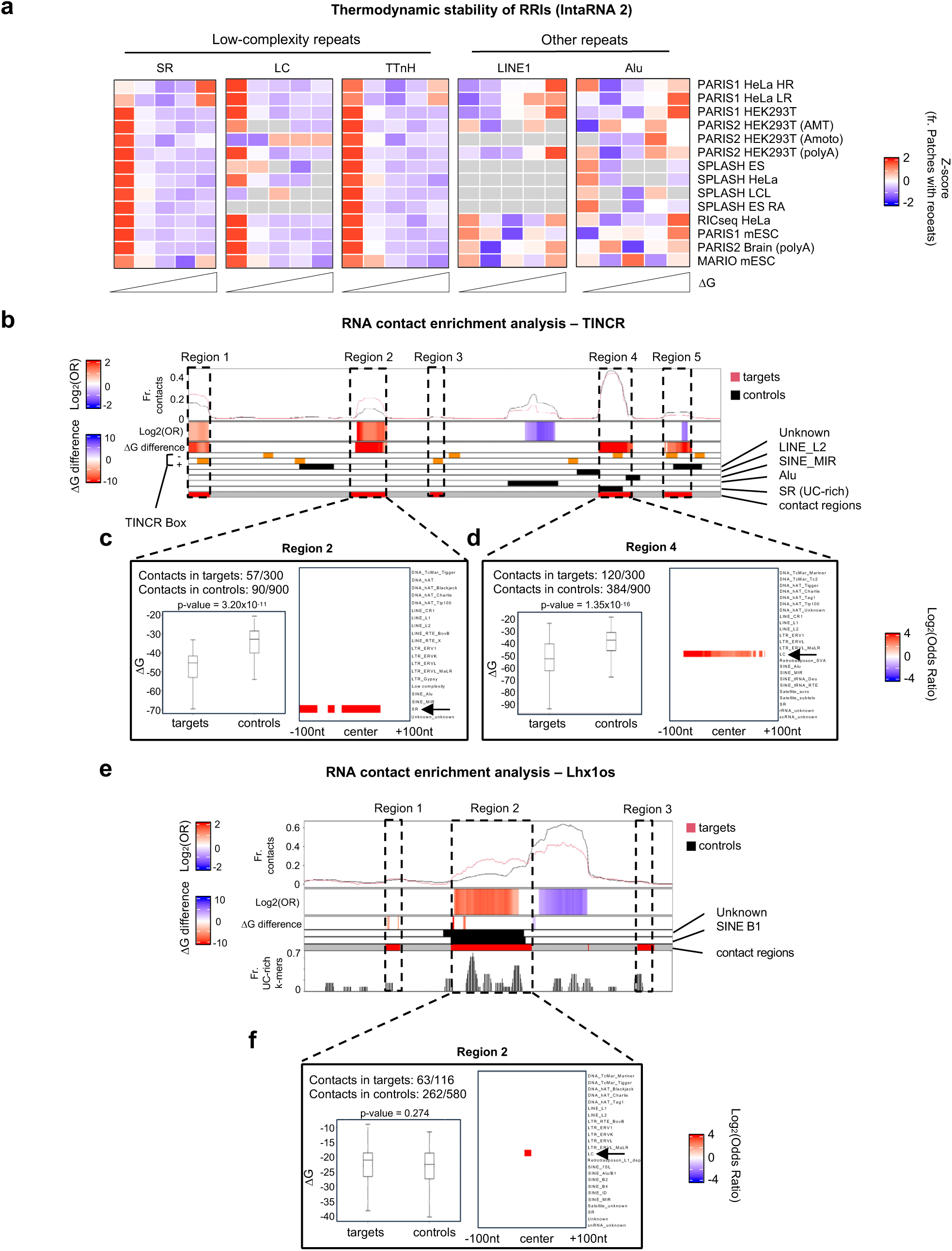
LCR-bearing RNA–RNA interactions exhibit thermodynamic stability. **a**, Heatmaps illustrating the row-scaled (Z-score) frequency of overlap between various repeat families and the RRIs detected by transcriptome-wide experiments (rows). Frequencies are measured across five equally sized groups of RNA–RNA interactions, ranked from left to right in order of decreasing thermodynamic stability. The ΔG prediction was performed using IntaRNA without seed constraint. **b**, Heatmap displaying the results of the RNA contact enrichment analysis for the TINCR RNA sequence. The sequence is represented from the 5’ end (left) to the 3’ end (right) in segments of 50 nucleotides, with a 10-nucleotide sliding step. The top panel line plot illustrates the frequency of RNAs from the target (red line) or the control (black line) group predicted by IntaRNA 2 software to be in contact with each segment. The first heatmap row displays log_2_(Odds Ratio) values, which measure the enrichment of contacts with target RNAs compared to control RNAs. Significant enrichment or depletion of contacts is indicated by red or blue colors, respectively; white denotes the absence of statistical significance. Statistical significance was assessed using two-sided Fisher’s exact test. The second row of the heatmap reports the difference between the median ΔG values of contacts with the target and control groups. Significant negative or positive difference of ΔG medians is indicated by red or blue colors, respectively; white denotes the absence of statistical significance. Statistical significance was assessed using two-tailed Mann-Whitney U-test. Orange boxes in the bottom annotation of the heatmap correspond to segments containing the TINCR box motifs in sense (+) or antisense (-) orientation. Black boxes correspond to bins with repeated elements from different families. Red boxes, also highlighted with dashed rectangles, indicate the 5 TINCR regions that displayed preferential binding with target RNAs according to at least one out of three prediction tools (IntaRNA 2, RIblast, ASSA). **c** and **d**, Visual representation of the results of RNA contact enrichment analysis performed on TINCR region 2 (**c**) and 4 (**d**). The text in the top-left area reports the amount of RNAs from the target and control groups that are predicted to contact the TINCR region. The box plot in the bottom-left area reports the ΔG distribution of predicted interactions between the TINCR region and the two transcript sets, calculated using IntaRNA 2. Boxes show the interquartile range (IQR), with the line indicating the median; whiskers extend to the most extreme data points within 1.5×IQR from the median; outliers were not included. Statistical significance was assessed using two-tailed Mann-Whitney U-test. The heatmap in the right panel presents the results of the meta-region analysis of repeat families (rows), with columns corresponding to nucleotide positions within a 200-nucleotide zone centered on regions from target or control RNAs that are predicted to contact the TINCR region. Significant enrichment or depletion of repeats in target RNA contact regions with respect to controls is indicated by red or blue colors, respectively; white denotes the absence of statistical significance. Statistical significance was assessed using two-sided Fisher’s exact test. **e**, Heatmap displaying the results of the RNA contact enrichment analysis for the Lhx1os RNA sequence. For a detailed explanation of the top rows, please refer to the **Fig. 5b** legend. The bottom row reports the frequency of UC-rich 7-mers found within the Lhx1os sequence. Dashed rectangles indicate the 3 Lhx1os regions that displayed preferential binding with target RNAs according to at least one out of three prediction tools (IntaRNA 2, RIblast, ASSA). **f**, Visual representation of the results of RNA contact enrichment analysis performed on Lhx1os region 2. For detailed explanations, refer to the legends of **Fig. 5c** and **d**.

Subsequently, we conducted an RNA contact enrichment analysis also on the Lhx1os pull-down dataset. As we hypothesized, thermodynamics-based predictions identified the SINE B1 as the region most prone to interact with the target RNAs, compared to the controls (**Fig. 5e**, **Supplementary Fig. 5l** and **Supplementary Data 4**). Additionally, the analysis of the predicted target regions contacted by Lhx1os SINE B1 confirmed the enrichment of the previously identified GA-rich LCR (**Fig. 5f**).

### Modelling RNA**–**RNA interactions with RIME, a deep learning approach based on language model embeddings

Our thermodynamic analysis of RNA–RNA interactions (**Supplementary Fig. 5d**) has revealed that, although the ΔG of experimentally detected duplexes is lower when compared to that of mock interactions generated with random RNA regions, it does not provide a neat separation between the two sets. Indeed, as already shown in other studies, thermodynamics-based computational tools often fail to predict biologically relevant interactions between long RNA molecules^26,58^. To handle the complexity of the interactions from a new perspective, beyond standard sequence representations like one-hot encoding^59–61^ and thermodynamic features, we exploited the Nucleotide Transformer (NT) LM^30^. This model provides dense vectorial representations of nucleotide sequences, known as embeddings, which efficiently capture underlying biological patterns. This LM proved particularly effective for training simple logistic regression models to detect the presence of LCRs, outperforming two well-known RNA LMs, RNA-FM^62^ and RNAErnie^63^ (**Suppleme**ntary Fig. 6a and b). Moreover, it enabled near-perfect discrimination between sequences with high and low predicted secondary structure stability (**Supplementary Fig. 6c**). Leveraging this advanced representation, we trained a deep neural network (RIME, RNA Interactions Model with Embeddings) to predict intermolecular RNA–RNA interactions, aiming to evaluate whether LMs can effectively model the determinants of RRIs (**Supplementary Fig. 6d**). The model was trained to infer whether two input RNA sequences, modeled using a contact matrix representation (see **Methods**), interact (positive class) or not (negative class). To ensure the identification of direct RNA–RNA interactions, independent of protein mediation, the positive set consisted of contact matrix windows encompassing duplexes detected by Psoralen-based methodologies (PP, Paired regions in interacting transcript Pairs, **Fig. 1a**, **6a** and **b**, see **Methods**). The negative set was constructed from three distinct classes of non-interacting windows (**Fig. 6a** and **b**, see **Methods**), sampled from the same RNAs used in the positive set to minimize abundance-related biases. These included randomly selected Non-paired regions in interacting transcript Pairs (NP), randomly selected Non-paired regions in Non-interacting transcript pairs (NN), and Paired regions already seen in the positive set combined in Non-interacting transcript pairs (PN). This setup allowed us to approach RRI prediction from two angles (**Fig. 6b** and **c**). Discriminating RNA–RNA Patches (DRP) focuses on distinguishing interacting regions (PP) from non-interacting ones (NP and NN). Discriminating RNA–RNA Interactors (DRI) involves identifying interacting regions within transcript pairs that are truly interacting (PP) versus those in randomly combined transcript pairs that are not interacting with each other (PN).

**Fig. 6.**
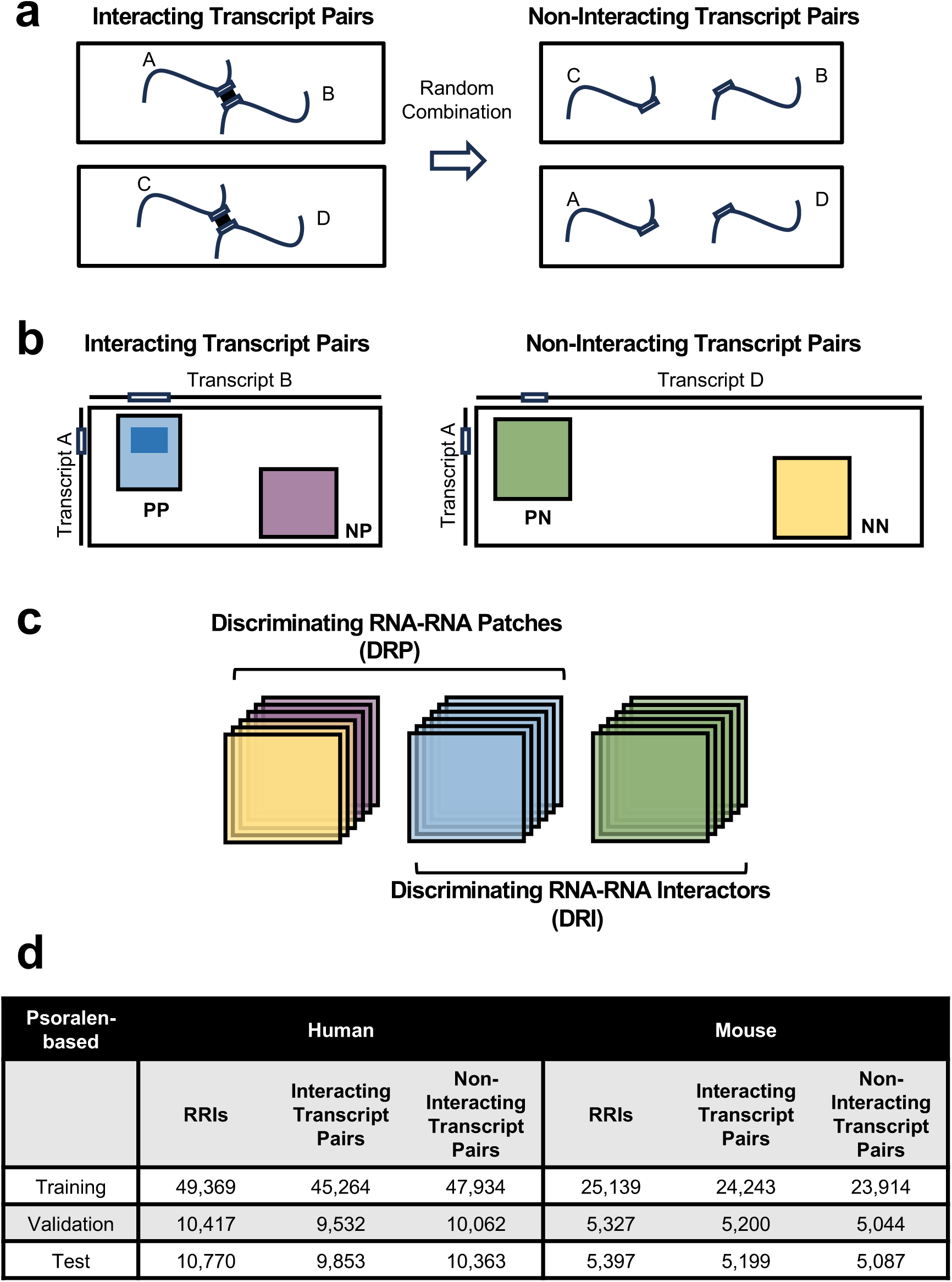
Preparation of datasets for training and evaluation of RIME. **a**, Schematic representation of the negative data generation process. Non-interacting pairs were generated by randomly combining interacting transcript pairs. The interacting sites are shown as white rectangles along transcript sequences, represented with lines. **b**, Illustration of Interacting Transcript Pairs (left) and Non-interacting Transcript Pairs (right), represented with contact matrices. Training and evaluation data were generated by sampling windows of varying sizes within these matrices, classified into four categories (PP, NP, NN, or PN) based on their positions. Interacting sites, depicted as white rectangles along the transcript sequence axes, are included in PP and PN windows but excluded from NN and NP windows. **c**, Visual representation of the DRP and DRI model evaluation tasks. In both cases, PPs serve as the positive set, while non-interacting regions (NN and NP) are used as the negative set for DRP, and randomly permuted interacting regions (PN) for DRI. **d**, Table reporting the amount of interacting region pairs (RRIs) and transcript pairs used to build the training, validation, and test sets for the Psoralen-based dataset.

The training set consisted of 70% of the total sample, while the remaining 30% was used for model evaluation (15% for validation and 15% for testing, **Fig. 6d**). To avoid potential biases we ensured similar RNA composition and length among the positive and negative set, as well as among the training, validation and test sets (**Supplementary Fig. 6e, f, g** and **h**), and verified that, overall, RNAs were sampled with comparable frequency in the positive and negative sets, preserving a balanced sampling also across the individual negative classes (NP, PN, NN) (**Supplementary Fig. 6i**). The F1 score on the validation set was monitored throughout training to guide model selection (**Supplementary Fig. 6j**). Furthermore, to verify the ability of the model to generalize beyond Psoralen-based methodologies, we evaluated its performance on protein-based datasets (RIC-seq and MARIO, **Supplementary Fig. 6k**).

### RIME enhances RNA**–**RNA interaction prediction beyond thermodynamic models

The extensive RRI set we compiled allowed us to perform a thorough evaluation of RIME performances, and to benchmark it against the most well-known thermodynamics-based prediction tools: IntaRNA 2^25^, ASSA^64^, RNAplex^65^, RNAhybrid^57^, RIsearch2^66^, pRIblast^67^, RNAup^68^, and RNAcofold^69^. We first assessed RIME’s prediction scores by comparing them with those from these tools. Thermodynamics-based programs clustered into two main groups, reflecting differences in how accessibility is considered into their predictions. Notably, RIME showed a weak to moderate correlation with thermodynamics-based tools, indicating that our method could capture novel interaction patterns (**Fig. 7a** and **Supplementary Fig. 7a**). To evaluate performance across different tools and account for errors in both positive and negative predictions, we used the Area Under the ROC Curve (AUC) as a global evaluation metric. This benchmark also included SPOT-RNA^70^, a deep learning-based RNA secondary structure prediction tool previously shown to be effective in predicting intermolecular contacts when concatenating the input sequences (SPOT-RNAc)^71^. The tools were tested on 200 nucleotide-long RNA regions pairs (**Supplementary Fig. 7b**). To evaluate the robustness of the tested methods, we assessed their performance on different subsets with progressively increasing quality, determined by the number of supporting reads (both for Psoralen-based and protein-based sets) and by the interacting regions length (only for the Psoralen-based methodologies, which rely on shorter chimeric reads).

**Fig. 7.**
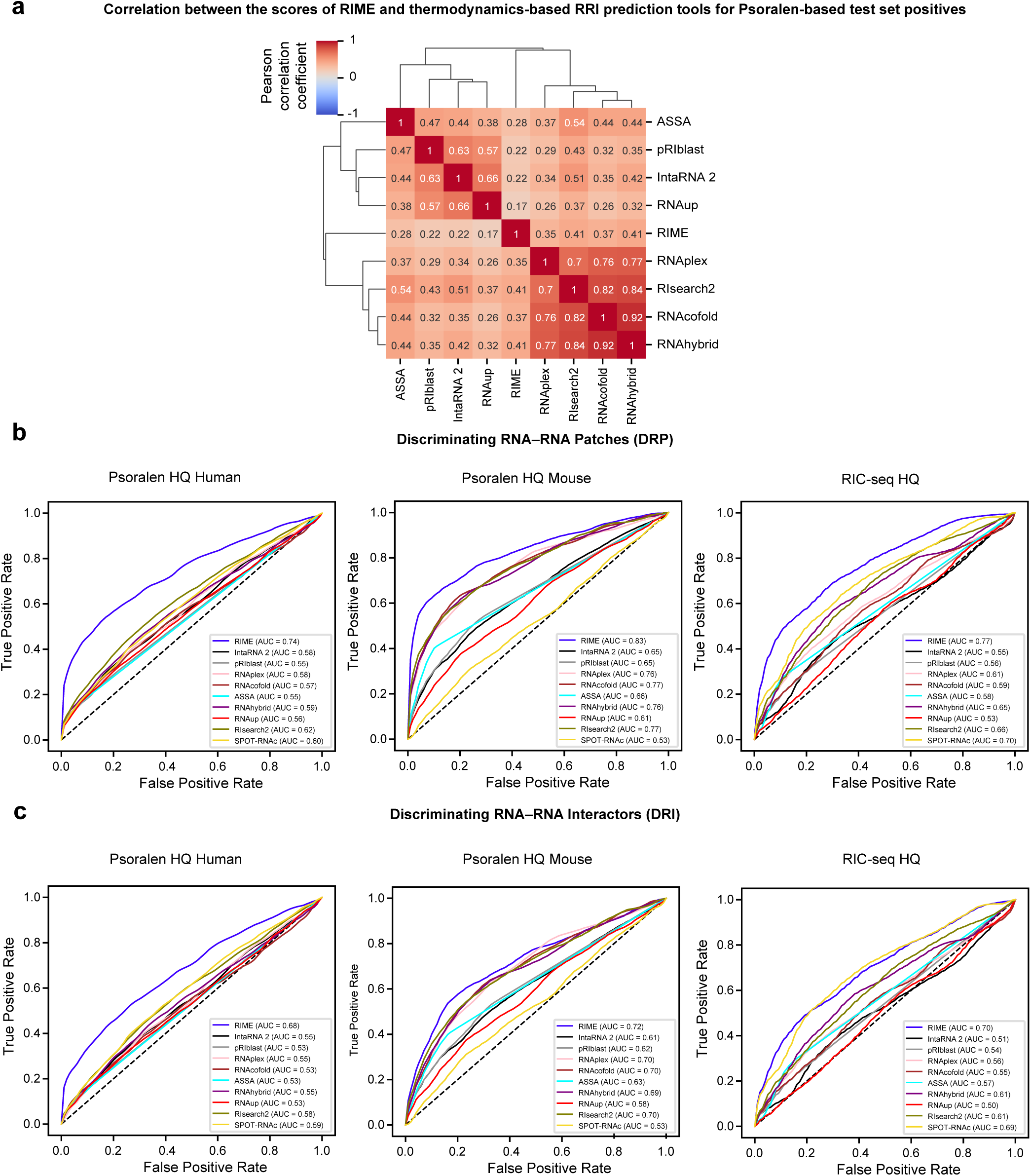
RIME overcomes the limitations of thermodynamics-based tools. **a**, Heatmap of Pearson correlation coefficients between RRI prediction tools, calculated using scores assigned to positive interactions in the Psoralen-based test set. For comparability with RIME scores, ΔG values from thermodynamics-based tools were converted by inverting their sign. Tools with similar scoring patterns were grouped using average linkage hierarchical clustering based on Euclidean distances. **b** and **c**, ROC curves describing the performances of the RRI prediction tools on the DRP (**b**) and DRI (**c**) tasks, shown for the Psoralen HQ Human (left), Psoralen HQ Mouse (middle), and RIC-seq HQ (right) datasets. To address class imbalance, negatives were randomly undersampled 100 times to match the number of positives, and ROC curves were averaged across iterations.

On the DRP task, RIME consistently outperformed all the other tools on human and murine Psoralen-based RNA–RNA interactions, especially when assessed against higher quality sets (**Supplementary Fig. 7c, d, e**, and **f,** left panel). In particular, for the human and murine subsets with at least 4 supporting reads or interacting regions length ≥ 35 nucleotides (hereby named Psoralen HQ Human and Psoralen HQ Mouse, **Supplementary Fig. 7g)**, RIME demonstrated an AUC of 0.74 and 0.83, respectively, while the top-performing thermodynamics-based tool (RIsearch2) reached 0.62 and 0.77, and SPOT-RNAc achieved 0.6 and 0.53 (**Fig. 7b**, left and middle panels, and **Supplementary Data 5**). Noteworthy, we observed a similar trend for the RIC-Seq dataset (**Supplementary Fig. 7h**, left panel), with RIME achieving an AUC of 0.77 when evaluated against interactions supported by at least 4 reads (RIC-seq HQ, **Fig. 7b**, right panel, and **Supplementary Fig. 7g**). In comparison, SPOT-RNAc, achieved an AUC of 0.70 on the same dataset, while thermodynamics-based tools reached a maximum AUC of 0.66 with RIsearch2. Although the overall performance metrics for the DRI task were comparatively lower, RIME still stood out as the top-performing tool, with its AUC improving alongside dataset quality (**Supplementary Fig. 7c, d, e**, and **f,** right panel). On the Psoralen HQ Human and Mouse datasets RIME demonstrated an AUC of 0.68 and 0.72, respectively, while the leading thermodynamics-based tool (RIsearch2) achieved an AUC of 0.58 and 0.70, and SPOT-RNAc achieved 0.59 and 0.53 (**Fig. 7c**, left and middle panels, and **Supplementary Data 5**). Consistently, on the RIC-seq HQ dataset, RIME’s AUC (0.7) was significantly better than RIsearch2’s one (0.61) and slightly better than SPOT-RNAc’s one (0.69, **Fig. 7c**, right panel, and **Supplementary Data 5**), which showed the best performance when evaluated on the full set (**Supplementary Fig. 7h**, right panel). On the MARIO dataset, consisting of murine protein-facilitated RRIs, the performance of the tools was generally lower (**Supplementary Data 5**). On the DRP task, RIME ranked second with 0.66, while the leading tool, RNACofold, achieved 0.68. For the DRI task, all tools performed at a level comparable to random chance, suggesting a limited ability of the methodology to capture specific interactions.

Finally, to evaluate the impact of our negative sample selection strategy on model’s performance, we generated RIME models trained on either the DRP or the DRI task, and evaluated their performance (**Supplementary Data 5**). When tested on the Psoralen-based sets, the models trained on the DRI tasks showed reduced performance on almost all sets, while the ones trained on the DRP task showed improved results on the DRP evaluation and a decrease in the DRI performance only for the HQ human set. However, the model trained jointly on both DRP and DRI tasks outperformed the ones trained on a single task when evaluated against the RIC-seq HQ and the MARIO datasets, underscoring the benefits of joint training strategy and the use of rigorously constructed negative sets in achieving better generalization to external data.

### RIME identifies key determinants of RNA interactions

The analysis of RIME’s performance prompted us to further explore the relationship between three features: data quality, which reflects the types of evidence supporting the reliability of experimentally identified interactions; RIME’s prediction score, indicating the model confidence in predicting positive or negative interactions; and RIME’s performances, representing the model overall predictive capability (**Fig. 8a**, left panel). At first, we focused on the relationship between data quality and RIME’s performances. As indicated by previous results, when selecting RRIs with gradually more stringent quality criteria, such as number of chimeric reads (**Supplementary Fig. 8a**) or interacting region size (**Supplementary Fig. 8b**), we observed that RIME’s performances increased monotonically. This result supports the idea that the model has understood the complex patterns behind true interactions and that the lower performance on the global dataset can be imputed to the higher proportion of false positives (**Figure 8a**, right panel). To further explore this aspect, we analyzed the connection between RIME’s prediction confidence and data quality. We found that the model’s confidence increased with data quality, (**Supplementary Fig. 8c** and d), indicating that the prediction score can be effectively used to prioritize interactions likely to be true positive (**Figure 8a**, right panel). Additionally, we analyzed the relationship between RIME’s performance and confidence for both positive and negative classifications, utilizing precision for positives and negative predictive value (NPV) for negatives (**Supplementary Fig. 8e**). We observed a positive correlation between the model’s confidence and its performance, highlighting the prediction score’s effectiveness in identifying reliable interactions (**Figure 8a**, right panel). We previously demonstrated that LCRs are crucial sequence determinants involved in RNA–RNA interactions. Additionally, we found that thermodynamics-based tools are highly effective at detecting interactions involving LCRs, as these typically exhibit low ΔG values across nearly all the analyzed RRI sets (**Supplementary Fig. 8f**). To assess whether also our deep learning model captured the significance of these sequence features, we analyzed the confidence scores for all the positive sets. Notably, we found that, consistent with other thermodynamics-based tools, RIME identifies LCRs as key determinants in RNA–RNA interactions, assigning higher scores to RRIs involving them, particularly when both interaction patches contain these elements (**Fig. 8b**). To gain deeper insight into the criteria driving RIME’s predictions, we employed Grad-CAM^72^, a gradient-based visualization technique that highlights the regions with the highest influence on the model’s outputs. Notably, Grad-CAM localization maps consistently revealed high-intensity regions near actual interacting sites, with this proximity improving as the maximum intensity in the localization map increased and the confidence score of the model improved (**Supplementary Fig. 8g**). This indicates that, especially for high-confidence predictions, it is possible to precisely pinpoint the regions involved in the interaction by looking at the areas most important for the model’s decisions (**Fig. 8c**). Moreover, for both Psoralen-based and RIC-seq datasets, the average distance between the highest-intensity regions and interaction sites was lower for RRIs involving LCRs than for those without LCRs, further highlighting that the convolutional layers capture biologically meaningful interaction features (**Supplementary Fig. 8h**).

**Fig. 8.**
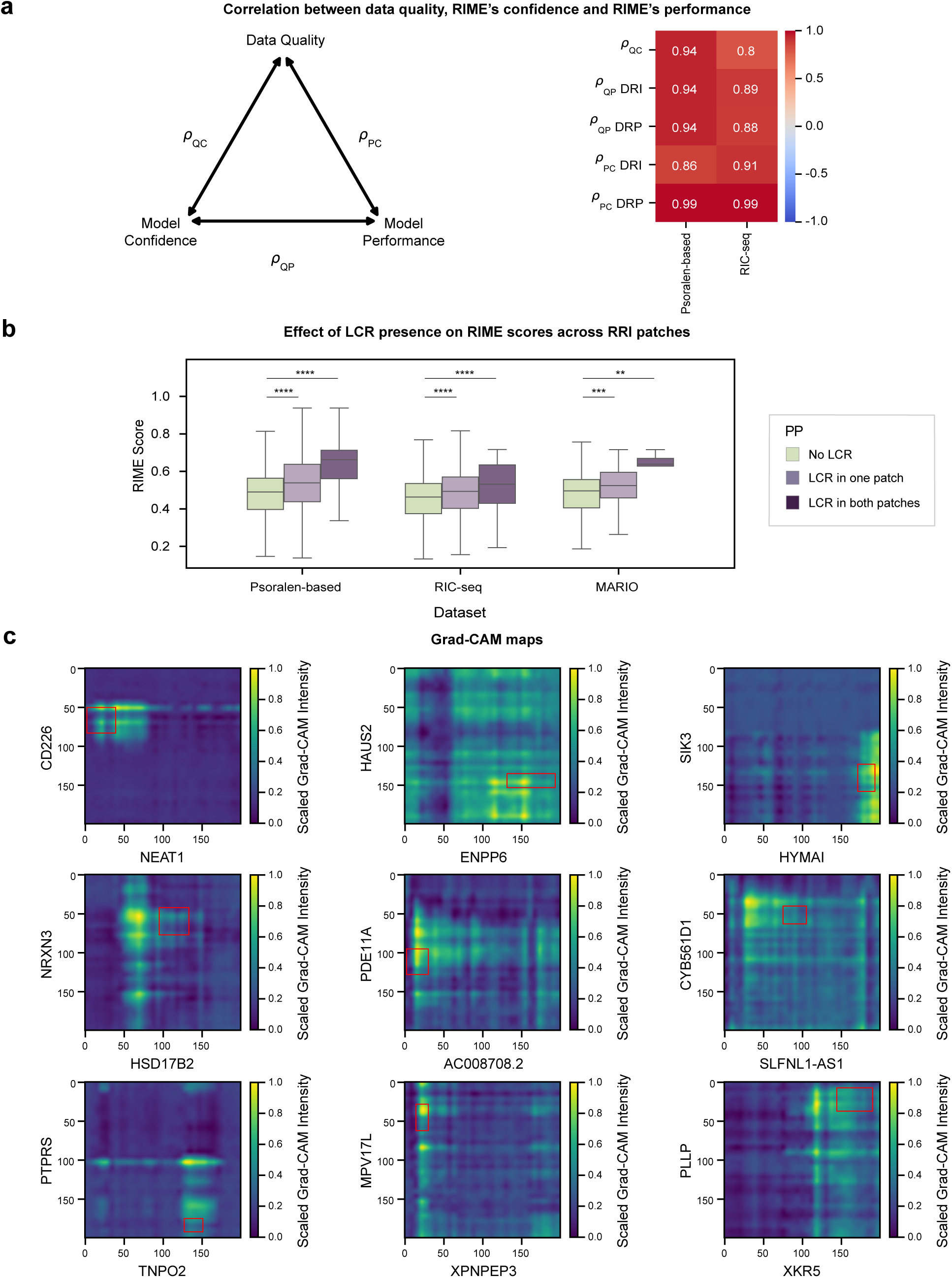
RIME captures critical features driving RNA–RNA interactions. **a**, Left panel: triangular diagram illustrating the relationships between data quality, RIME’s confidence, and RIME’s performance. Right panel: heatmap showing Pearson correlation coefficients between the three features, calculated for the Protein-based and RIC-seq datasets. p_QC_ is the correlation between data quality, represented by the minimum number of reads supporting an interaction, and model confidence, reflected by the mean RIME prediction score. p_QP_ was calculated for DRP and DRI tasks, between data quality (minimum number of reads supporting an interaction) and model performance (ROC-AUC), averaged over 100 iterations with negatives undersampled at various thresholds to match the positive set and quality criteria. p_PC_ is calculated for DRP and DRI tasks, measuring the Pearson correlation between model performance (calculated using the Precision metric) and model confidence (RIME score), which ranks predicted positive samples across progressively stricter confidence thresholds (100% of predicted positives, top 76%, top 52%, top 28%, top 5%). **b**, Box plots illustrating the distribution of RIME scores across the Psoralen-based, RIC-seq, and MARIO test sets. The scores are categorized based on the overlap of LCRs with the interacting patches: pairs where neither patch overlaps with LCRs, pairs where LCRs overlap with one patch, and pairs where both patches overlap with LCRs. Boxes show the interquartile range (IQR), with the line indicating the median; whiskers extend to the most extreme data points within 1.5×IQR from the median; outliers were not included. Two-tailed Mann-Whitney U test was used to test for differences between distributions. P-values were corrected with the Benjamini-Hochberg procedure. *, p < 0.05; **, p < 0.01; ***, p < 0.001; ****, p < 0.0001. **c**, Grad-CAM visualization of the top 9 (from left to right, from top to bottom) interactions from the Psoralen-based dataset. Interactions were ranked by averaging separate rankings based on the RIME score and the maximum Grad-CAM intensity value. Grad-CAM intensities were normalized between 0 and 1. The red bounding box highlights the true interacting regions.

### RIME accurately predicts high-confidence and functional interactions

To maximize data utilization and achieve the best possible model, we developed a final version of RIME, RIMEfull, by increasing the training data through the combination of the Psoralen-based training and validation datasets. To ensure the selection of the optimal model based on true interactions, we used a set of higher-quality interactions as the reference for model evaluation and selection (see **Methods**). Notably, this expanded training dataset enhanced the model’s performance when evaluated on the high-quality RIC-seq data (**Supplementary Fig. 9a** and **Supplementary Data 5**).

To further prove the reliability of RIMEfull’s predictions on external datasets, we tested its ability to identify RNA–RNA interactions involving the NORAD lncRNA, which were recently profiled using the COMRADES technique^73^. By reanalyzing the COMRADES data, we identified 17 mRNAs that specifically interact with the lncRNA (see **Methods**, **Supplementary Fig. 9b** and c). Mapping the interactions captured by the experiment onto the lncRNA sequence revealed regions more prone to contact the target RNAs (**Fig. 9a**, upper panel and **Supplementary Fig. 9d**). We used RIME to predict interacting regions between NORAD and each target RNA. Notably, we found that the top RIME predictions for each target clearly recapitulate the chimeric reads profile on the NORAD sequence, underscoring the ability of the model to recognize RNA–RNA interaction patches (**Fig. 9a**, lower panel). For instance, when examining EMP2, the NORAD interactor with the highest number of chimeric reads (**Supplementary Fig. 9e**), we observed a specific region near the 5’ end of the transcript that engages in multiple interactions with NORAD RNA (**Supplementary Fig. 9f,** left panel). Notably, RIMEfull successfully pinpointed this primary target region contacted by NORAD (**Supplementary Fig. 9f**, right panel). Furthermore, analysis of all 17 RNA–RNA pairs showed that the regions predicted by RIMEfull as interacting were significantly enriched in chimeric reads compared to those predicted as non-interacting (Fisher’s exact test p-value = 1.16×10C^16^, **Supplementary Fig. 9g**).

**Fig. 9.**
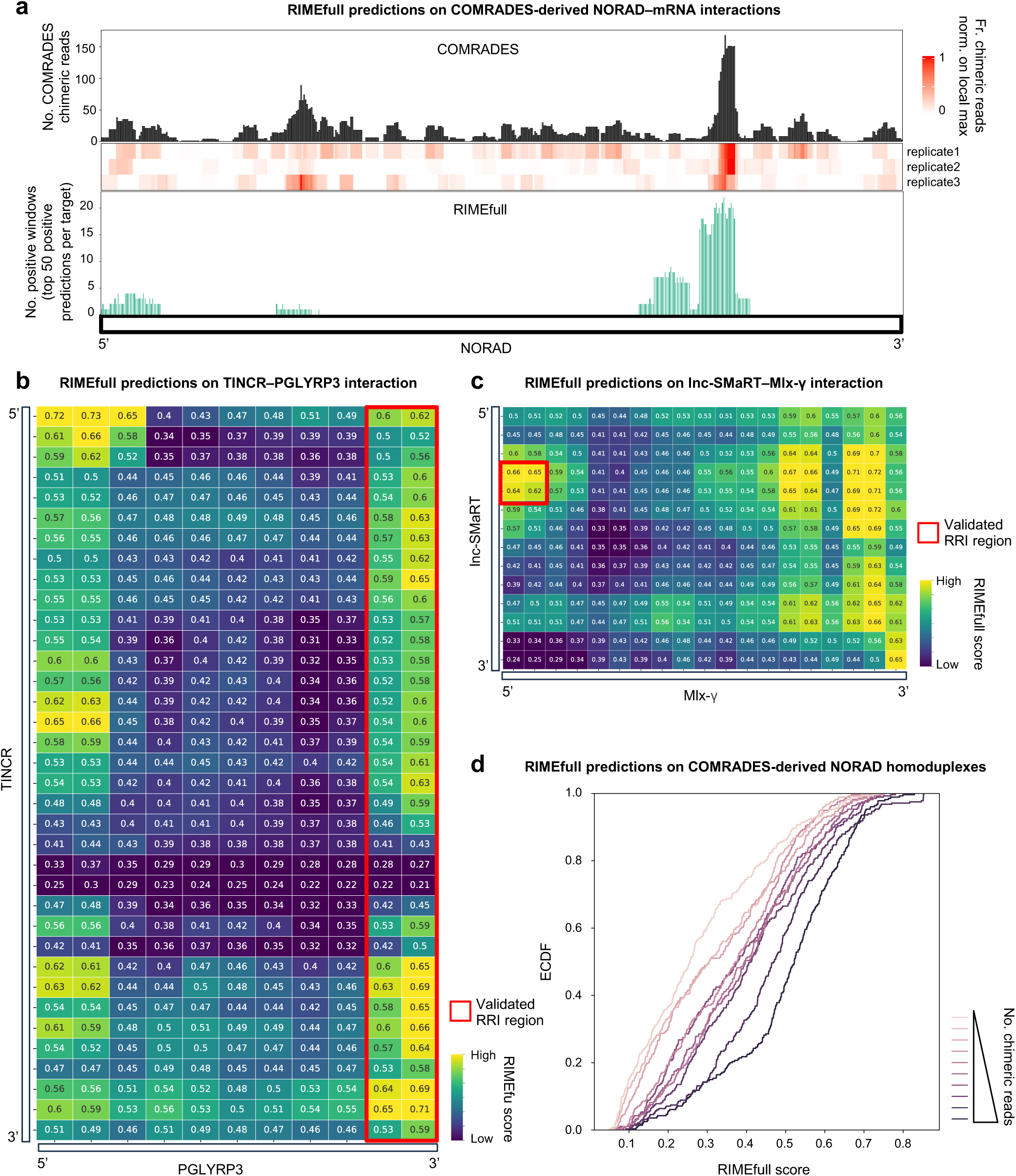
RIMEfull reliably identifies lncRNA–mRNA interactions. **a.** Plot summarizing the comparison between COMRADES-derived RRIs and RIME predictions for interactions between NORAD and its 17 mRNA partners. The NORAD sequence is represented from the 5′ end (left) to the 3′ end (right) in 50-nucleotide segments with a 10-nucleotide sliding step. The top bar plot reports, for each segment, the total number of COMRADES-derived chimeric reads connecting NORAD to its 17 target mRNAs across all experimental replicates. The central heatmap shows, for each replicate, the frequency of chimeric reads covering each segment, normalized on the maximum value observed across the segments. The bottom bar plot shows, for each segment, the frequency with which it was predicted as interacting by RIMEfull, considering the top 50 positive (prediction score > 0.5) windows for each target. **b** and **c**. Heatmaps showing the RIMEfull predictions for the TINCR–PGLYRP3 (**b**) and lnc-SMaRT–Mlx-γ (**c**) transcript pairs. Transcripts were segmented using a 200-nucleotide sliding window with a 100-nucleotide step. Each cell reports the RIMEfull prediction score for the interaction between the corresponding RNA windows. Previously validated interacting regions are highlighted with red boxes. **d**, Empirical cumulative distribution functions (ECDFs) of RIMEfull scores calculated for pairs of 200-nucleotide NORAD segments whose interaction is supported by varying numbers of chimeric reads. Each curve corresponds to one decile of the chimeric read count distribution, with darker colors indicating higher chimeric read support.

As a further validation, we evaluated the effectiveness of RIMEfull in predicting experimentally validated RNA–RNA interactions. As a first case study, we examined the interaction between TINCR and the PGLYRP3 mRNA, which depends on a region in its 3’ end for binding^16^.

Remarkably, RIME successfully identified this region as one of the most likely to interact with TINCR (**Fig. 9b**), underscoring the model’s ability to precisely identify RNA regions that mediate contacts with other RNAs. Next, we tested whether RIME could identify RNA–RNA interaction regions with known functional roles. To do this, we analyzed the interactions through which the murine lncRNA SMaRT negatively regulates the translation of Mlx-γ^74^ and Spire1^75^ mRNAs. In both cases, RIME efficiently recognized the experimentally validated interacting regions, while also identifying additional regions that were found to be putatively involved in the interaction between the lncRNA and its targets (**Fig. 9c** and **Supplementary Fig. 9h**). Notably, in the case of Mlx, these additional regions may explain why pull-down experiments captured Mlx isoforms lacking the primary interaction region (**Fig. 9c**).

Finally, we assessed the ability of RIMEfull to generalize to intramolecular interactions underlying RNA secondary structure. First,we divided the NORAD sequence into 200-nucleotide segments and computed RIMEfull scores for every possible segment pair (**Supplementary Fig. 9i**, left panel). Strikingly, the pattern of RIMEfull scores clearly resembled that of COMRADES chimeric reads supporting intramolecular interactions (**Supplementary Fig. 9i**, right panel). Moreover, when we stratified segment pairs by the number of chimeric reads, the distribution of RIMEfull scores progressively shifted toward higher values as the number of chimeric reads increased (**Fig. 9d**), while no trend was observed by repeating this analysis after shuffling the RIMEfull scores assigned to each pair (**Supplementary Fig. 9j**). These results indicate that RIMEfull is able to predict intramolecular duplexes and that its score correlates with confidence. This ability was further confirmed by the strong positive correlation observed between the RIMEfull scores obtained for intramolecular contacts within 100 randomly chosen transcripts and the average base pairing probability calculated using the RNAfold algorithm^23^ (**Supplementary Fig. 9k**), and by the negative correlation between RIMEfull scores and RNA base accessibility measured using DMS-MapSeq^76^ (**Supplementary Fig. 9l**).

## Discussion

In 1979, Britten and Davidson hypothesized a crucial role for repetitive sequences in forming RNA–RNA duplexes that could potentially regulate gene expression^77^. This early idea laid the foundation for current research recognizing repetitive RNAs as essential players in gene regulation^78^. Here, we conducted the first systematic investigation into the role of repetitive elements in mediating interactions between long RNA molecules, detected through transcriptome-wide and targeted approaches. Our analysis identified low-complexity repeats, such as tandem repeats, as the most prominent repeat families involved in RNA–RNA interactions. The functional relevance of LCRs was highlighted by their tendency to form interaction patches engaged in multiple contacts, as well as their significant enrichment in the RNA targets of long non-coding RNAs. These findings suggest that low-complexity repeats, by nature of their sequence composition, may act as “sticky” hubs within RNA interaction networks, promoting complementary interactions with multiple transcripts. Functionally, LCR-associated interactions frequently involve genes regulating developmental processes, consistent with a recent study highlighting connections between specific repeat subtypes and gene function^34^. LCR-bearing RRI patches also strongly overlap with target sites of RBPs, especially those controlling translation and decay, as exemplified by the significant representation of LCRs among the heteroduplexes recognized by STAU1, many of which involve mRNAs encoding transcription factors targeted by STAU1-mediated RNA decay. Among them, NR2F2, MNX1 and POU3F2 have a well-established role in neuronal development. For POU3F2 in particular, we identified a lncRNA that could potentially regulate it through LCR-mediated RRI. Extending this observation, we experimentally identified the RNA interactors of a lncRNA critical for motor neuron development and confirmed the presence of complementary LCRs that could be involved in contacts between the lncRNA and its targets. This reinforces the idea that LCRs play a role in coordinating the RNA interactome essential for cellular differentiation and tissue development. Finally, we show that heteroduplexes involving LCRs may act as preferential platforms for ADAR-mediated editing, underscoring their broader importance in post-transcriptional regulation and RNA metabolism. We hypothesize that low-complexity repeats evolved as central hubs for RNA interactions because their simple, repetitive sequences can arise readily through replication slippage or recombination, providing abundant and flexible modules for multivalent RNA–RNA and RNA-protein interactions.

On the pathological front, the association between LCRs and RNA–RNA interactions opens avenues for exploring disease mechanisms. Indeed, many repetitive elements are already linked to diseases, including Huntington’s disease^79^, autism^80^ and ALS^81^. Our findings raise the question of whether these diseases could also be investigated as “RNA–RNA interaction pathologies”. Notably, repeat-containing transcripts can phase-separate to form RNA *foci* within living cells, stabilized by intermolecular base-pairing interactions, which are frequently associated with disordered regions^82^. These structures may contribute to the long-term toxicity observed in neurodegenerative diseases^83–85^, underscoring the critical role of repeat-mediated RNA–RNA interactions in the pathology of these conditions. Our findings that LCR-bearing RRI patches often associate with proteins included in P-bodies and SGs further underscores their potential role in pathological granule formation.

Our *in silico* analysis revealed that LCRs act as sequence determinants that enable the formation of thermodynamically stable RNA–RNA interactions. However, while thermodynamic stability, as calculated by predictive tools, offers valuable insights, it is not always sufficient to accurately infer whether two RNA molecules interact *in vivo*. To address this, we developed RIME, a deep learning model that utilizes context-specific nucleotide sequence embeddings to capture the sequence determinants driving the interaction between long RNA molecules. The model was rigorously tested by assessing its ability to identify regions involved in RNA–RNA interactions and to distinguish, among potential pairings of these regions, those that form actual interacting duplexes. To our knowledge, this comprehensive evaluation approach has not been applied before. Moreover, we curated RNA–RNA interaction datasets for model training and evaluation, which are made available to the community to foster further advancements in the field. RIME demonstrated superior performance compared to thermodynamics-based models and to a deep-learning based tool. Notably, RIME excelled at predicting high-quality interactions, demonstrating its ability to extract meaningful patterns, such as the contribution of LCRs, from the noisy high-throughput experimental datasets used for its training. In fact, even if these sets are dominated by mRNA heteroduplexes, RIME successfully recapitulated the contacts formed between lncRNA regulators and their mRNA targets. It also showed good performance in predicting murine RNA–RNA interactions, despite the training set being largely composed of human RRIs. Moreover, despite not being explicitly trained to predict the exact positions of interacting regions, RIME showed a tendency to focus on areas in proximity to the actual contact sites to inform its decisions, which could be exploited to predict binding sites with higher resolution. Notably, RIME also proved effective at predicting intramolecular RNA contacts, a task for which it had no direct training, indicating that the intermolecular interactions features learned by RIME are also informative for intramolecular contacts. Despite these advances, RIME has some limitations. Although it generalizes well to RRIs detected by other experimental approaches, its training relied exclusively on Psoralen-based data, highlighting the need to incorporate additional datasets to mitigate potential biases associated with a single methodology. Moreover, because the current training set inevitably includes false positives, additional high-confidence RRIs will be essential to further improve accuracy. Finally, RIME relies on language model embeddings that are not RNA-specific, which, while powerful, could miss some important features captured by RNA-focused LMs. Nevertheless, given its superior performance and ability to capture the complexities of RNA interactions, RIME holds great potential for accelerating research in RNA biology, especially if integrated with gene expression data^86^, and may pave the way for novel therapeutic strategies targeting RNA-mediated regulatory networks.

## Methods

### Construction of an RRI dataset from transcriptome-wide studies

The genomic coordinates of PARIS1 and MARIO interactions were retrieved from the RISE database^87^. The coordinates of PARIS2 interactions were retrieved from the supplementary tables provided in the original publication^7^, and subsequently filtered to retain only canonical chromosomes. The RNA–RNA interactions coordinates from all these methodologies are referred to the hg38 genome assembly for experiments in *Homo sapiens* and to mm10 for those in *Mus musculus*. The interactions obtained *via* SPLASH experiments^8^, described by coordinates on the RefSeq human transcriptome^88^, were obtained from the resource available at the link https://csb5.github.io/splash/. Finally, the RIC-seq interactions^11^, provided by the authors upon request, were originally mapped to the hg19 genome assembly and subsequently converted to hg38 using the UCSC LiftOver tool^89^. Subsequently, the interactions of the RRI dataset annotated with genomic coordinates were mapped onto the reference human or mouse transcriptomes using the Ensembl 99 GTF annotation^89,90^. This mapping was performed using a custom Python script that runs the intersect module from BEDtools v2.29.1 suite^91^ with the parameters *-s-wao*. For genes with multiple isoforms, we employed the longest one, selecting the longest protein-coding variant in the case of protein-coding genes. Only RNA–RNA interactions where both patches were entirely mapped onto exons were retained, excluding interactions where both patches were mapped onto a single transcript, which likely correspond to intramolecular interactions. In cases where interaction patches mapped to ambiguous regions and were thus assigned to multiple genes, we selected a single representative gene using a specific prioritization strategy: genes with the most prevalent biotypes in the dataset and those with interactions in regions most involved in RNA–RNA duplex formation were given priority. Interactions involving small and/or abundant RNAs with the following biotypes were then filtered out: *Mt_rRNA*, *Mt_tRNA*, *misc_RNA*, rRNA, *rRNA_pseudogene*, *snRNA*, *snoRNA*, *scaRNA*, *miRNA*, and *scRNA*. Moreover, with the same intent, we filtered out interactions overlapping transcript regions annotated by RepeatMasker v4.1.1^32^ as belonging to the following repeat families: *SINE_tRNA*, *SINE_tRNA-Deu*, *SINE_tRNA-RTE*, *SINE_5S-Deu-L2*, *scRNA_unknown*, *rRNA_unknown*, *snRNA_unknown*, *tRNA_unknown*, *srpRNA_unknown*, *SINE_7SL*. Finally, due to the high amount of interactions in the RIC-seq RRI set (134,550, **Supplementary Data 1**), we further refined this set by selecting the interactions supported by at least two chimeric reads (9,808).

The biomaRt v2.48.3 R package^92^ was used to annotate RRIs, classifying them by the biotype of the interacting transcripts and the specific transcript regions involved.

### Detection of repeated elements

Interspersed repeats and low complexity sequences were found by scanning transcript sequences with RepeatMasker v4.1.1^32^ with default settings, specifying the proper organism (*Homo sapiens* or *Mus musculus*) and using rmblastn v2.11.0+ as internal engine. The RepeatMasker output was then converted to a BED file using the RM2Bed.py util of the software. Tandem repeats were also detected using Tantan v49 software^33^ with the *-f 4* parameter. The repeats identified by Tantan were divided in two groups: TanTanHomopolymeric (TTH), in which the repeating unit is composed by a single nucleotide and TanTan Not Homopolymeric (TTnH), in which the repeating unit is composed by multiple nucleotides.

### Repeat enrichment analyses

The nucleotide-level repeat enrichment analysis was performed as follows: for each repeat family, the BED file containing repeat coordinates was initially merged using the bioframe v0.2.0 Python library^93^, and the same was done for the BED file of the interaction patches from each RRI detection experiment. Subsequently, the two BED files were intersected using bioframe, and the number of nucleotides involved in interactions and belonging to the analyzed repeat family was calculated. The number of nucleotides specific to either the repeats or to the RRIs was computed by reciprocally applying the Bioframe *subtract* function. The number of nucleotides not involved in interactions and not belonging to repeats was determined by subtracting the sum of the nucleotides from the other three categories from the total length of the interacting transcripts. After that, a contingency table was populated with the four numbers (**Supplementary Fig. 1g**) and the statistical significance of the association between repeats and RNA patches was calculated using Fisher’s exact test with Benjamini-Hochberg correction for multiple testing. The analysis was also repeated by using only interaction patches with a minimum length or supported by a minimum number of chimeric reads. Length thresholds were independently determined for each experiment by dividing the patch length distribution into quintiles and using the resulting breakpoints (**Supplementary Fig. 1h**, upper panel).

For each experiment, the repeat enrichment profile along the meta-transcript was computed by applying the same procedure described above to each meta-transcript bin. Meta-transcript bins were generated as described in the following “Region-level RRI enrichment analysis” section.

Using BEDtools coverage, nucleotides within each bin were categorized into four groups: those exclusively involved in RRIs, those exclusively belonging to a repeat family, those belonging to both categories, and those belonging to neither. The abundance of each category was quantified, and a nucleotide-level repeat enrichment analysis was conducted for each bin. Finally, p-values were adjusted using the Bonferroni multiple testing correction.

The permutation-based repeat enrichment analysis was performed as follows: initially, the localization of repeat regions within the functional regions of RNAs (5’UTR, CDS, 3’UTR, or the whole transcript if non-coding) was determined using transcript information retrieved from Ensembl using BioMart^94^. Subsequently, 100 sets of mock repeats were generated by shuffling the repeat positions 100 times, while preserving the localization within the same functional regions (**Supplementary Fig. 1i**). For each experiment, BEDtools intersect was used to assess the overlap between RRIs and both real and mock repeats. The number of RRIs overlapping with real repeats was transformed to a Z-score by using the mean and the standard deviation of the distribution obtained by counting the overlaps of RRIs with the 100 shuffled mock repeat sets.

For each repeat family, the configurations X-R or R-X were assigned to interactions where the repeat overlap with only one of the two interacting RNA patches, while the R-R configuration was assigned to interactions where repeats of the same family are present in both the patches (**Supplementary Fig. 1j**). The overlap between repeats and patches was determined using BEDtools intersect.

### GO term enrichment analysis

Gene Ontology (GO) biological process term enrichment analyses of RNAs with SR or TTnH repeat-overlapping interaction patches were performed with the WebGestaltR v0.4.6 R package^95^, using all the interacting RNAs of the specified experiments as background. Only experiments with at least 5,000 interacting RNAs were taken into consideration.

### Analysis of co-occurrence between RRI sites and RBP target sites

To ensure robustness and to minimize noise derived from low interaction counts, the co-localization with eCLIP-derived RBP target sites analyses was assessed only for human RRI datasets containing more than 1,000 interactions. The selection of control regions for RRI patches was performed as follows. According to isoform annotations from Ensembl release 99, RRI patches mapped to mRNAs were assigned to their corresponding functional regions (5’UTR, CDS or 3’UTR), whereas those mapped to non-coding RNAs were labeled as “NC”. For each RRI patch, five control regions not involved in RRIs were randomly selected from transcripts detected within the same experiment, ensuring that each control belonged to the same region type as its corresponding RRI. This procedure was iteratively performed five times using BEDTools shuffle, specifying RRI patches as excluded regions (*-excl* parameter) and the coordinates of each RNA region as inclusion intervals (*-i* parameter). The entire transcript coordinate range (from 0 to the transcript length) was provided as the genome file (*-g* parameter). For non-coding RNAs, the full transcript sequence was directly used as the sampling universe. After control generation, both RRI and control patches were merged using BEDTools merge to avoid redundant evaluation of overlapping regions. The “LCR” or “No LCR” labels for RRI and control patches were then assigned using BEDTools intersect and the LCR annotation described in the “Detection of repeated elements” section. Finally, transcript coordinates of RRI patches and control regions were mapped to genomic coordinates (GRCh38).

RBP target regions derived from eCLIP experiments on human cell lines^35^ were retrieved by downloading IDR peaks from the ENCODE^96^ data portal (available at https://www.encodeproject.org/) using the accession IDs reported in the work by Van Nostrand and coworkers^44^. All IDR peaks corresponding to the same binding protein but originating from different experiments were merged.

Overlaps between RRI patches (or their controls) and RBP interaction sites were computed using BEDTools intersect with the *-s* parameter to ensure strand specificity. Only genes detected in the RRI detection experiments and showing a mean expression level greater than 1 Fragments Per Kilobase per Million mapped reads (FPKM) in the HepG2 and K562 systems were considered. Expression values were estimated from three random samples of the SCR control datasets generated by Van Nostrand et al.^44^. Finally, based on the overlap results, merged RRI patches or control regions were classified as either RBP-bound or unbound, and the associated RBPs were annotated accordingly.

For each RBP, we assessed whether its target sites preferentially colocalized with LCR-bearing RRI patches compared to their control regions using Fisher’s exact test followed by Bonferroni correction for multiple testing. This analysis was performed independently for each experiment. RBPs with corrected p-value < 0.05 and log_2_(Odds Ratio) > 0 were classified as enriched. Functional enrichment analysis of enriched RBPs was then performed independently for each experiment using the RBP functional annotations from Van Nostrand et al.^44^. For each functional category, enrichment significance was assessed using Fisher’s exact test, comparing the proportion of RBPs associated with that function between the enriched and the non-enriched ones. Given the limited number of features, only raw p-values were considered, without correction for multiple testing.

### Analysis of STAU1-bound intermolecular duplexes

STAU1-bound intermolecular RNA duplexes identified from STAU1 hiCLIP data using Tosca software were kindly provided by Chakrabarti and colleagues^48^. The dataset was filtered following the criteria described in the “Construction of an RRI dataset from transcriptome-wide studies” section. Enrichment of repeated elements within the interacting regions was then assessed by performing a nucleotide-level repeat enrichment analysis.

Raw reads produced from Flp-In T-REx 293 cells datasets under control (UT), Stau1 knockdown (KD) and rescue conditions were downloaded from the European Nucleotide Archive (ENA) using the following accessions: ERR605044, ERR605045, and ERR605046 for the Ribo-seq samples, and ERR618765, ERR618766, and ERR618767 for the corresponding RNA-seq libraries (Input). Quality control of reads was assessed using FASTQC software v0.11.9 (available at https://www.bioinformatics.babraham.ac.uk/projects/fastqc/). Demultiplexing and unique molecular identifier (UMI) extraction were performed jointly with Ultraplex v1.2.10^97^ using the NNNCGGANN and NNNTGGCNN barcodes to distinguish the first and second replicate, respectively, for each condition. Adapters and technical artefacts were removed with Trim Galore v0.6.10 (https://github.com/FelixKrueger/TrimGalore) using the parameters *--trim-n --length 20 -- 2colour 20*, with Cutadapt v1.18^98^ as the trimming engine. Reads were aligned to the GRCh38 reference genome with STAR v2.7.11b^99^, using gene annotations from Ensembl release 99 in the STAR index and the following parameters: *--winAnchorMultimapNmax 100 -- seedSearchStartLmax 20 --outFilterMultimapNmax 1 --outFilterMismatchNmax 3 --alignEndsType EndToEnd --readFilesCommand zcat --outSAMtype BAM SortedByCoordinate*. PCR duplicates were subsequently removed with UMI-tools v1.1.1^100^ using *dedup --extract-umi-method=read_id -- umi-separator=’rbc:’ --method unique*. Gene-level quantification was performed with featureCounts (Subread v2.1.1)^101^ using *-g gene_id-s 1*; the *-t* feature type was set to *exon* for expression analyses and to *CDS* for coding-sequence quantification used in translation efficiency estimates. The differential expression analysis was performed using the edgeR v3.34.1 R package ^102^ as follows. Genes not reaching a minimum of 10 counts in at least two replicates were excluded from subsequent analyses. Count data were upper-quartile normalized prior to statistical modeling, while FPKM values were calculated using the length of the exon/CDS union per gene according to the Ensembl GTF annotation (release 99). Dispersion estimates were obtained using estimateDisp function with *robust=TRUE* parameter, and model fitting was performed with glmFit. Contrasts were defined using makeContrasts to compute log_2_(fold changes) (log_2_FC) between conditions, specifically KD vs UT or rescue vs KD for gene expression comparisons. Significantly deregulated genes were defined as those with Benjamini–Hochberg adjusted p-value < 0.05.

For translation efficiency analysis, interaction contrasts of the form (Ribo-seq_KD - Input_KD) - (Ribo-seq_UT - Input_UT) and (Ribo-seq_rescue - Input_rescue) - (Ribo-seq_KD - Input_KD) were applied to estimate the Delta log_2_FC values reflecting differential translation efficiency between conditions. To retain only expressed genes, the results were further filtered retaining only genes with average FPKM > 1 in Input samples according to exonic features quantifications.

To detect transcription factors potentially regulated *via* LCR-driven, STAU1-mediated RNA decay, we selected the genes significantly up-regulated upon STAU1 KD and involved in LCR-bearing, STAU1-bound heteroduplexes. Biological Process, Molecular Function, and Cellular Component GO terms associated to these genes were retrieved from Ensembl 99 *via* BioMart, and genes whose GO terms explicitly described transcription factor activity were identified (GO:0005667, GO:0000981, GO:0003700, GO:0005669, and GO:0005673).

### Analysis of A-to-I editing in RNA–RNA interaction sites

To assess the relationship between A-to-I editing and intermolecular RNA duplexes, we performed a genome-wide analysis of ADAR-responsive editing events in three biological systems: human HEK293T cell and mouse brain. For HEK293T cells, we analyzed two publicly available RNA-seq datasets, retrieved from Gene Expression Omnibus (GEO) database^103^ The first dataset was produced by Chung et al.^52^, who used CRISPR/Cas9 to generate ADAR1 knockout HEK293T lines (GEO accession GSE99249, sample IDs: SRR5564268, SRR5564272, SRR5564273, SRR5564274, SRR5564275, SRR5564276). The second was generated by Hu et al.^53^, who profiled HEK293T cells lacking ADAR1, as well as cells expressing catalytically inactive ADAR1 mutants bearing the E912A substitution (GEO accession GSE198386, sample IDs: GSM5946011, GSM5946010, GSM5946009, GSM5946008, GSM5946007, GSM5946006). For the mouse brain, we used the RNA-seq reads produced by Heraud-Farlow and colleagues^51^, who generated ADAR1 editing– deficient mice by introducing a homozygous E861A point mutation in the Adar1 gene, abolishing the enzyme’s RNA editing activity (GEO accession GSE94387, sample accessions SRR5223114, SRR5223115, SRR5223116, SRR5223117, SRR5223118, and SRR5223119); to rescue the otherwise lethal phenotype, these animals were crossed onto an Ifih1-deficient background (Adar1^E861A/E861A^Ifih1-/-).

Reads were first trimmed using Trim Galore v0.6.10 (https://www.bioinformatics.babraham.ac.uk/projects/trim_galore) to remove adapter sequences and low-quality bases with parameters *--2colour 20 --trim-n* and a minimum read length after trimming of 18, and the resulting reads were aligned to the reference genome using STAR v2.7.7a^97^ with alignment options *--readFilesCommand zcat --outSAMtype BAM Unsorted --outFilterType BySJout --outFilterMultimapNmax 20 --alignSJoverhangMin 8 --alignSJDBoverhangMin 1 -- outFilterMismatchNmax 999 --outFilterMismatchNoverReadLmax 0.04 --alignIntronMin 20 -- alignIntronMax 1000000 --alignMatesGapMax 1000000 --peOverlapNbasesMin 15 --quantMode GeneCounts --chimOutType Junctions*. Multimapping reads (NH < 2) were filtered out using BamTools v2.5.1^98^, and BAM files were indexed and annotated with MD tags using SAMtools v1.7^99^. ADAR-mediated RNA editing events were then identified with JACUSA2 (v2.0.4)^100^ in *call-1* mode, with parameters *-c 1 -f B -P RF-FIRSTSTRAND*, filtering background noise and accounting for strand specificity. The reference genome used was GRCm38 for mouse and GRCh38 for human samples. Base call counts were aggregated across replicates of the same condition, and only reference adenosines with a minimum coverage of 15 after pooling were retained for downstream analyses. For each RNA-seq experiment, BEDtools intersect with the *-s* option was used to select the adenines on the reference belonging to the exons of transcripts involved in RRIs identified in the same biological system. These adenines were classified based on the proportion of observed A-to-G mismatches. Positions with no A-to-G conversion in WT were labeled as “Unedited”. To identify ADAR1-responsive editing sites, adenines with at least one A-to-G substitution in WT were selected, and a one-sided Fisher’s exact test was applied to assess whether the G/A ratio significantly decreased in the ADAR1 loss-of-function (LOF) condition compared to the wild type (WT). Editing sites with a p-value < 0.05 and a log_2_(Odds Ratio) > 1 (supporting decreased editing in LOF) were classified as “ADAR1-responsive”. Using BEDtools intersect with the *-s* option, we further classified Unedited and ADAR1-responsive adenines according to their overlap with RRIs detected in the same biological system, distinguishing between RRIs with or without LCRs. For each RRI category, the proportion of Unedited adenines overlapping with RRIs was compared to that of ADAR1-responsive adenines using two-sided Fisher’s exact tests, followed by Benjamini-Hochberg correction for multiple comparisons.

### Region-level RRI enrichment analysis

The functional structure of interacting mRNAs, described by the lengths of the 5’UTR, CDS, and 3’UTR regions, was summarized into a meta-transcript representation. Region lengths, together with the transcript sequences, were retrieved from the Ensembl 99 database using the biomaRt v2.48.3 R package^92^. mRNA regions within the same class (5’UTR, CDS, or 3’UTR) were divided into an equal number of bins, depending on the ratio between the median length of that class and the median length of the other classes: 20 for the 5’UTR, 130 for the CDS and 150 for the 3’UTR. The bin ranges for each RNA were compiled into a BED file and, for each experiment, the amount of RRIs in meta-transcript bins was calculated using BEDtools intersect. Mock interaction sets were generated using BEDtools shuffle with the parameters *-chromFirst -bedpe.* A random distribution was obtained by counting the overlaps of meta-transcript bins for each of the 100 mock interaction sets, and its mean and standard deviation were used to calculate the Z-score for real RRIs in each bin. Graphical representations of the meta-transcript analyses were generated using the ComplexHeatmap v2.8.0 R package^104^.

### RRI network analysis

RRI network analysis was performed for each transcriptome-wide experiment using the NetworkX v2.5.1 Python package^105^. Repeat family-specific networks were constructed by combining RRIs that overlap with repeats from a specific family with an equal number of randomly selected RRIs that do not overlap with repeats from that family. If the number of repeat-overlapping RRIs exceeded 500, a random subset of 500 was selected to build the network, in which each node represents a transcript and the edges represent the interactions. The weights of the spring network were defined by the number of chimeric reads that supported each interaction. Nodes were positioned using the Fruchterman-Reingold force-directed algorithm provided by the *spring_layout* function. Then, for each node, the euclidean distance from the center (coordinates 0, 0) of the spring network was calculated by applying the *distance.euclidean* function from scipy.spatial Python module. For each edge, the distance from the network center was calculated as the average of the distances of the two nodes involved in the interaction.

To integrate gene expression into the network analyses, we used fastq-dump from SRA Toolkit v2.10.9 to retrieve bulk RNA-seq data from the GEO database corresponding to the biological systems profiled for RRIs: HEK293T (GSE99249: SRR5564274, SRR5564275, SRR5564276)^52^, mouse brain (GSE94387: SRR5223117, SRR5223118, SRR5223119)^51^, HeLa (GSE217878: GSM7470721, GSM7470724, GSM7470718, GSM7470715)^106^, and mESC (GSE160578: GSM4875577, GSM4875578, GSM4875579)^107^. Processing followed the workflow detailed in the “Analysis of A-to-I editing in RNA–RNA interaction sites*”* section. Gene expression was summarized as FPKM and averaged across replicates; gene length for FPKM normalization was computed as the number of nucleotides in the union of exonic intervals from Ensembl release 99. The selection of the RRIs for the network analysis proceeded as follows. First, for each RRI, an expression metric was defined as the mean of the two genes’ FPKM values.Then, for each LCR type, we selected a subsample of 100 repeat-overlapping RRIs and, for every repeat-overlapping RRI, one control interaction was chosen from the pool of non-repeat-overlapping RRIs by ranking candidates on the absolute difference in expression metric (|Δexpression|) and retaining the one with the smallest |Δexpression|. The subsample size was reduced from 500 to 100 to mitigate residual imbalances in the expression metric between LCR-bearing RRIs and their controls. The comparability of expression metrics between repeat-overlapping RRIs and their matched controls was verified with a two-sided Mann-Whitney U test, and only comparisons with pC>C0.05 were retained.

The interaction degree was calculated as follows. Non-redundant interacting RNA patches were identified by collapsing all the overlapping RNA regions involved in RRI across each experimentally-derived set using BEDtools merge, considering the strand of the feature (**Supplementary Data 1**). Next, BEDtools intersect was used to map RRI patches to RNA regions, enabling the counting of unique RRI identifiers assigned to each RNA patch. BEDtools intersect was also applied to identify RNA patches overlapping with repetitive elements.

To analyze the interaction degree while controlling for abundance, we selected from each RRI dataset transcripts with at least one LCR-bearing RRI patch and at least one non-LCR-bearing patch. Then, for each of these transcripts we calculated the mean interaction degree of the LCR-bearing patches (LCR meta-patch) and of the non-LCR-bearing patches (No LCR meta-patch), resulting in two sets of meta-patch scores, representing the average interaction scores associated to LCR meta-patches and to No LCR meta-patches, respectively. Meta-patches were then categorized based on their contact degree into high-degree (≥2) and low-degree (<2) groups. We evaluated the enrichment of LCR meta-patches within the high-degree group relative to the low-degree group, and statistical significance was assessed using Fisher’s exact test with FDR correction for multiple testing.

### Thermodynamic stability prediction of the RRIs from transcriptome-wide studies

To calculate the ΔG for each RRI and obtain comparable scores within individual datasets, a fixed-size nucleotide region centered on the interaction was selected for each interaction patch. The size of this region was determined *ad hoc* for each dataset to ensure that it fully encompassed 95% of the interactions within the dataset. Then, the ΔG prediction was performed using IntaRNA 2 v2.3.0 software^25^ with default parameters for the standard prediction; *--noSeed* for the prediction without seed and *--outCsvCols* set to extract the individual energy components corresponding to query accessibility penalty (ED1), target accessibility penalty (ED2), and hybridization energy (E_hybrid); *--mode=M --noSeed --qAccW=0 --qAccL=0 --tAccW=0 --tAccL=0 --tIntLenMax=25 -- qIntLenMax=25* and *--noSeed --tAcc=N --qAcc=N* parameters for RNAup^56^ and RNAhybrid^57^ emulation, respectively, as described in https://github.com/BackofenLab/IntaRNA. To ensure comparability across RRI sets of varying length, length-normalized ΔG scores from IntaRNA 2 were used.

For comparison with mock interactions, the patches in each dataset were randomly shuffled within the transcripts using BEDtools shuffle with the parameters *-chromFirst -bedpe*, avoiding the formation of intramolecular interactions. Subsequently, the predictions were conducted using the same criteria described above.

### Cell cultures conditions and treatments

All cell lines used in this study were grown at 37°C, 5% CO2 and tested for mycoplasma contamination. Murine HBG3 ES cells (embryonic stem cells derived from HB9::GFP transgenic mice) were cultured on gelatin-coated or MEF-coated dishes and maintained in mESC medium (Dulbecco’s Modified Eagle’s Medium for ES, 15% Fetal Bovine Serum for ES, 1X GlutaMAX, 1X Non Essential AmminoAcids, 1X nucleosides, 1X 2-mercaptoethanol and 1X Penicillin/Streptomycin) supplemented with LIF (103 unit/mL), FGFRi (1,5 μM) and Gsk-3i (1,5 μM) (LIF+2i condition). Medium was changed every day and cells were passaged every 2–3 days with 1X Trypsin-EDTA solution. Spinal motor neurons (MNs) were differentiated from mESCs HB9::GFP according to Wichterle et al.^108^ and Errichelli et al.^109^. After papain dissociation, cells were plated on polyornithine/laminin-coated dishes with N2B27 medium (50% DMDM F12, 50% Neurobasal, 1X Glutamax, 1X Penicillin-Streptomycin, 1X B27, 1X N2, 200 ng/mL Ascorbic Acid, 20 ng/mL BDNF, 10 ng/mL CNTF, 10 ng/mL GDNF, 10 nM Rhok inhibitor) and differentiation was allowed to proceed for three additional days.

### RNA extraction and quantification by qRT-PCR

Total RNA was isolated from cell cultures using miRNeasy Kit (Qiagen), and retrotranscribed with Superscript Vilo cDNA synthesis Kit (11754050, Invitrogen) according to the manufacturer’s protocol, in a final reaction volume of 10 μL. qRT-PCR was performed using PowerUp SYBR Green Master Mix (A25742, Life Technologies). Relative RNA quantity was calculated as the fold change (FC, 2^−ΔΔCt^) with respect to the control sample set as 1, unless differently specified. Oligonucleotides used for qRT-PCR are provided in **Supplementary Data 3**. DNA amplification was monitored with an ABI 7500 Fast qPCR instrument. Data analysis was performed using the SDS Applied Biosystem 7500 Fast Real-Time PCR system software.

### Native RNA pull-down

Native RNA pull-down was performed in mESC-derived neural mixed population obtained from EB cells at day 6 that were dissociated and replated, allowing differentiation to proceed for additional 3 days (DIV3) as described by Errichelli et al.^109^. The RNA pull-down procedure was performed as described by Martone et al.^74^. The sequences of the biotinylated oligonucleotides belonging to ODD and EVEN probe sets (consisting of six and five probes, respectively), and LacZ control are listed in **Supplementary Data 3**.

### RNA pull-down sequencing and analysis

Library preparation for Input, ODD, EVEN, and LacZ duplicate samples was performed using Stranded Total RNA Prep with Ribo-Zero Plus (Illumina) according to manufacturer’s documentation. The sequencing reaction was performed on an Novaseq 6000 sequencing system at the Genomic Facility in Istituto Italiano di Tecnologia (Genoa, Italy) and produced an average of 43,6 million 150**-**nucleotide long paired-end read pairs. Quality of reads was assessed using FASTQC software v0.11.9 (available at https://www.bioinformatics.babraham.ac.uk/projects/fastqc/). Reads were processed using Cutadapt v3.2^98^ with *-u 1 -U 1 –trim-n –nextseq-trim=20 -m 35* parameters to trim low-quality bases and sequencing artifacts. Reads aligning to the sequence of murine rRNAs (retrieved from NCBI Gene database^110^), 7SL or 7SK transcripts (retrieved from Ensembl database, **Supplementary Data 3**) were filtered out; this first alignment was conducted using Bowtie2 v2.4.2^111^, aligning the first read in each pair with the *--nofw* parameter and the second read with the *--norc* parameter. Only read pairs in which both mates did not map were retained for downstream analysis. STAR v2.7.7a software^99^ was used to align reads to GRCm38 genome using the following parameters: *-- outSAMstrandField intronMotif --outSAMattrIHstart 0 --outFilterType BySJout -- outFilterMultimapNmax 20 --alignSJoverhangMin 8 --alignSJDBoverhangMin 1 -- outFilterMismatchNmax 999 --outFilterMismatchNoverLmax 0.04 --outFilterIntronMotifs RemoveNoncanonical --peOverlapNbasesMin 50* l. PCR duplicates were removed from all samples using MarkDuplicates command from Picard v2.24.1 suite^112^. Uniquely mapping fragments from de-duplicated BAM files were counted for each annotated gene (Ensembl release 99) using Htseq v0.13.5 software^113^. edgeR v3.34.1 R package^102^ was used to compare pull-down-enriched RNAs to their relative Input samples. RNAs with p-value <0.05 and log_2_FC > 0.58 in both ODD versus Input and EVEN versus Input contrasts, and log_2_FC < 0 in the LacZ versus Input contrast were defined as enriched. Additionally, only RNAs with expression levels exceeding 1 FPKM (Fragments Per Kilobase of exon per Million mapped reads, calculated using the edgeR *rpkm* function) in the Input sample were kept. Isoform quantification was performed with Salmon v1.6.0 software^114^ using *– libType ISR –validateMapping*s parameters and a full decoy transcriptome index created using the Ensembl 99 transcriptome and the GRCm38 genome. In case of genes with multiple isoforms, the most expressed isoform in the Input sample (according to Salmon TPM quantification) and with the same biotype of the gene was selected for downstream analyses.

The top 300 TINCR interactors were selected based on the ‘log2_escore’ column reported in the table of the 100 nucleotide-long regions enriched in the TINCR pull-down (retrieved from the GEO record GSE40121). Only interactors also present in the Ensembl release 99 database were included in the top set. A re-analysis of the TINCR pull-down experiment^16^ was performed to calculate log_2_FC values between Pull-down samples and Input, enabling the identification of RNAs primarily depleted in the pull-down (log_2_FC < -1), from which controls were selected. The data were downloaded from GEO (GEO accession GSE40121) using fastq-dump from SRA Toolkit v2.10.9. Subsequently, reads were aligned to the GRCh38 genome using STAR v2.7.7a^99^ with the ENCODE standard parameters, along with the following additional settings: *--outSAMstrandField intronMotif --outSAMattrIHstart 0 --outFilterIntronMotifs RemoveNoncanonical*. The subsequent steps, including de-duplication and quantification of uniquely mapping fragments for each annotated gene (*Homo sapiens*, Ensembl release 99 annotation), were performed as previously described.

### Generation of negative controls for lncRNAs’ targets and repeat enrichment analysis

For each of the identified RNA interactors of the Lhx1os or TINCR lncRNAs, 5 and 3 negative controls, respectively, were selected for nucleotide-based RNA–RNA interaction prediction analysis and screened for the occurrence of interspersed repeats. The controls were chosen from transcripts expressed in the Input samples (FPKM > 1) but depleted in the pull-down extracts. For Lhx1os, we selected controls within the bottom 50% of log_2_FC values in the pull-down versus Input comparison. For TINCR, controls were selected from transcripts with log_2_FC values below -1 in the same comparison. The controls were selected to match the corresponding RNA targets in gene and transcript biotypes, as well as in 5’ UTR, CDS, 3’ UTR, and overall transcript length. The selection process followed these steps: (I) 5’ UTR, CDS, 3’ UTR, and overall lengths were obtained from Ensembl Release 99 using BioMart; (II) for each interactor, the percentage difference (%Δ) between the interactor and candidate controls was calculated for each parameter; (III) the %Δ values across all parameters were summed for each candidate; (IV) controls with the lowest total %Δ and matching biotypes were selected. Each control was uniquely assigned to a single interactor to prevent redundancy. Finally, controls were tested using the Wilcoxon-Mann-Whitney test to confirm the absence of significant differences compared to interactors (p-value > 0.05). For each repeat family and element, we used Fisher’s exact test to compare the proportions of targets and controls containing them. P-values were then adjusted for multiple comparisons using the Benjamini-Hochberg procedure.

### RNA contact enrichment analysis

Initially, TINCR and Lhx1os sequences were segmented into 50- or 10-nucleotide windows with steps of 10 or 1 nucleotide, respectively. The resulting bins were then compiled into a BED file. To predict the most favorable interactions between the analyzed RNA and each transcript in the target or control group, we used IntaRNA 2 software with standard parameters, ASSA v1.0.1 and pRIblast v0.03 (a parallelized implementation of RIblast^115^). Interaction predictions for both the target and control groups were then mapped to the lncRNA bins using BEDtools intersect. For each bin, the statistical significance of the over-representation of contacts with pull-down RNAs compared to contacts with control RNAs was evaluated using Fisher’s exact test. Additionally, the difference between the ΔG distributions obtained for pull-down RNAs and controls was assessed using the Mann-Whitney U-test. Benjamini-Hochberg correction was applied for both tests.

Meta-region analysis of predicted interaction patches in the pull-down and control groups was conducted as follows. A region extending from -100 nt to +100 nt around the middle point of interaction patches predicted on target and control RNAs was analyzed using a sliding window of 10 nucleotides, with steps of 1 nucleotide. Windows falling partially or totally outside transcripts were discarded. Then, RepeatMasker v4.1.1 was used to identify repeated elements in pull-down and control RNA groups, and BEDtools intersect was used to assign repeats to the overlapping windows. Also the interaction patches were assigned to windows in order to calculate the frequency of contacts in each window. Finally, a Fisher’s exact test was performed to test the enrichments of repeat-overlapping predicted interaction patches in the pull-down group compared to control RNAs. P-values were corrected for multiple testing using the Benjamini-Hochberg method.

The identification of Lhx1os 7-mers complementary to GA-rich low-complexity sequences was performed as follows: since no specific annotations for GA-rich (-) or TC-rich (+) sequences are available in RepeatMasker, we retrieved GA-rich low-complexity sequences annotated by the software within the mESC-derived neural mixed population transcriptome (i.e. Input samples of Lhx1os pull-down experiment). These sequences were then partitioned into 7-mers, and the reverse-complementary sequences were mapped onto the Lhx1os transcript.

### Evaluation of Nucleotide Transformer and RNA language models

The language models Nucleotide Transformer (NT-v1, Multispecies 2.5B model)^30^, RNA-FM^62^, and RNAErnie v1.0^63^ were evaluated for their ability to capture information relative to LCRs and Minimum Free Energy (MFE) of RNA folding. Starting from the Ensembl 99 human transcriptome, LCRs were annotated as described in the “Detection of repeated elements section”. LCRs included in CDS regions were excluded to ensure a fair comparison across models, as RNA-FM and RNAErnie were trained exclusively on non-coding transcripts. A control set was generated using BEDTools shuffle, randomly repositioning each LCR while preserving its regional context— ensuring that elements originally located within specific transcript regions (5′UTR, 3′UTR or non-coding RNA) were shuffled only within the same region type. Transcripts were subdivided into consecutive 200-nt windows using BEDTools makewindows with parameters -w 200 and -s 200. From these, 1,000 windows overlapping LCRs by at least 30 nucleotides (LCR windows) and 1,000 windows overlapping shuffled controls by at least 30 nucleotides (CTRL windows, without any overlap with LCRs) were randomly selected. The sampling procedure was designed to include an equal number of windows derived from the same transcript regions in both LCR and CTRL window sets, maintaining the following proportions: 5% from 5′UTRs, 60% from 3′UTRs, and 35% from non-coding RNAs. To compute the NT, RNA-FM, and RNAErnie embeddings for each selected window while preserving the appropriate sequence context, the largest possible sequence segment was extracted according to the maximum input length supported by each pretrained model (6,000 nt for NT, 1,022 nt for RNA-FM, and 512 nt for RNAErnie). When transcript sequences were shorter than these limits, the full sequence was used. For NT, which employs 6-mer tokenization, the maximum sequence length was adjusted to the nearest multiple of six. For each window and model, the mean embedding vector was obtained by averaging the embeddings of all tokens overlapping that window.

An additional classification of the 200-nt windows was performed according to RNAfold v2.4^23^ predictions. Based on the MFE distribution, windows were divided into two groups, labeled “low MFE” (below the median) and “high MFE” (above the median).

Finally, logistic regression classifiers were trained using the scikit-learn v0.24.2 Python package^116^ to distinguish between LCR and CTRL windows, or between low and high MFE windows, based on the mean embedding vectors. Model accuracy was evaluated using 10-fold cross-validation, with 10% of the data used for testing in each fold.

### RIME architecture

RIME is a deep learning model that leverages Nucleotide Transformer (NT-v1, Multispecies 2.5B model)^30^ embeddings to predict RNA–RNA interactions. It first takes two RNA sequences as input, each represented as a series of 6-mer tokens. The NT model then transforms these tokenized sequences into 2,560-dimensional vectors, where each token captures the global sequence context. We introduced a Contact Matrix (CM) operation in our pipeline to map the two embeddings into a contact matrix, where each spatial point corresponds to a token pair combination from the two RNA sequences. Contact matrices provide a compact way to represent the spatial relationships between nucleotides and have been already used for secondary structure prediction^117^. The feature space of the contact matrix is formed by concatenating the respective embeddings of the token pairs. The output tensor can be seen as an image, and we designed specific modules commonly employed in computer vision. First, since the dimensionality of each ‘pixel’ is too high after concatenation (2x2,560), we jointly learned an independent transformation, denoted as ψ, to reduce the dimensionality to 800. Second, we designed a Feature Extractor transformation that effectively mixes the information across both spatial and feature spaces using standard convolutional neural network (CNN) layers. The advantage of convolutional layers is their length independence, which allows our pipeline to handle RNA sequences of varying lengths. Moreover, they ensure computational efficiency compared to attention-based models, which have quadratic complexity O(n²) with respect to input length n, while convolutional layers scale linearly (O(n)), are highly parallelizable, and thus generally more efficient on long inputs^118^. Each input contact matrix has different dimensions, so we standardized them by applying global average pooling. The final layer for binary prediction is a linear layer with bias. RIME architecture is schematized in **Supplementary Fig. 6d**.

Hyperparameters, including ψ dimensionality (800), convolutional channels (300 for each layer), batch size (32), and learning rate (5×10□□), were selected based on a random search over a space of 243 possible configurations, sampling 121 to balance coverage with computational cost. To further accelerate the search, each configuration was trained for a single epoch. This approach, supported by recent evidence, has been shown to reliably separate promising from poor hyperparameter configurations while avoiding the expense of full training^119^. Model performance was evaluated on the validation set using the weighted cross entropy loss function with a class weight ratio of 1.5:1.0, as explained in the section about RIME training. The random search was performed over a space defined by batch sizes of 8, 16, or 32, learning rates of 5×10□□, 1×10□□, or 5×10□□, ψ output dimensionalities of 200, 500, or 800, and convolutional channels of 100, 300, or 600 in both layers. We selected hyperparameter values based on their overall behavior across sampled configurations, excluding consistently poor performers and screening the remaining ones to balance predictive performance with training efficiency. A batch size of 32 was ultimately selected because it offered a favorable compromise between computational cost and stability, reducing the number of parameter updates required per epoch without degrading performance.

Given this relatively large batch size, we adopted the highest learning rate to accelerate convergence. While a learning rate of 5×10^-5^ did not yield uniformly superior mean validation performance, it exhibited greater variance across runs—consistent with the expectation that higher learning rates explore the loss landscape more broadly and can escape sharp local minima, provided that subsequent weight decay stabilizes training. Accordingly, we used 5×10^-5^ with the AdamW optimizer, applying a weight decay of 1×10^-4^ throughout training. For the multi-layer perceptron output dimensionality, we explored several configurations to reduce the 2×2,560-dimensional input embeddings. No clear monotonic trend was observed across the tested values; we selected 800 units as a compromise that reduced the input dimensionality by a factor of ∼6 (compared with ∼25 for 200 units, which proved overly restrictive), thus preserving a greater fraction of embedding information while remaining computationally feasible. For the convolutional channels, one-epoch evaluations indicated that 600 channels tended to yield the poorest results, whereas configurations between 100 and 300 showed similar performance. We therefore selected 300 channels, providing greater representational capacity during full training while maintaining computational feasibility.

This choice is consistent with the observation that single-epoch evaluations introduce some sampling variability, making it preferable to favor slightly more expressive configurations for subsequent full training.

RIME was developed using the Pytorch v1.9.1 Python library^120^. It comprises 4,184,094 trainable parameters; for input windows of 200×200 nucleotides, the model performs ∼8.7 GFLOPs per forward pass, which is computationally comparable to ResNet-50 on 224×224 pixel images.

### Construction of positive and negative sets for RRI prediction tasks

The positive set used to train and evaluate the RIME model was generated from the RRI collection derived by Psoralen-based experiments. To enhance data consistency and avoid redundancy, we mapped overlapping interaction sites from different experiments into a unified interaction region, representing the minimal region encompassing all identified interaction sites and providing a cohesive view of the interaction dynamics. To provide sufficient context to the model, the positive set consisted of two-dimensional windows encompassing the interaction sites. The negative set was composed of non-interacting windows generated from the interacting transcripts according to the following criteria. The NP negative set was created by randomly selecting windows from the contact matrices of interacting transcript pairs, while ensuring that interacting sites were excluded from the selection. To generate the windows that compose the PN and NN classes, we first generated random transcript pairs, ensuring that they did not interact with each other, thus creating non-interacting contact matrices. Then, NN windows were generated by randomly selecting RNA regions from these matrices. Finally, PN windows were positioned in regions encompassing interaction patches of both transcripts. The described creation of non-interacting contact matrices was performed by a random pair-swapping approach, designed to prevent biases while preserving experimental context and biological relevance. We restricted swapping within each RRI dataset to ensure negative pairs reflected the specific experimental conditions and remained consistent with the original data. We also confined the non-interacting transcript pairs generation by preserving the biotype composition of interacting sites. In particular, we categorized each interacting site into four types - CDS, 3’UTR, 5’UTR (protein-coding transcripts), and ncRNA (non-coding transcripts) - yielding 16 possible combinations (e.g., CDS-CDS, CDS-3’UTR). We confined swapping within these combinations to ensure negative pairs matched the interaction region of their positive counterparts, maintaining a consistent genetic grammar. We generated two non-interacting contact matrices for each interacting one where possible, though the exclusive presence of a single interaction type for certain biotype composition could prevent the generation of non-interacting contact matrices. In this process, we ensured that the generated transcript pairs were never seen interacting in the following datasets: RIC-seq, PARIS2 and MARIO, PARIS1, RAIDv2.0, CLASH, SPLASH, NPInterv3.0, RIAseq, RAINv1.0, RAP, LIGRseq from RISE database.

Building on this approach, we evaluated the problem of predicting RRIs by considering two distinct evaluation perspectives: DRP, whose aim is to classify PPs as positives and NNs and NPs as negatives, and DRI, whose aim is to classify PPs as positives and PNs as negatives. The DRP task assesses the model’s ability to identify interaction-prone regions, while the DRI task evaluates its accuracy in identifying the transcript pairs that are actually interacting.

The Psoralen-based dataset was split into training (70%), validation (15%) and test (15%) sets. To ensure similar RNA composition between positives and negatives, we performed the randomized combination procedure generating the non-interacting contact matrices within each set.

Furthermore, we ensured that the training, test, and validation sets maintained similar contact matrix area distribution and RNA region composition, reflecting the properties of the original dataset (**Supplementary Fig. 6e**, right panel, and **h**).

### Data preparation for RIME training

To accommodate the NT model’s input sequence limit of 6,000 nucleotides, we processed each sample into sub-sequences of up to this length, resulting in sub-contact matrices with dimensions of up to 6,000×6,000 nucleotides. To note, Nucleotide Transformer operates on 6-mers, encoding each 6,000-nucleotide region as a sequence of 1,000 embedding tokens. To enhance the coverage of longer transcript pairs during training, additional embeddings were generated. Specifically, for contact matrices where both RNAs exceeded 7,000 nucleotides, two embeddings were created (22% of the training set), and for pairs exceeding 8,500 nucleotides, three embeddings were generated (2%).

The computation of NT embeddings required ∼1.5 days for ∼150,000 sequences.

Once embeddings were created, we sampled contact matrix windows corresponding to four classes—PP, PN, NP, and NN—which collectively formed the training set. Each window captured smaller segments of the original embeddings but, since NT embeddings are designed to encode each token within the context of the entire sequence, the broader sequence context was preserved. As a result, the feature space of each window reflected the influence of the full embedded transcript.

In each epoch, we used a balanced sampling strategy across three classes of negative samples: PNs made up 50% of the total negative samples, while NPs and NNs each contributed 25%. This resulted in a balanced sample distribution between DRI and DRP tasks.

To create a balanced training set and prevent the model from being biased toward positive or negative interactions, the total number of negative samples was matched to the number of PP samples. This was achieved by adjusting the sampling probabilities of NN, NP, and PN windows so that their combined total matched the number of PPs, while preserving the 50%-25%-25% distribution of PN, NN, and NP windows.

To evaluate whether the sampling procedure introduced representation bias, we monitored RNA occurrence across positive and negative sets over 10 training epochs. This analysis was repeated under different negative sampling schemes—using the RIME distribution (50% PN, 25% NN, 25% NP), or exclusively PN, NN, or NP negatives—always maintaining balanced sets.

During training, the four classes—PN, NN, NP, and PP—were sampled from varying regions of the embeddings across epochs, increasing coverage of the 6,000-nucleotide embeddings, enhancing sequence context diversity, and mitigating overfitting.

Additionally, during training we sampled positive and negative windows with side lengths that varied according to a distribution centered at 200 nucleotides (**Supplementary Fig. 6f** and g), equalizing the length distributions across classes. This dynamic sampling ensured that the same interaction site (for PP) was sampled with varying granularity and sequence context, preventing memorization, enhancing generalization, and enabling the model to handle varying lengths.

To accommodate batch training with transcripts of different lengths, we normalized tensor lengths within each batch to match the shortest sequence by averaging groups of tokens. Specifically, for each longer tensor, we divided it into groups such that the number of groups equaled the shortest tensor length. We then computed the mean value for each group, resulting in a reduced tensor that matched the shortest tensor length. The number of tokens in each group was determined by the ratio of the tensor length to the shortest batch length (with any remainder grouped into the last group).

This approach was made possible since the data within the batch represent a vector space comparable with the original embedding space, with original sequences treated as tensors with group sizes of one. This averaging acted as an additional form of augmentation, avoiding the inefficiencies of zero-padding, which would otherwise lead to the creation of sparse matrices. As an additional augmentation technique, we also reversed the order of transcript pairs during training, ensuring that the model did not develop an order-based bias, allowing it to learn from varied configurations of the same data.

### Data preparation for RIME evaluation

The Psoralen-based validation set, Psoralen-based test set, as well as the RIC-seq and MARIO test sets, consisted of 200×200 nucleotide positive and negative windows.

For each interacting transcript pair, we generated a 6,000x6,000 contact matrix embedding, sampling one PP and one NP window. Similarly, for non-interacting pairs, we created embeddings and sampled one NN and one PN window. For transcript pairs longer than 8,500 nucleotides (5% in RIC-seq, 1% in MARIO, 2% in the Psoralen-based test set), we generated additional embeddings without interaction sites (and without randomly combined sites for non-interacting pairs), sampling one NP window for interacting pairs and one NN window for non-interacting pairs.

We excluded all interacting transcript pairs observed in the Psoralen-based training set from both the RIC-seq and MARIO datasets to ensure an out-of-sample evaluation context. Additionally, we excluded pairs where the interaction site on either transcript exceeded 200 nucleotides.

All the results for the DRI and DRP tasks were evaluated by balancing the positive and negative sets according to the proportions of negatives considered for the specific task.

Testing was conducted on the full Psoralen-based, RIC-seq, and MARIO datasets, as well as on subsets with progressively higher quality, determined by selecting interactions based on the number of supporting reads and the minimum length of the interacting region. To calculate AUC values, for each threshold negative samples were randomly undersampled 100 times to match the number of positive samples, yielding 100 ROC-AUC values, which were then averaged to calculate the final performance metric. The same undersampling-based approach was adopted to calculate Accuracy, Precision, NPV, Recall, and F1 score (**Supplementary Data 5**). To calculate Precision and NPV against RIME’s confidence (**Supplementary Fig. 8e**), the imbalance between the positive and the negative sets was addressed by undersampling before ranking and selecting the top and bottom predictions at each data proportion threshold. For each threshold the scores were averaged over 50 iterations.

### RIME training and model selection

RIME was trained for 74 epochs with early stopping and a patience of 30 epochs, using a batch size of 32. The model that achieved the highest F1 score on the validation set was selected. The F1 score was calculated on a subset of the validation set that mirrored the composition of the training set, maintaining the 50%-25%-25% distribution of PN, NN, and NP windows and ensuring their total matched the number of the entire set of positive windows. To ensure an unbiased F1 score the negative set undersampling process was repeated 50 times, resulting in equal numbers of positive and negative pairs for each iteration. The F1 scores were then averaged across the iterations. In each epoch, we uniformly and randomly sampled half of all possible RRIs in the Psoralen-based training set, ensuring that, on average, the model was exposed to the entire set of interacting sites every two epochs.

To prioritize the reliability of positive interaction predictions, we optimized a weighted cross entropy loss function with a class weight ratio of 1.5:1.0, giving greater emphasis to the positive class. We used the AdamW optimizer with a learning rate of 5 × 10^−5^, weight decay as 1× 10^−4^, and standard values for other hyperparameters: β1 = 0.9, β2 = 0.999, ε = 1 × 10^−8^. RIME was trained on an NVIDIA Quadro RTX 5000 GPU with a peak memory usage of 3.6 GiB. To assess the impact of the negative sample selection strategy, we trained additional RIME models restricted to a single task. For the DRI setting, the negative set consisted entirely of PN windows, while for the DRP setting negatives were evenly split between NN and NP windows. For each task, two variants were trained: one using the same validation composition as RIME (50% PN, 25% NN, 25% NP) and one using a validation set mirroring the task-specific training distribution (PP/PN for DRI and PP with 50% NN and 50% NP for DRP), resulting in a total of four models.

We trained RIMEfull by expanding the dataset size through the incorporation of the validation set into the training phase. Model selection was performed on a subset of the Psoralen-based test set, chosen for its reliability and the strong performance of RIME on high-quality interactions. The positive class included interactions with region length ≥ 35 nucleotides, or supported by ≥ 6 PARIS1 reads, or ≥ 3 PARIS2 reads. RIMEfull was trained for 86 epochs, with the F1 score used to select the best model, while all other hyperparameters were set identically to those used for RIME training. Since the entire Psoralen-based dataset was used for training and model selection, the performance of RIMEfull was evaluated on the external RIC-seq and MARIO datasets.

### Comparative benchmark of RNA–RNA interaction prediction algorithms

The benchmark of RNA–RNA interaction prediction algorithms was performed evaluating RIME performances together with ASSA v1.0.1, IntaRNA v3.4.0 (with IntaRNA2 personality), pRIblast v0.03, RIsearch2 v2.1, RNAcofold v2.6.4, RNAplex 2.6.4, RNAhybrid, RNAup and SPOT-RNAc. All the tools were evaluated on the same set of 200×200 windows already described in the “Data preparation for RIME evaluation” section. For each analyzed window, we measured the interaction propensity score of the thermodynamics-based algorithms as the minimum ΔG value returned by the software. ASSA was used with *--all_sites* parameter; RNAcofold with *--output-format=D*; for RIsearch2, we retained only RNA–RNA interactions filtering out predictions with negative strand; RNAhybrid and RNAup algorithms were emulated by IntaRNA 2 software as previously specified. SPOT-RNAc was employed following the guidelines in the benchmark work by Lang et al.^71^. Specifically, we employed SPOT-RNA^70^ (https://github.com/jaswindersingh2/SPOT-RNA), rather than SPOT-RNA2^121^, as the former performs better for intermolecular interactions, using the code available at https://github.com/meilanglang/RNA-RNA-Interaction. The method was initially designed to predict secondary structure, but proved effective in predicting intermolecular base-pairing probabilities when concatenating two input RNAs (hence the c in SPOT-RNAc). For each window, the two input RNAs were concatenated without a linker, and base-pair probability matrices were generated for both sequence orders (AB and BA). For each matrix, we computed the mean probability of intramolecular base pairs, and then derived a final prediction score as the average of the two means. This scoring approach yielded better performance than either the maximum intramolecular probability or the difference between average intermolecular and intramolecular probabilities. For all the other tools we used standard parameters.

### Grad-CAM analysis

To enhance the interpretability of our classifier, we used Grad-CAM, an explainable Artificial Intelligence technique originally designed for computer vision tasks. Grad-CAM provides spatial explanations for model predictions by analyzing how gradients influence neurons in the last convolutional layer of the CNN, thereby highlighting the most relevant regions. Grad-CAM predictions are class-specific, and we focused on the positive class for our analyses. In our model, Grad-CAM was applied after the last convolutional layer, where it is expected to capture optimal high-level semantics. Since the convolutional blocks reduced the spatial dimensions of the RNA– RNA contact matrix, the resulting Grad-CAM matrix also had lower resolution. To align its dimensions with the original contact matrix, we resized the 2D Grad-CAM localization map using nearest-neighbor interpolation.

### Preparation of RRI sets for RIMEfull evaluation

The COMRADES data relative to NORAD lncRNA were retrieved from the GEO database under accession number GSE188445, considering only untreated samples. Chimeric reads associated with the NORAD transcript (ENSG_ENST_NORAD_lincRNA) were retained, excluding those corresponding to secondary structures. For each potential RNA interactor we quantified the number of chimeric reads mapped in COMRADES experiments with proper cross-linking (GSM5683127_Nr1 [replicate 1], GSM5683128_Nr2 [replicate 2], and GSM5683129_Nr3 [replicate 3]) and in control samples (GSM5683135_Nr1C [replicate 1], GSM5683136_Nr2C [replicate 2], and GSM5683137_Nr3C [replicate 3]). To identify NORAD-specific interactors, we evaluated the enrichment of chimeric reads for each RNA target in the COMRADES samples compared to the controls. This analysis was conducted independently for each replicate, and NORAD interactors were defined as RNAs with a significant enrichment of chimeric reads in COMRADES in at least two replicates. The statistical significance of the enrichment was determined using Fisher’s exact test, selecting targets with log_2_(Odds Ratio) > 1 and p-value < 0.05. To annotate the *bona fide* interacting regions, chimeric read arm sequences were re-mapped using the regex v2.4.104 Python library to the NORAD sequence (NR_027451.1) and to that of the identified interactors, retrieved from Ensembl 99 annotations. Reads with multiple alignments or more than five nucleotide mismatches were excluded to ensure high-confidence mappings. RIMEfull predictions between NORAD and its identified targets was performed on this transcript sequence set.

The PGLYRP3 region that interact with TINCR lncRNA was retrieved from the work from Kretz and colleagues considering the TINCR box-depleted PGLYRP3 mutant (PGLYRP3Δ96 bp). lnc-SMaRT interacting regions with Mlx-γ and Spire1 transcripts were retrieved from the works of Martone and colleagues^50^ and Scalzitti and colleagues^51^, respectively.

### Evaluation of RIMEfull’s performance on intramolecular RNA interactions

Reads supporting NORAD intramolecular contacts were extracted from COMRADES experiments GSM5683127_Nr1 (replicate 1), GSM5683128_Nr2 (replicate 2), and GSM5683129_Nr3 (replicate 3), selecting only the chimeric ones in which both arms mapped to the NORAD sequence (ENSG_ENST_NORAD_lincRNA) and allowing a maximum of five mismatches with the sequence. The NORAD sequence (NR_027451.1) was divided into 200-nt bins with a 100-nt step size. A pairwise contact matrix was then generated representing all the possible bin pairs. Based on the genomic coordinates of each bin pair, NORAD chimeric reads were mapped using BEDTools pairtopair with the parameter *-type both*, thus enabling the quantification of chimeric reads associated with each cell. For each replicate, counts in each cell were normalized as CPM (counts per million) relative to the total number of NORAD–NORAD chimeric reads, and an intramolecular contact index was defined as the mean CPM across the three replicates. Cells were then stratified into ten equally sized groups according to increasing intramolecular contact index values. To generate a control set, such indices were randomly shuffled across cells. RIMEfull scores were subsequently computed for each cell, and an empirical cumulative distribution function (ECDF) of RIMEfull scores was derived for each group.

To assess RIMEfull’s ability to recapitulate thermodynamics-based secondary structure predictions, a random set of 100 human transcripts longer than 2000 nucleotides was selected from Ensembl 99 annotation. Their secondary structure was predicted with RNAfold v2.4, setting the *-p* parameter to compute the partition function and obtain base-pairing probability matrices. The probability matrices were parsed into overlapping 200×200 nucleotide windows with a step size of 100, and the mean base-pairing probability within each window was calculated. This allowed for a direct comparison with RIMEfull prediction scores obtained for the corresponding RNA segment pairs. For each transcript, we computed the Spearman correlation between RIMEfull scores and mean base-pairing probabilities, yielding a distribution of correlation coefficients. This was contrasted with a null distribution obtained by randomly shuffling RIMEfull scores 1,000 times and recalculating the Spearman correlation with mean base-pairing probabilities. The difference between the two distributions was assessed using the Mann-Whitney U-test.

To evaluate the concordance between RIMEfull’s predictions and transcriptome-wide RNA structure probing data, we retrieved RNA accessibility profiles measured using DMS-MaPseq in K562 cells by Rouskin et al.^76^ from the RASP database^122^. We focused on human lncRNAs between 200 and 6000 nucleotides in length selected from the Ensembl 99 annotation. Each transcript was segmented into 200-nucleotide segments with a step size of 100 nucleotides. Transcripts were retained if they contained at least 15 segments with a minimum of 20 nucleotides covered by valid DMS accessibility scores. This filtering yielded 47 transcripts, corresponding to 1,087 windows for downstream analysis, for which we calculated the average accessibility score across the positions with available scores. For each segment, we ran RIMEfull against itself, as well as against segments located 50 nucleotides upstream and downstream (when available), and calculated the mean of the three resulting scores to obtain a single measure describing the segment’s local base-pairing tendency. For each transcript, we then computed Spearman correlation coefficients between these local pairing scores and mean DMS accessibility values. As before, the resulting distribution was compared with a null distribution, produced by randomly shuffling the local pairing scores, using the Mann-Whitney U test.

### RIME web server

To enable access to RIME, we developed a user-friendly web server featuring the RIMEfull model for RNA–RNA interaction prediction. The platform, freely accessible at https://tools.tartaglialab.com/rna_rna, allows users to input two RNA molecules and compute RIMEfull prediction scores across 200×200 nucleotide windows with a 100-nucleotide step. The results are visualized as a heatmap, which allows to identify RNA region pairs with a high likelihood of interaction.

## Supporting information

Description of Supplementary Data

## Data availability

The RNA-seq data generated in this study have been deposited in the Gene Expression Omnibus (GEO) under accession number GSE290073, https://www.ncbi.nlm.nih.gov/geo/query/acc.cgi?acc=GSE290073. Reviewers can access these data using the following token: mpunyiqgnvqtzav.

Raw reads from Flp-In T-REx 293 cells under control, STAU1 knockdown, and rescue conditions were downloaded from the European Nucleotide Archive (ENA) using the following accessions: ERR605044, ERR605045, and ERR605046 for the Ribo-seq samples, and ERR618765, ERR618766, and ERR618767 for the corresponding RNA-seq libraries.

Bulk RNA-seq data used to study RNA editing and measure gene expression were retrieved from GEO under the accession numbers GSE99249, GSE198386, GSE94387, GSE217878, and GSE160578.

TINCR RIA-seq data were retrieved from GEO under the under accession number GSE40121. NORAD COMRADES data were retrieved from GEO under the under accession number GSE188445.

## Code availability

A web server leveraging the RIMEfull model for RNA–-RNA interaction prediction is freely available at https://tools.tartaglialab.com/rna_rna. A repository providing code, RIMEfull model weights and instructions is available at https://github.com/tartaglialabIIT/RIME. Code and model weights are also available in Zenodo at https://zenodo.org/records/14926723, along with training, validation and test sets.

## Author contributions

A.S. and A.C. retrieved the RRI datasets analyzed in the present study. A.S. performed most of the bioinformatics data analysis, including repeat enrichment evaluation and thermodynamic characterization of RRIs, with help from A.D. for RNA editing and secondary structure analysis. G.B. conceived, developed and tested RIME, formalizing the DRP and DRI model evaluation tasks, with help from V.M., M.M. and E.R. in model design and negative samples generation, from G.P. in GRAD-CAM analyses, and D.M.V. in benchmarking RIME against thermodynamics-based tools. F.P. performed and analyzed the Lhx1os pull-down experiment. A.C. and G.G.T. conceived the study and wrote the manuscript, with contributions from A.S. and G.B. I.B., G.R., J.M. and M.M. provided valuable input in the interpretation of data and results. All the authors read and approved the manuscript.

## Acknowledgements

The authors would like to thank the ‘RNA initiative’ at IIT and all the members of Tartaglia’s and Irene Bozzoni’s lab at Sapienza and IIT. Special recognition goes to Dr. Davide Mariani for expertly curating the sequencing library preparation. The authors would also like to thank Prof. Yuanchao Xue and Dr. Changchang Cao for providing RIC-seq data, as well as Prof. Jernej Ule and Dr. Anob M. Chakrabarti for sharing STAU1 hiCLIP data.

## References

1. Dai, X., Zhang, S. & Zaleta-Rivera, K. RNA: interactions drive functionalities. Molecular Biology Reports 47, 1413–1434 (2019).

2. Romero-Barrios, N., Legascue, M. F., Benhamed, M., Ariel, F. & Crespi, M. Splicing regulation by long noncoding RNAs. Nucleic Acids Research 46, 2169 (2018).

3. Werner, A., Kanhere, A., Wahlestedt, C. & Mattick, J. S. Natural antisense transcripts as versatile regulators of gene expression. Nature Reviews Genetics 25, 730–744 (2024).

4. Singh, S., Shyamal, S. & Panda, A. C. Detecting RNA-RNA interactome. Wiley interdisciplinary reviews. RNA 13, (2022).

5. Bose, M., Rankovic, B., Mahamid, J. & Ephrussi, A. An architectural role of specific RNA–RNA interactions in oskar granules. Nature Cell Biology 26, 1934–1942 (2024).

6. Lu, Z. et al. RNA Duplex Map in Living Cells Reveals Higher-Order Transcriptome Structure. Cell 165, 1267–1279 (2016).

7. Zhang, M. et al. Optimized photochemistry enables efficient analysis of dynamic RNA structuromes and interactomes in genetic and infectious diseases. Nature Communications 12, 1–14 (2021).

8. Aw, J. G. A. et al. In Vivo Mapping of Eukaryotic RNA Interactomes Reveals Principles of Higher-Order Organization and Regulation. Mol Cell 62, 603–617 (2016).

9. Sharma, E., Sterne-Weiler, T., O’Hanlon, D. & Blencowe, B. J. Global Mapping of Human RNA-RNA Interactions. Molecular cell 62, (2016).

10. Wu, T. et al. KARR-seq reveals cellular higher-order RNA structures and RNA–RNA interactions. Nature Biotechnology 1–12 (2024).

11. Cai, Z. et al. RIC-seq for global in situ profiling of RNA–RNA spatial interactions. Nature 582, 432– 437 (2020).

12. Nguyen, T. C. et al. Mapping RNA–RNA interactome and RNA structure in vivo by MARIO. Nature Communications 7, 1–12 (2016).

13. Skeparnias, I. & Zhang, J. Cooperativity and Interdependency between RNA Structure and RNA-RNA Interactions. Non-coding RNA 7, (2021).

14. Tieng, F. Y. F. et al. A Hitchhiker’s guide to RNA–RNA structure and interaction prediction tools. Briefings in Bioinformatics 25, bbad421 (2023).

15. Adinolfi, M. et al. Discovering sequence and structure landscapes in RNA interaction motifs. Nucleic acids research 47, (2019).

16. Kretz, M. et al. Control of somatic tissue differentiation by the long non-coding RNA TINCR. Nature 493, (2013).

17. Liang, L. et al. Complementary Alu sequences mediate enhancer–promoter selectivity. Nature 619, 868–875 (2023).

18. Duszczyk, M. M., Wutz, A., Rybin, V. & Sattler, M. The Xist RNA A-repeat comprises a novel AUCG tetraloop fold and a platform for multimerization. RNA 17, 1973 (2011).

19. Pintacuda, G., Young, A. N. & Cerase, A. Function by Structure: Spotlights on Xist Long Non-coding RNA. Front Mol Biosci 4, 90 (2017).

20. Verheyden, N. A. et al. A high-resolution map of functional miR-181 response elements in the thymus reveals the role of coding sequence targeting and an alternative seed match. Nucleic Acids Res 52, 8515–8533 (2024).

21. Wong, M. S., Shay, J. W. & Wright, W. E. Regulation of human telomerase splicing by RNA:RNA pairing. Nature Communications 5, 1–6 (2014).

22. Gong, C. & Maquat, L. E. lncRNAs transactivate STAU1-mediated mRNA decay by duplexing with 3’ UTRs via Alu elements. Nature 470, 284–288 (2011).

23. Lorenz, R. et al. ViennaRNA Package 2.0. Algorithms for Molecular Biology 6, 1–14 (2011).

24. Busch, A., Richter, A. S. & Backofen, R. IntaRNA: efficient prediction of bacterial sRNA targets incorporating target site accessibility and seed regions. Bioinformatics (Oxford, England) 24, (2008).

25. Mann, M., Wright, P. R. & Backofen, R. IntaRNA 2.0: enhanced and customizable prediction of RNA-RNA interactions. Nucleic Acids Res 45, W435–W439 (2017).

26. Umu, S. U. & Gardner, P. P. A comprehensive benchmark of RNA–RNA interaction prediction tools for all domains of life. Bioinformatics 33, 988–996 (2016).

27. Delli Ponti, R., Marti, S., Armaos, A. & Tartaglia, G. G. A high-throughput approach to profile RNA structure. Nucleic Acids Res. 45, e35 (2017).

28. Sato, K., Akiyama, M. & Sakakibara, Y. RNA secondary structure prediction using deep learning with thermodynamic integration. Nat. Commun. 12, 941 (2021).

29. Ji, Y., Zhou, Z., Liu, H. & Davuluri, R. V. DNABERT: pre-trained Bidirectional Encoder Representations from Transformers model for DNA-language in genome. Bioinformatics (Oxford, England) 37, (2021).

30. Dalla-Torre, H. et al. Nucleotide Transformer: building and evaluating robust foundation models for human genomics. Nature Methods 1–11 (2024).

31. Chung, T. H., Zhuravskaya, A. & Makeyev, E. V. Regulation potential of transcribed simple repeated sequences in developing neurons. Hum. Genet. 143, 875–895 (2024).

32. Smit, AFA, Hubley, R & Green, P. RepeatMasker Open-4.0. http://www.repeatmasker.org (2013-2015).

33. Frith, M. C. A new repeat-masking method enables specific detection of homologous sequences. Nucleic Acids Res 39, e23 (2011).

34. Lu, J. Y. et al. Genomic Repeats Categorize Genes with Distinct Functions for Orchestrated Regulation. Cell Rep 30, 3296–3311.e5 (2020).

35. Van Nostrand, E. L. et al. Robust transcriptome-wide discovery of RNA-binding protein binding sites with enhanced CLIP (eCLIP). Nature methods 13, (2016).

36. Kim, Y. K. & Maquat, L. E. UPFront and center in RNA decay: UPF1 in nonsense-mediated mRNA decay and beyond. RNA (New York, N.Y.) 25, (2019).

37. Fischer, J. W., Busa, V. F., Shao, Y. & Leung, A. K. L. Structure-mediated RNA decay by UPF1 and G3BP1. Molecular cell 78, 70 (2020).

38. Chu, C. Y. & Rana, T. M. Translation repression in human cells by microRNA-induced gene silencing requires RCK/p54. PLoS biology 4, (2006).

39. Eulalio, A., Behm-Ansmant, I., Schweizer D & Izaurralde, E. P-body formation is a consequence, not the cause, of RNA-mediated gene silencing. Molecular and cellular biology 27, (2007).

40. Ayache, J. et al. P-body assembly requires DDX6 repression complexes rather than decay or Ataxin2/2L complexes. Molecular biology of the cell 26, (2015).

41. Biegel, J. M. et al. Cellular DEAD-box RNA helicase DDX6 modulates interaction of miR-122 with the 5’ untranslated region of hepatitis C virus RNA. Virology 507, (2017).

42. Ripin, N., de Vasconcelos L, M., Ugay, D. A. & Parker, R. DDX6 modulates P-body and stress granule assembly, composition, and docking. The Journal of cell biology 223, (2024).

43. Shih, C. Y., Chen, Y. C., Lin, H. Y. & Chu, C. Y. RNA Helicase DDX6 Regulates A-to-I Editing and Neuronal Differentiation in Human Cells. International journal of molecular sciences 24, (2023).

44. Van Nostrand, E. L. et al. A large-scale binding and functional map of human RNA-binding proteins. Nature 583, (2020).

45. Kim, Y. K., Furic, L., Desgroseillers, L. & Maquat, L. E. Mammalian Staufen1 recruits Upf1 to specific mRNA 3’UTRs so as to elicit mRNA decay. Cell 120, (2005).

46. Ricci, E. P. et al. Staufen1 senses overall transcript secondary structure to regulate translation. Nature structural & molecular biology 21, (2014).

47. Sugimoto, Y. et al. hiCLIP reveals the in vivo atlas of mRNA secondary structures recognized by Staufen 1. Nature 519, (2015).

48. Chakrabarti, A. M., Iosub, I. A., Lee, F. C. Y., Ule, J. & Luscombe, N. M. A computationally-enhanced hiCLIP atlas reveals Staufen1-RNA binding features and links 3’ UTR structure to RNA metabolism. Nucleic acids research 51, (2023).

49. Chen, C. et al. POU3F2 is a regulator of a gene coexpression network in brain tissue from patients with neuropsychiatric disorders. Science translational medicine 10, eaat8178 (2018).

50. Keegan, L. P., Leroy, A., Sproul, D. & O’Connell, M. A. Adenosine deaminases acting on RNA (ADARs): RNA-editing enzymes. Genome biology 5, (2004).

51. Heraud-Farlow, J. E. et al. Protein recoding by ADAR1-mediated RNA editing is not essential for normal development and homeostasis. Genome biology 18, (2017).

52. Chung, H. et al. Human ADAR1 Prevents Endogenous RNA from Triggering Translational Shutdown. Cell 172, (2018).

53. Hu, S. B. et al. ADAR1p150 prevents MDA5 and PKR activation via distinct mechanisms to avert fatal autoinflammation. Molecular cell 83, (2023).

54. Sharma, S. & Baysal, B. E. Stem-loop structure preference for site-specific RNA editing by APOBEC3A and APOBEC3G. PeerJ 5, e4136 (2017).

55. Pellegrini, F., et al. A KO mouse model for the lncRNA Lhx1os produces motor neuron alterations and locomotor impairment. iScience 26, (2022).

56. Mückstein, U. et al. Thermodynamics of RNA-RNA binding. Bioinformatics 22, 1177–1182 (2006).

57. Rehmsmeier, M., Steffen, P., Hochsmann, M. & Giegerich, R. Fast and effective prediction of microRNA/target duplexes. RNA 10, 1507–1517 (2004).

58. Antonov, I. V., Mazurov, E., Borodovsky, M. & Medvedeva, Y. A. Prediction of lncRNAs and their interactions with nucleic acids: benchmarking bioinformatics tools. Brief. Bioinform. 20, 551–564 (2019).

59. Kelley, D. R., Snoek, J. & Rinn, J. L. Basset: learning the regulatory code of the accessible genome with deep convolutional neural networks. Genome Res 26, 990–999 (2016).

60. Zhou, J. & Troyanskaya, O. G. Predicting effects of noncoding variants with deep learning-based sequence model. Nature methods 12, (2015).

61. Avsec, Ž., et al. Effective gene expression prediction from sequence by integrating long-range interactions. Nature Methods 18, 1196–1203 (2021).

62. Chen, J., et al. Interpretable RNA Foundation Model from Unannotated Data for Highly Accurate RNA Structure and Function Predictions. arXiv [q-bio.QM] (2022).

63. Wang, N. et al. Multi-purpose RNA language modelling with motif-aware pretraining and type-guided fine-tuning. Nature Machine Intelligence 6, 548–557 (2024).

64. Antonov, I., Marakhonov, A., Zamkova, M. & Medvedeva, Y. ASSA: Fast identification of statistically significant interactions between long RNAs. Journal of bioinformatics and computational biology 16, (2018).

65. Tafer, H. & Hofacker, I. L. RNAplex: a fast tool for RNA-RNA interaction search. Bioinformatics (Oxford, England) 24, (2008).

66. Alkan, F. et al. RIsearch2: suffix array-based large-scale prediction of RNA-RNA interactions and siRNA off-targets. Nucleic acids research 45, (2017).

67. Amatria-Barral, I., González-Domínguez, J. & Touriño, J. pRIblast: A highly efficient parallel application for comprehensive lncRNA–RNA interaction prediction. Future Generation Computer Systems 138, 270–279 (2023).

68. Mückstein, U. et al. Translational control by RNA-RNA interaction: Improved computation of RNA-RNA binding thermodynamics. in Communications in Computer and Information Science 114–127 (Springer Berlin Heidelberg, Berlin, Heidelberg, 2008).

69. Bernhart, S. H. et al. Partition function and base pairing probabilities of RNA heterodimers. Algorithms Mol. Biol. 1, 3 (2006).

70. Singh, J., Hanson, J., Paliwal, K. & Zhou, Y. RNA secondary structure prediction using an ensemble of two-dimensional deep neural networks and transfer learning. Nature communications 10, (2019).

71. Lang, M., Litfin, T., Chen, K., Zhan, J. & Zhou, Y. Benchmarking the methods for predicting base pairs in RNA-RNA interactions. Bioinformatics (Oxford, England) 41, (2025).

72. Selvaraju, R. R. et al. Grad-CAM: Visual explanations from deep networks via gradient-based localization. in 2017 IEEE International Conference on Computer Vision (ICCV) 618–626 (IEEE, 2017).

73. Farberov, S. et al. Structural features within the NORAD long noncoding RNA underlie efficient repression of Pumilio activity. Nat. Struct. Mol. Biol. (2024) doi:10.1038/s41594-024-01393-5.

74. Martone, J. et al. SMaRT lncRNA controls translation of a G-quadruplex-containing mRNA antagonizing the DHX36 helicase. EMBO Rep. 21, e49942 (2020).

75. Scalzitti, S. et al. Lnc-SMaRT translational regulation of Spire1, A new player in muscle differentiation. J. Mol. Biol. 434, 167384 (2022).

76. Rouskin, S., Zubradt, M., Washietl, S., Kellis, M. & Weissman, J. S. Genome-wide probing of RNA structure reveals active unfolding of mRNA structures in vivo. Nature 505, (2014).

77. Davidson, E. H. & Britten, R. J. Regulation of gene expression: possible role of repetitive sequences. Science 204, 1052–1059 (1979).

78. Trigiante, G., Blanes Ruiz, N. & Cerase, A. Emerging roles of repetitive and repeat-containing RNA in nuclear and chromatin organization and gene expression. Front. Cell Dev. Biol. 9, 735527 (2021).

79. Rué, L. et al. Targeting CAG repeat RNAs reduces Huntington’s disease phenotype independently of huntingtin levels. J. Clin. Invest. 126, 4319–4330 (2016).

80. Trost, B. et al. Genome-wide detection of tandem DNA repeats that are expanded in autism. Nature 586, 80–86 (2020).

81. DeJesus-Hernandez, M. et al. Expanded GGGGCC hexanucleotide repeat in noncoding region of C9ORF72 causes chromosome 9p-linked FTD and ALS. Neuron 72, 245–256 (2011).

82. Vandelli, A., Cid Samper, F., Torrent Burgas, M., Sanchez de Groot, N. & Tartaglia, G. G. The interplay between disordered regions in RNAs and proteins modulates interactions within Stress Granules and Processing Bodies. J. Mol. Biol. 434, 167159 (2022).

83. Wojciechowska, M. & Krzyzosiak, W. J. Cellular toxicity of expanded RNA repeats: focus on RNA foci. Hum. Mol. Genet. 20, 3811–3821 (2011).

84. Jain, A. & Vale, R. D. RNA phase transitions in repeat expansion disorders. Nature 546, 243–247 (2017).

85. Sanders, D. W. & Brangwynne, C. P. Neurodegenerative disease: RNA repeats put a freeze on cells. Nature 546, 215–216 (2017).

86. Fiorentino, J., Armaos, A., Colantoni, A. & Tartaglia, G. G. Prediction of protein-RNA interactions from single-cell transcriptomic data. Nucleic Acids Res. 52, e31 (2024).

87. Gong, J. et al. RISE: a database of RNA interactome from sequencing experiments. Nucleic acids research 46, (2018).

88. O’Leary, N. A. et al. Reference sequence (RefSeq) database at NCBI: current status, taxonomic expansion, and functional annotation. Nucleic Acids Res 44, D733–45 (2016).

89. Speir, M. L. et al. The UCSC Genome Browser database: 2016 update. Nucleic Acids Res 44, D717–25 (2016).

90. Yates, A. D. et al. Ensembl 2020. Nucleic acids research 48, (2020).

91. Quinlan, A. R. & Hall, I. M. BEDTools: a flexible suite of utilities for comparing genomic features. Bioinformatics (Oxford, England) 26, (2010).

92. Durinck, S., Spellman, P. T., Birney, E. & Huber, W. Mapping identifiers for the integration of genomic datasets with the R/Bioconductor package biomaRt. Nature protocols 4, (2009).

93. Abdennur, N. et al. Bioframe: operations on genomic intervals in Pandas dataframes. Bioinformatics (Oxford, England) 40, (2024).

94. Smedley, D. et al. BioMart--biological queries made easy. BMC genomics 10, (2009).

95. Liao, Y., Wang, J., Jaehnig, E. J., Shi, Z. & Zhang, B. WebGestalt 2019: gene set analysis toolkit with revamped UIs and APIs. Nucleic acids research 47, (2019).

96. Moore, J. E. et al. Expanded encyclopaedias of DNA elements in the human and mouse genomes. Nature 583, (2020).

97. Wilkins, O. G., Capitanchik, C., Luscombe, N. M. & Ule, J. Ultraplex: A rapid, flexible, all-in-one fastq demultiplexer. Wellcome open research 6, (2021).

98. Martin, M. Cutadapt removes adapter sequences from high-throughput sequencing reads. EMBnet.journal 17, 10–12 (2011).

99. Dobin, A. et al. STAR: ultrafast universal RNA-seq aligner. Bioinformatics 29, 15–21 (2013).

100. Smith, T., Heger, A. & Sudbery, I. UMI-tools: modeling sequencing errors in Unique Molecular Identifiers to improve quantification accuracy. Genome research 27, (2017).

101. Liao, Y., Smyth, G. K. & Shi, W. featureCounts: an efficient general purpose program for assigning sequence reads to genomic features. Bioinformatics (Oxford, England) 30, (2014).

102. Robinson, M. D., McCarthy, D. J. & Smyth, G. K. edgeR: a Bioconductor package for differential expression analysis of digital gene expression data. Bioinformatics 26, 139–140 (2010).

103. Clough, E. et al. NCBI GEO: archive for gene expression and epigenomics data sets: 23-year update. Nucleic Acids Res 52, D138–D144 (2024).

104. Gu, Z., Eils, R. & Schlesner, M. Complex heatmaps reveal patterns and correlations in multidimensional genomic data. Bioinformatics (Oxford, England) 32, (2016).

105. Hagberg, A. A., Schult, D. A. & Swart, P. J. Exploring Network Structure, Dynamics, and Function using NetworkX. scipy (2008) doi:10.25080/TCWV9851.

106. Zhang, B. et al. ADAR1 links R-loop homeostasis to ATR activation in replication stress response. Nucleic acids research 51, (2023).

107. Wulfridge, P. & Sarma, K. A nuclease- and bisulfite-based strategy captures strand-specific R-loops genome-wide. eLife 10, (2021).

108. Wichterle, H. & Peljto, M. Differentiation of mouse embryonic stem cells to spinal motor neurons. Curr. Protoc. Stem Cell Biol. Chapter 1, Unit 1H.1.1–1H.1.9 (2008).

109. Errichelli, L. et al. FUS affects circular RNA expression in murine embryonic stem cell-derived motor neurons. Nat. Commun. 8, 14741 (2017).

110. Sayers, E. W. et al. Database resources of the national center for biotechnology information. Nucleic Acids Res. 50, D20–D26 (2022).

111. Langmead, B. & Salzberg, S. L. Fast gapped-read alignment with Bowtie 2. Nature methods 9, 357 (2012).

112. Broad Institute. Picard Toolkit. Broad Institute, GitHub repository. https://broadinstitute.github.io/picard/ (2019).

113. Anders, S., Pyl, P. T. & Huber, W. HTSeq--a Python framework to work with high-throughput sequencing data. Bioinformatics 31, 166–169 (2015).

114. Patro, R., Duggal, G., Love, M. I., Irizarry, R. A. & Kingsford, C. Salmon provides fast and bias-aware quantification of transcript expression. Nature Methods 14, 417–419 (2017).

115. Fukunaga, T. & Hamada, M. RIblast: an ultrafast RNA-RNA interaction prediction system based on a seed-and-extension approach. Bioinformatics (Oxford, England) 33, (2017).

116. Pedregosa, F. et al. Scikit-learn: Machine Learning in Python. Journal of Machine Learning Research 12, 2825–2830 (2011).

117. Fu, L. et al. UFold: fast and accurate RNA secondary structure prediction with deep learning. Nucleic Acids Res. 50, e14 (2022).

118. Vaswani, A., et al. Attention Is All You Need. arXiv [cs.CL] (2017).

119. Egele, R., Mohr, F., Viering, T. & Balaprakash, P. The Unreasonable Effectiveness Of Early Discarding After One Epoch In Neural Network Hyperparameter Optimization. arXiv [cs.LG] (2024).

120. Paszke, A., et al. PyTorch: An imperative style, high-performance deep learning library. arXiv [cs.LG] (2019).

121. Singh, J. et al. Improved RNA secondary structure and tertiary base-pairing prediction using evolutionary profile, mutational coupling and two-dimensional transfer learning. Bioinformatics (Oxford, England) 37, (2021).

122. Li, P., Zhou, X., Xu, K. & Zhang, Q. C. RASP: an atlas of transcriptome-wide RNA secondary structure probing data. Nucleic acids research 49, (2021).

