## Supplementary Information for "Decoding RNA–RNA Interactions: The Role of Low-Complexity Repeats and a Deep Learning Framework for Sequence-Based Prediction"

### Supplementary Fig. 1

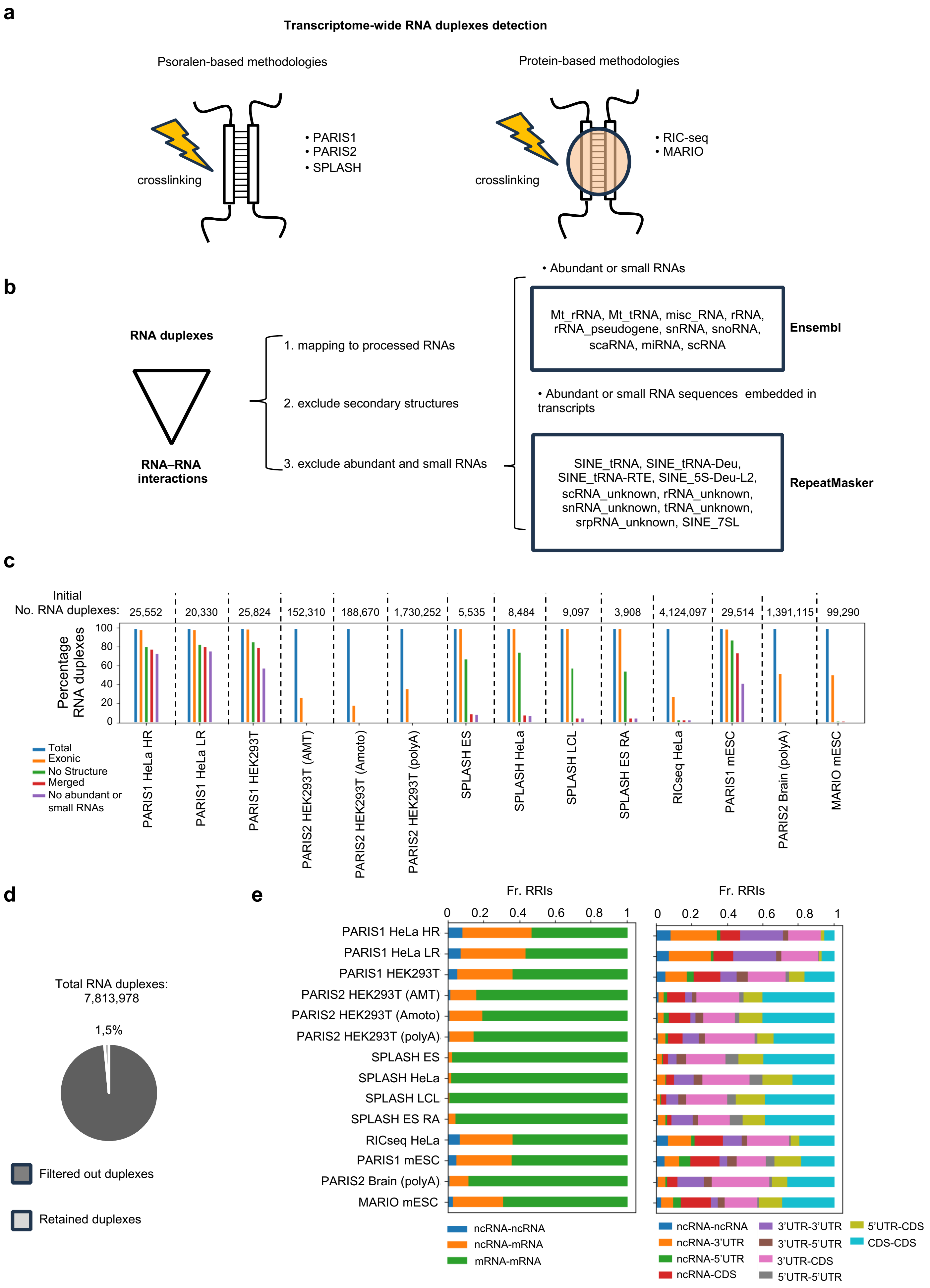

f

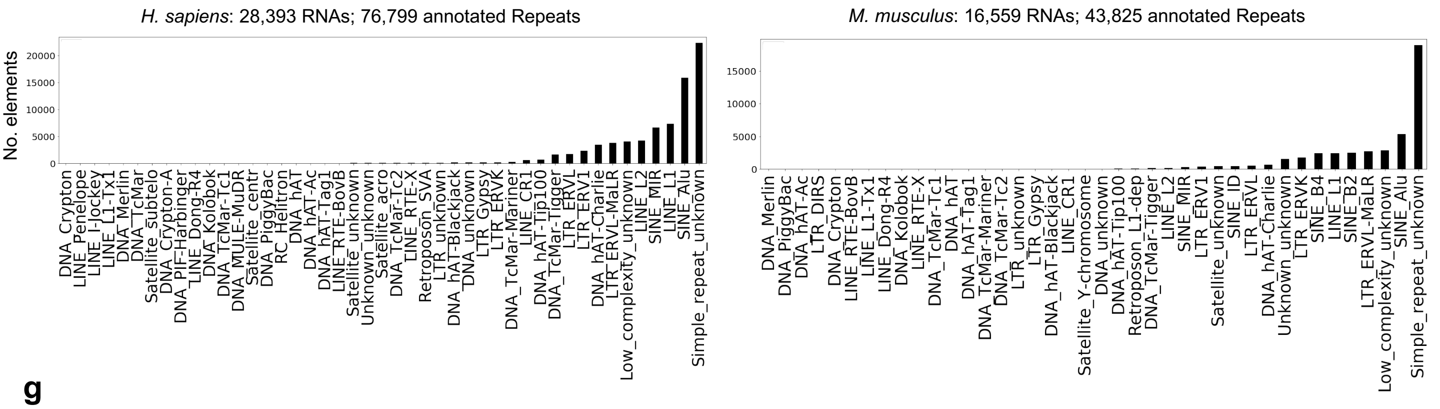
H. sapiens: 28,393 RNAs; 76,799 annotated Repeats
No. elements
0
5000
10000
15000
20000
DNA_Crypton
LINE_Penelope
LINE_I-Jockey
LINE_L1-Tx1
DNA_Merlin
DNA_TcMar
Satellite_subtelo
DNA_Crypton-A
DNA_PIF-Harbinger
LINE_Dong-R4
DNA_Kolobok
DNA_TcMar-Tc1
DNA_MULE-MuDR
Satellite_centr
DNA_PiggyBac
RC_Helitron
DNA_hAT
DNA_hAT-Ac
DNA_hAT-Tag1
LINE_RTE-BovB
Satellite_unknown
Unknown_unknown
Satellite_acro
DNA_TcMar-Tc2
LINE_RTE-X
Retroposon_SVA
LTR_unknown
DNA_hAT-Blackjack
DNA_unknown
LTR_Gypsy
LTR_ERVK
DNA_TcMar-Mariner
LINE_CR1
DNA_hAT-Tip100
DNA_TcMar-Tigger
LTR_ERVL
LTR_ERV1
DNA_hAT-Charlie
LTR_ERVL-MaLR
Low_complexity_unknown
LINE_L2
SINE_MIR
LINE_L1
SINE_Alu
Simple_repeat_unknown
M. musculus: 16,559 RNAs; 43,825 annotated Repeats
0
5000
10000
15000
DNA_Merlin
DNA_PiggyBac
DNA_hAT-Ac
LTR_DIRS
DNA_Crypton
LINE_RTE-BovB
LINE_L1-Tx1
LINE_Dong-R4
DNA_Kolobok
LINE_RTE-X
DNA_TcMar-Tc1
DNA_hAT
DNA_hAT-Tag1
DNA_TcMar-Mariner
DNA_TcMar-Tc2
LTR_unknown
LTR_Gypsy
DNA_hAT-Blackjack
LINE_CR1
Satellite_Y-chromosome
DNA_unknown
DNA_hAT-Tip100
Retroposon_L1-dep
DNA_TcMar-Tigger
LINE_L2
SINE_MIR
LTR_ERV1
SINE_ID
Satellite_unknown
LTR_ERVL
DNA_hAT-Charlie
Unknown_unknown
LTR_ERVK
SINE_B4
LINE_L1
SINE_B2
LTR_ERVL-MaLR
Low_complexity_unknown
SINE_Alu
Simple_repeat_unknown

g

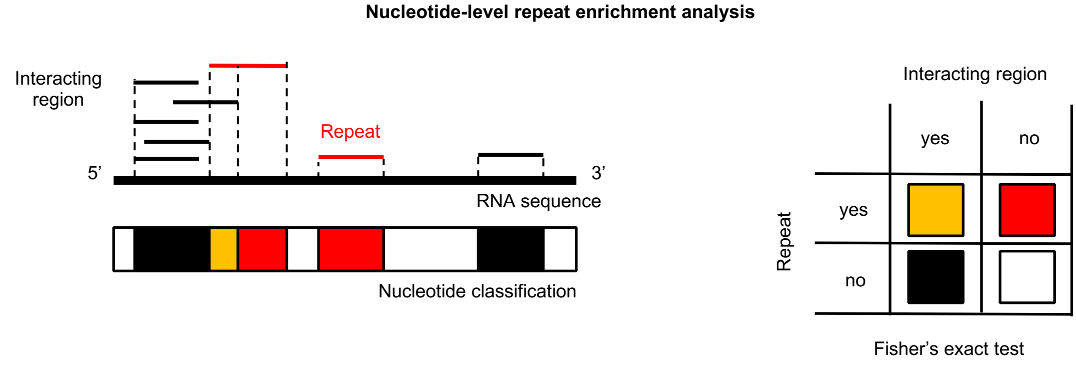
Nucleotide-level repeat enrichment analysis
Interacting region
Repeat
5'
3'
RNA sequence
Nucleotide classification
Interacting region
yes
no
Repeat
yes
no
Fisher's exact test

h

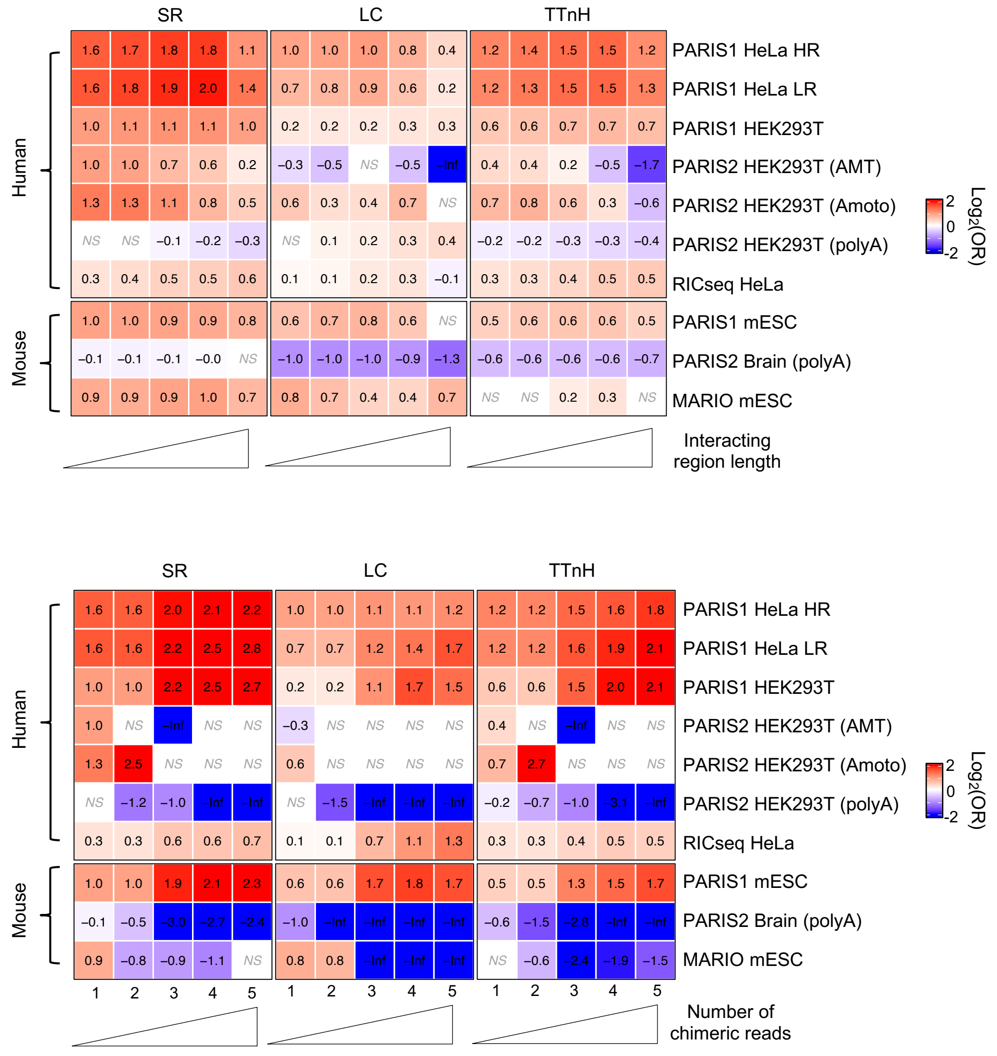
SR
LC
TTnH
Human
Mouse
1.6 1.7 1.8 1.8 1.1 | 1.0 1.0 1.0 0.8 0.4 | 1.2 1.4 1.5 1.5 1.2 PARIS1 HeLa HR
1.6 1.8 1.9 2.0 1.4 | 0.7 0.8 0.9 0.6 0.2 | 1.2 1.3 1.5 1.5 1.3 PARIS1 HeLa LR
1.0 1.1 1.1 1.1 1.0 | 0.2 0.2 0.2 0.3 0.3 | 0.6 0.6 0.7 0.7 0.7 PARIS1 HEK293T
1.0 1.0 0.7 0.6 0.2 | −0.3 −0.5 NS −0.5 −Inf | 0.4 0.4 0.2 −0.5 −1.7 PARIS2 HEK293T (AMT)
1.3 1.3 1.1 0.8 0.5 | 0.6 0.3 0.4 0.7 NS | 0.7 0.8 0.6 0.3 −0.6 PARIS2 HEK293T (Amoto)
NS NS −0.1 −0.2 −0.3 | NS 0.1 0.2 0.3 0.4 | −0.2 −0.2 −0.3 −0.3 −0.4 PARIS2 HEK293T (polyA)
0.3 0.4 0.5 0.5 0.6 | 0.1 0.1 0.2 0.3 −0.1 | 0.3 0.3 0.4 0.5 0.5 RICseq HeLa
1.0 1.0 0.9 0.9 0.8 | 0.6 0.7 0.8 0.6 NS | 0.5 0.6 0.6 0.6 0.5 PARIS1 mESC
−0.1 −0.1 −0.1 −0.0 NS | −1.0 −1.0 −1.0 −0.9 −1.3 | −0.6 −0.6 −0.6 −0.6 −0.7 PARIS2 Brain (polyA)
0.9 0.9 0.9 1.0 0.7 | 0.8 0.7 0.4 0.4 0.7 | NS NS 0.2 0.3 NS MARIO mESC
2
0
-2
Log2(OR)
Interacting region length
SR
LC
TTnH
Human
Mouse
1.6 1.6 2.0 2.1 2.2 | 1.0 1.0 1.1 1.1 1.2 | 1.2 1.2 1.5 1.6 1.8 PARIS1 HeLa HR
1.6 1.6 2.2 2.5 2.8 | 0.7 0.7 1.2 1.4 1.7 | 1.2 1.2 1.6 1.9 2.1 PARIS1 HeLa LR
1.0 1.0 2.2 2.5 2.7 | 0.2 0.2 1.1 1.7 1.5 | 0.6 0.6 1.5 2.0 2.1 PARIS1 HEK293T
1.0 NS −Inf NS NS | −0.3 NS NS NS NS | 0.4 NS −Inf NS NS PARIS2 HEK293T (AMT)
1.3 2.5 NS NS NS | 0.6 NS NS NS NS | 0.7 2.7 NS NS NS PARIS2 HEK293T (Amoto)
NS −1.2 −1.0 −Inf −Inf | NS −1.5 −Inf −Inf −Inf | −0.2 −0.7 −1.0 −3.1 −Inf PARIS2 HEK293T (polyA)
0.3 0.3 0.6 0.6 0.7 | 0.1 0.1 0.7 1.1 1.3 | 0.3 0.3 0.4 0.5 0.5 RICseq HeLa
1.0 1.0 1.9 2.1 2.3 | 0.6 0.6 1.7 1.8 1.7 | 0.5 0.5 1.3 1.5 1.7 PARIS1 mESC
−0.1 −0.5 −3.0 −2.7 −2.4 | −1.0 −Inf −Inf −Inf −Inf | −0.6 −1.5 −2.8 −Inf −Inf PARIS2 Brain (polyA)
0.9 −0.8 −0.9 −1.1 NS | 0.8 0.8 −Inf −Inf −Inf | NS −0.6 −2.4 −1.9 −1.5 MARIO mESC
1 2 3 4 5 | 1 2 3 4 5 | 1 2 3 4 5
2
0
-2
Log2(OR)
Number of chimeric reads

i

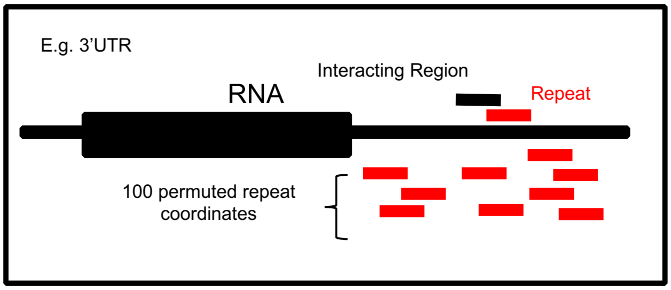

E.g. 3'UTR
Interacting Region
RNA
Repeat
100 permuted repeat coordinates


j

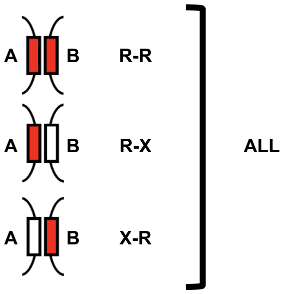

A B R-R
A B R-X
A B X-R
ALL


k

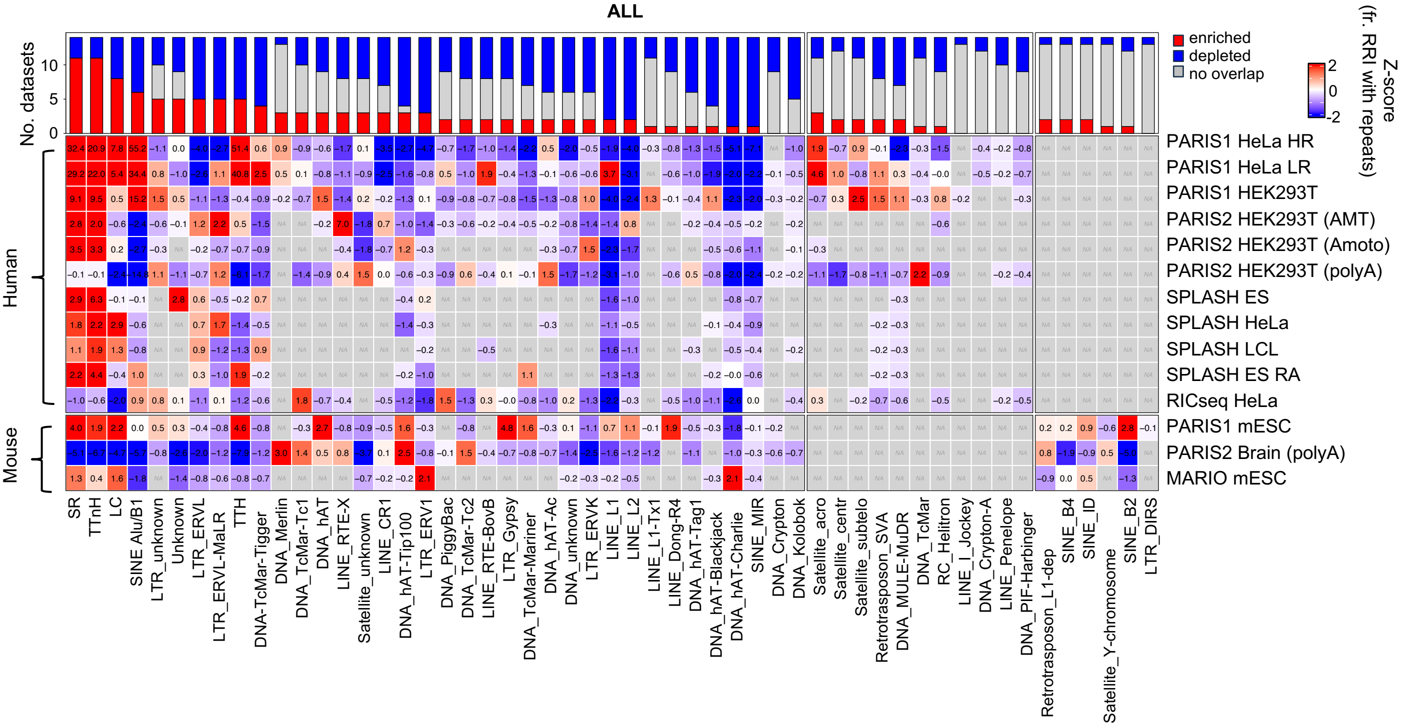

ALL
No. datasets
enriched
depleted
no overlap
Z-score (fr. RRI with repeats)
Human
Mouse
PARIS1 HeLa HR
PARIS1 HeLa LR
PARIS1 HEK293T
PARIS2 HEK293T (AMT)
PARIS2 HEK293T (Amoto)
PARIS2 HEK293T (polyA)
SPLASH ES
SPLASH HeLa
SPLASH LCL
SPLASH ES RA
RICseq HeLa
PARIS1 mESC
PARIS2 Brain (polyA)
MARIO mESC
SR
TTnH
LC
SINE Alu/B1
LTR_unknown
Unknown
LTR_ERVL
LTR_ERVL-MaLR
TTH
DNA-TcMar-Tigger
DNA_Merlin
DNA_TcMar-Tc1
DNA_hAT
LINE_RTE-X
Satellite_unknown
LINE_CR1
DNA_hAT-Tip100
LTR_ERV1
DNA_PiggyBac
DNA_TcMar-Tc2
LINE_RTE-BovB
LTR_Gypsy
DNA_TcMar-Mariner
DNA_hAT-Ac
DNA_unknown
LTR_ERVK
LINE_L1
LINE_L2
LINE_L1-Tx1
LINE_Dong-R4
DNA_hAT-Tag1
DNA_hAT-Blackjack
DNA_hAT-Charlie
SINE_MIR
DNA_Crypton
DNA_Kolobok
Satellite_acro
Satellite_centr
Satellite_subtelo
Retrotrasposon_SVA
DNA_MULE-MuDR
DNA_TcMar
RC_Helitron
LINE_I_Jockey
DNA_Crypton-A
LINE_Penelope
DNA_PIF-Harbinger
Retrotrasposon_L1-dep
SINE_B4
SINE_ID
Satellite_Y-chromosome
SINE_B2
LTR_DIRS


l

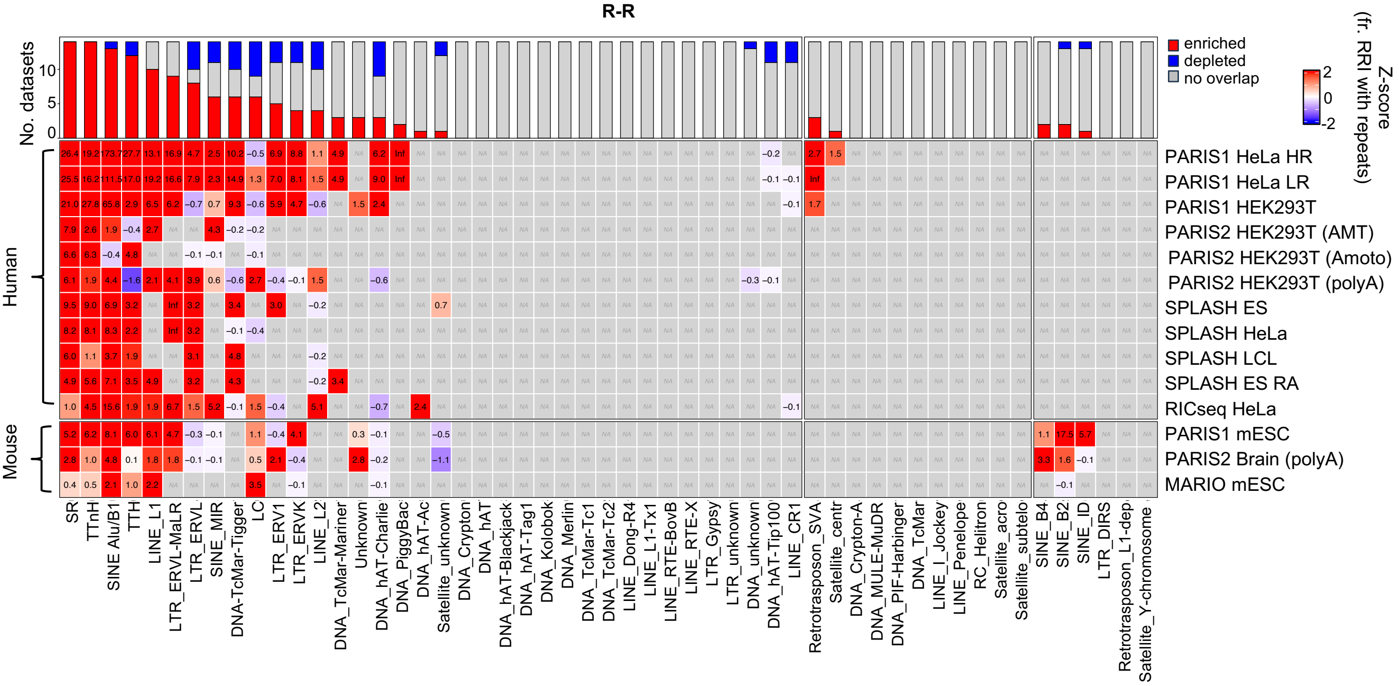

R-R
No. datasets
enriched
depleted
no overlap
Z-score (fr. RRI with repeats)
Human
Mouse
PARIS1 HeLa HR
PARIS1 HeLa LR
PARIS1 HEK293T
PARIS2 HEK293T (AMT)
PARIS2 HEK293T (Amoto)
PARIS2 HEK293T (polyA)
SPLASH ES
SPLASH HeLa
SPLASH LCL
SPLASH ES RA
RICseq HeLa
PARIS1 mESC
PARIS2 Brain (polyA)
MARIO mESC
SR
TTnH
SINE Alu/B1
TTH
LINE_L1
LTR_ERVL-MaLR
LTR_ERVL
SINE_MIR
DNA-TcMar-Tigger
LC
LTR_ERV1
LTR_ERVK
LINE_L2
DNA_TcMar-Mariner
Unknown
DNA_hAT-Charlie
DNA_PiggyBac
DNA_hAT-Ac
Satellite_unknown
DNA_Crypton
DNA_hAT
DNA_hAT-Blackjack
DNA_hAT-Tag1
DNA_Kolobok
DNA_Merlin
DNA_TcMar-Tc1
DNA_TcMar-Tc2
LINE_Dong-R4
LINE_L1-Tx1
LINE_RTE-BovB
LINE_RTE-X
LTR_Gypsy
LTR_unknown
DNA_unknown
DNA_hAT-Tip100
LINE_CR1
Retrotrasposon_SVA
Satellite_centr
DNA_Crypton-A
DNA_MULE-MuDR
DNA_PIF-Harbinger
DNA_TcMar
LINE_I_Jockey
LINE_Penelope
RC_Helitron
Satellite_acro
Satellite_subtelo
SINE_B4
SINE_B2
SINE_ID
LTR_DIRS
Retrotrasposon_L1-dep
Satellite_Y-chromosome

**m**

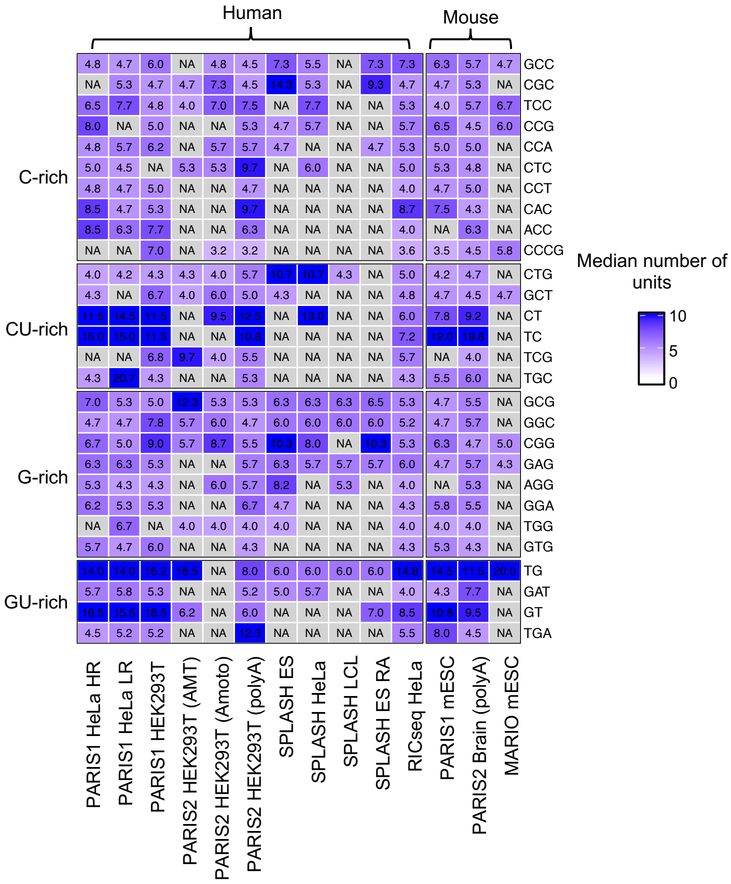
Human
Mouse
C-rich
CU-rich
G-rich
GU-rich
4.8 4.7 6.0 NA 4.8 4.5 7.3 5.5 NA 7.3 7.3 6.3 5.7 4.7 GCC
NA 5.3 4.7 4.7 7.3 4.5 14.3 5.3 NA 9.3 4.7 4.7 5.3 NA CGC
6.5 7.7 4.8 4.0 7.0 7.5 NA 7.7 NA NA 5.3 4.0 5.7 6.7 TCC
8.0 NA 5.0 NA NA 5.3 4.7 5.7 NA NA 5.7 6.5 4.5 6.0 CCG
4.8 5.7 6.2 NA 5.7 5.7 4.7 NA NA 4.7 5.3 5.0 5.0 NA CCA
5.0 4.5 NA 5.3 5.3 9.7 NA 6.0 NA NA 5.0 5.3 4.8 NA CTC
4.8 4.7 5.0 NA NA 4.7 NA NA NA NA 4.0 4.7 5.0 NA CCT
8.5 4.7 5.3 NA NA 9.7 NA NA NA NA 8.7 7.5 4.3 NA CAC
8.5 6.3 7.7 NA NA 6.3 NA NA NA NA 4.0 NA 6.3 NA ACC
NA NA 7.0 NA 3.2 3.2 NA NA NA NA 3.6 3.5 4.5 5.8 CCCG
4.0 4.2 4.3 4.3 4.0 5.7 10.7 10.7 4.3 NA 5.0 4.2 4.7 NA CTG
4.3 NA 6.7 4.0 6.0 5.0 4.3 NA NA NA 4.8 4.7 4.5 4.7 GCT
11.5 14.5 11.5 NA 9.5 12.5 NA 13.0 NA NA 6.0 7.8 9.2 NA CT
15.0 15.0 11.5 NA NA 10.8 NA NA NA NA 7.2 12.0 19.8 NA TC
NA NA 6.8 9.7 4.0 5.5 NA NA NA NA 5.7 NA 4.0 NA TCG
4.3 20.7 4.3 NA NA 5.3 NA NA NA NA 4.3 5.5 6.0 NA TGC
7.0 5.3 5.0 12.3 5.3 5.3 6.3 6.3 6.3 6.5 5.3 4.7 5.5 NA GCG
4.7 4.7 7.8 5.7 6.0 4.7 6.0 6.0 6.0 6.0 5.2 4.7 5.7 NA GGC
6.7 5.0 9.0 5.7 8.7 5.5 10.3 8.0 NA 10.3 5.3 6.3 4.7 5.0 CGG
6.3 6.3 5.3 NA NA 5.7 6.3 5.7 5.7 5.7 6.0 4.7 5.7 4.3 GAG
5.3 4.3 4.3 NA 6.0 5.7 8.2 NA 5.3 NA 4.0 NA 5.3 NA AGG
6.2 5.3 5.3 NA NA 6.7 4.7 NA NA NA 4.3 5.8 5.5 NA GGA
NA 6.7 NA 4.0 4.0 4.0 4.0 NA NA NA 4.0 4.0 4.0 NA TGG
5.7 4.7 6.0 NA NA 4.3 NA NA NA NA 4.3 5.3 4.3 NA GTG
14.0 14.0 16.2 16.5 NA 8.0 6.0 6.0 6.0 6.0 14.8 14.5 11.5 20.0 TG
5.7 5.8 5.3 NA NA 5.2 5.0 5.7 NA NA 4.0 4.3 7.7 NA GAT
16.5 15.5 18.5 6.2 NA 6.0 NA NA NA 7.0 8.5 10.8 9.5 NA GT
4.5 5.2 5.2 NA NA 12.3 NA NA NA NA 5.5 8.0 4.5 NA TGA
PARIS1 HeLa HR
PARIS1 HeLa LR
PARIS1 HEK293T
PARIS2 HEK293T (AMT)
PARIS2 HEK293T (Amoto)
PARIS2 HEK293T (polyA)
SPLASH ES
SPLASH HeLa
SPLASH LCL
SPLASH ES RA
RICseq HeLa
PARIS1 mESC
PARIS2 Brain (polyA)
MARIO mESC
Median number of units
10
5
0

**n**

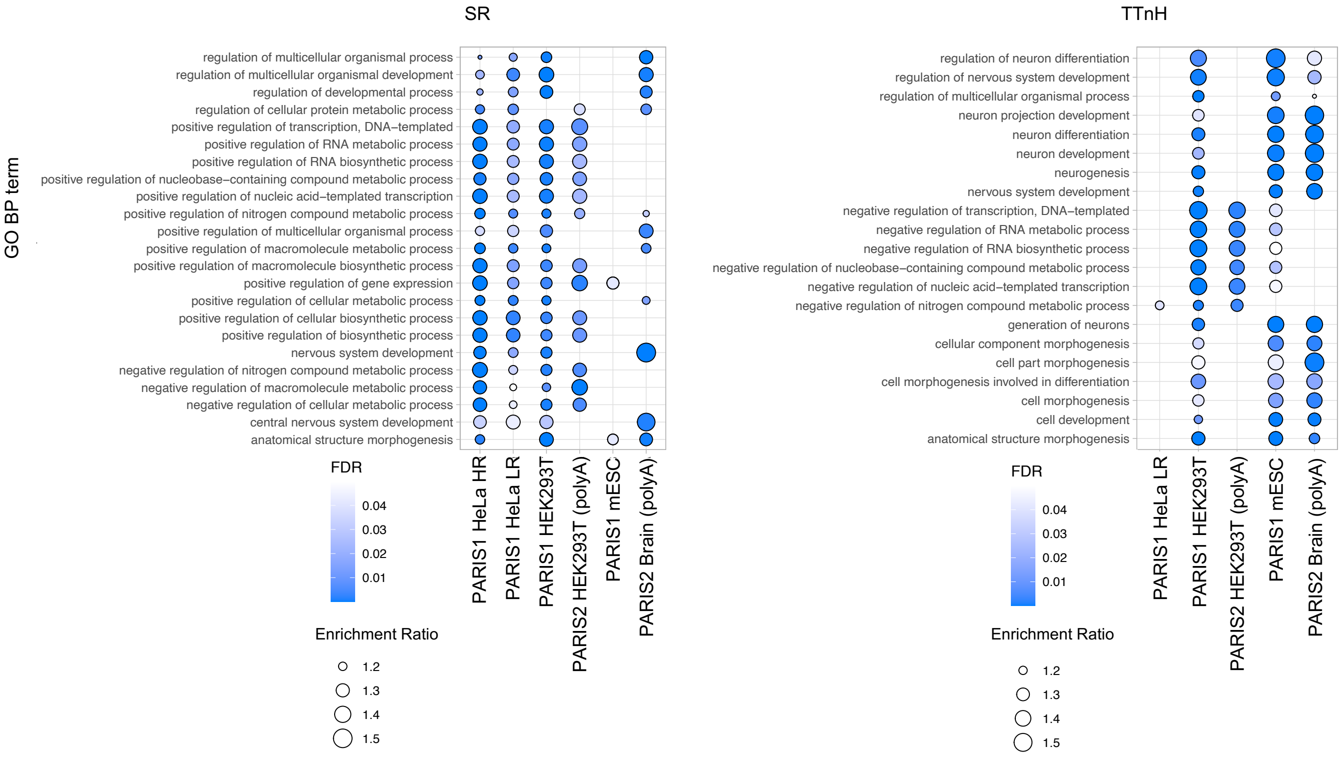
SR
TTnH
GO BP term
regulation of multicellular organismal process
regulation of multicellular organismal development
regulation of developmental process
regulation of cellular protein metabolic process
positive regulation of transcription, DNA–templated
positive regulation of RNA metabolic process
positive regulation of RNA biosynthetic process
positive regulation of nucleobase–containing compound metabolic process
positive regulation of nucleic acid–templated transcription
positive regulation of nitrogen compound metabolic process
positive regulation of multicellular organismal process
positive regulation of macromolecule metabolic process
positive regulation of macromolecule biosynthetic process
positive regulation of gene expression
positive regulation of cellular metabolic process
positive regulation of cellular biosynthetic process
positive regulation of biosynthetic process
nervous system development
negative regulation of nitrogen compound metabolic process
negative regulation of macromolecule metabolic process
negative regulation of cellular metabolic process
central nervous system development
anatomical structure morphogenesis
PARIS1 HeLa HR
PARIS1 HeLa LR
PARIS1 HEK293T
PARIS2 HEK293T (polyA)
PARIS1 mESC
PARIS2 Brain (polyA)
FDR
0.04
0.03
0.02
0.01
Enrichment Ratio
1.2
1.3
1.4
1.5
regulation of neuron differentiation
regulation of nervous system development
regulation of multicellular organismal process
neuron projection development
neuron differentiation
neuron development
neurogenesis
nervous system development
negative regulation of transcription, DNA–templated
negative regulation of RNA metabolic process
negative regulation of RNA biosynthetic process
negative regulation of nucleobase–containing compound metabolic process
negative regulation of nucleic acid–templated transcription
negative regulation of nitrogen compound metabolic process
generation of neurons
cellular component morphogenesis
cell part morphogenesis
cell morphogenesis involved in differentiation
cell morphogenesis
cell development
anatomical structure morphogenesis
PARIS1 HeLa LR
PARIS1 HEK293T
PARIS2 HEK293T (polyA)
PARIS1 mESC
PARIS2 Brain (polyA)
FDR
0.04
0.03
0.02
0.01
Enrichment Ratio
1.2
1.3
1.4
1.5

**Supplementary Fig. 1. Low-complexity repeats are key contributors to RNA–RNA contacts. a**, Schematic representation of the transcriptome-wide RNA duplex detection methodologies previously applied to generate the data analyzed in this study. **b**, Description of the filtering procedure employed to select the RRI interactions from the transcriptome-wide studies. The black boxes on the right list the Ensembl RNAs biotypes (top) and the RepeatMasker RNA repeats (bottom) used to filter out interactions mediated by highly abundant or small RNAs. **c**, Bar plot illustrating, for each transcriptome-wide experiment (x-axis), the percentage of RNA heteroduplexes selected in each step of the processing procedure (y-axis). The starting amount of RRIs from each experiment is indicated at the top of the bar plot. **d**, Pie chart reporting the percentage of RRIs selected after the processing procedure with respect to the total amount of RNA heteroduplexes collected from the transcriptome-wide studies. **e**, Bar plot depicting the distribution of RRIs based on two classifications: the biotypes of the interacting RNAs (left) and the RNA functional regions where the interacting patches are located (5'UTR, CDS, 3'UTR, or ncRNA, right). **f**, Bar plots summarizing the number of repetitive elements identified using RepeatMasker within the human (left panel) and murine (right panel) transcripts involved in the selected RRIs. The total number of transcripts and annotated repeats are indicated at the top of each bar plot. **g**, Schematic representation of the nucleotide-level repeat enrichment analysis. The diagram on the left illustrates an example transcript, highlighting the nucleotide classification used to assess co-occurrence of repeats and RRIs across the transcriptome-wide experiments. The panel on the right presents the contingency table employed to determine statistical significance for each repeat family using two-sided Fisher's exact test. **h**, Heatmaps illustrating the results of the nucleotide-level repeat enrichment analysis performed for LCRs on RRIs with progressively higher quality. Each row represents a transcriptome-wide experiment; rows are grouped by species. Columns, which represent RRI sets with progressively higher minimum quality (as schematically represented by the triangular diagrams at the bottom of the heatmaps), are grouped based on the tested LCR type. Quality is defined as the minimum number of chimeric reads supporting the interaction (top panel) or the minimum length of the interacting region, defined dynamically for each RRI set (lower panel, see **Methods**). $Log_2$(Odds Ratio) values are shown in cells corresponding to significant repeat enrichment or depletion (two-sided Fisher's exact test p-value adjusted with Benjamini-Hochberg method $< 0.05$ ). The color gradient ranges from blue (depletion) to red (enrichment), with the intensity of the color reflecting the strength of enrichment or depletion. Non-significant results are indicated with NS. The SPLASH dataset was excluded from these analyses because it contains few interactions supported by a minimum of four reads, and because it uses bins of standard length (100 nucleotides) to define interacting regions. **i**, Diagram illustrating the permutation-based repeat enrichment analysis. The thicker region within the schematized RNA molecule corresponds to the CDS. At the top, an example interacting region (black line) overlapping a repeated element (red line) within the 3'UTR is shown. Below, the red lines indicate 100 random permutations of the repeated element position within the 3' UTR region. By comparing the observed overlap to a null distribution derived from the permutations, it is possible to calculate a Z-score expressing the enrichment or depletion of the repeat family with respect to the RRIs. **j**, Diagram illustrating the possible configurations of an RNA–RNA interaction overlapping a repeated element: R-X (repeat on interacting patch A), X-R (repeat on interacting patch B), or R-R (repeat on both interacting patches). **k** and **l**, Heatmaps reporting, for each transcriptome-wide experiment (rows), the Z-scores of the permutation-based repeat enrichment analysis, performed considering ALL (**k**) or only R-R (**l**) configurations for each repeat family (columns). Enrichment or depletion of repeats in RRIs is indicated by red or blue colors, respectively; gray denotes the absence of overlap between RRIs and repeats (NA). The bar plot above the heatmap shows, for each tested repeat family, the number of transcriptome-wide experiments where Z-scores are positive (red), negative (blue), or not available (gray). **m**, Heatmap depicting the median number of repeated units for TTnH repeats that overlap RNA–RNA interactions from different transcriptome-wide experiments (rows). TTnH repeats are grouped by nucleotide categories (GU-rich, G-rich, CU-rich, and C-rich), while the experiments are grouped by species (human and mouse). Only the most common TTnH repeats with at least one overlap in at least seven datasets were considered. The intensity of the blue gradient indicates the number of repeated units for each motif, ranging from white (0) to dark blue ($\geq 10$). Grey cells labeled as NA represent cases where no overlap was observed. **n**, Dot plots illustrating the Gene Ontology (GO)

Biological Process (BP) terms enriched for RNAs involved in RRIs through SR (left panel) and TTnH (right panel). The x-axis lists transcriptome-wide experiments, while the y-axis shows significantly enriched GO BPs (FDR < 0.05); only categories enriched for at least four experiments were plotted. Each dot corresponds to an enriched GO term, with its size indicating the enrichment ratio (larger dots signify higher enrichment scores). Dot color reflects the FDR-adjusted p-value, with darker blue denoting greater statistical significance (lower FDR values).

### Supplementary Fig. 2

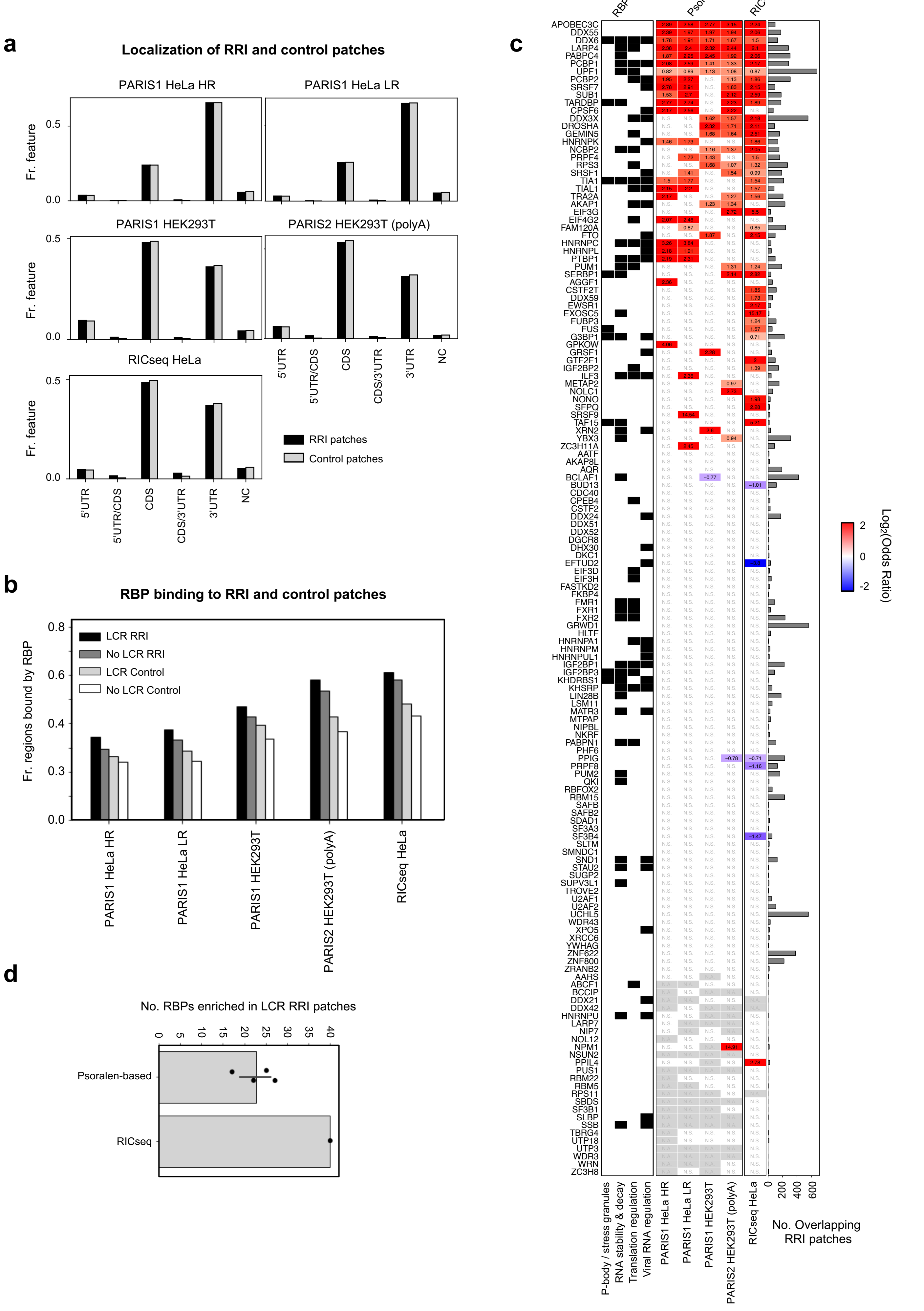

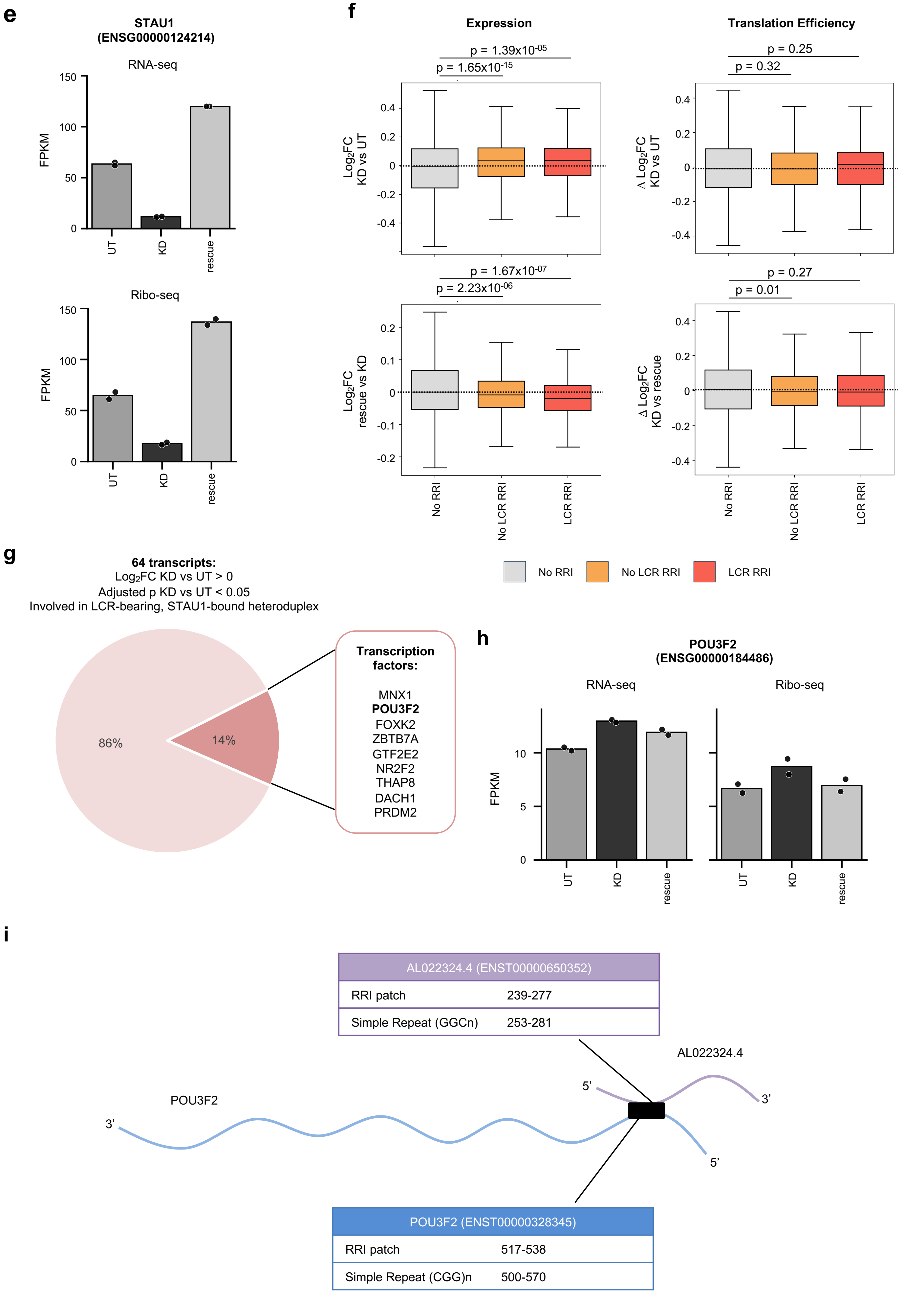
e
STAU1 (ENSG00000124214)
RNA-seq
FPKM
150
100
50
0
UT
KD
rescue
Ribo-seq
FPKM
150
100
50
0
UT
KD
rescue
f
Expression
p = 1.39x10-05
p = 1.65x10-15
Log2FC KD vs UT
0.4
0.2
0
-0.2
-0.4
p = 1.67x10-07
p = 2.23x10-06
Log2FC rescue vs KD
0.2
0.1
0
-0.1
-0.2
No RRI
No LCR RRI
LCR RRI
Translation Efficiency
p = 0.25
p = 0.32
Δ Log2FC KD vs UT
0.4
0.2
0
-0.2
-0.4
p = 0.27
p = 0.01
Δ Log2FC KD vs rescue
0.4
0.2
0
-0.2
-0.4
No RRI
No LCR RRI
LCR RRI
No RRI
No LCR RRI
LCR RRI
g
64 transcripts:
Log2FC KD vs UT > 0
Adjusted p KD vs UT < 0.05
Involved in LCR-bearing, STAU1-bound heteroduplex
86%
14%
Transcription factors:
MNX1
POU3F2
FOXK2
ZBTB7A
GTF2E2
NR2F2
THAP8
DACH1
PRDM2
h
POU3F2 (ENSG00000184486)
RNA-seq
Ribo-seq
FPKM
10
5
0
UT
KD
rescue
UT
KD
rescue
i
AL022324.4 (ENST00000650352)
RRI patch 239-277
Simple Repeat (GGCn) 253-281
AL022324.4
5'
3'
POU3F2
3'
5'
POU3F2 (ENST00000328345)
RRI patch 517-538
Simple Repeat (CGG)n 500-570

**Supplementary Fig. 2. LCR-bearing RRI patches colocalize with target sites of proteins regulating translation and decay. a**, Bar plots reporting, for each human RRI transcriptome-wide experiment examined for RBP binding, the fraction of RRI and control patches found in different transcript regions (5′UTR, 5′UTR/CDS boundary, CDS, CDS/3′UTR boundary, 3′UTR, non-coding RNA). Only transcripts expressed in HepG2 and K562 cells were used for the analysis. Black bars, RRI patches; grey bars, matched controls sampled from non-interacting regions of the same transcriptome. **b**, Bar plots showing, for each of the aforementioned RRI datasets (x-axis), the fraction of RRI and control patches that overlap with at least one eCLIP-derived RBP target site (y-axis). Patches are further classified based on whether they overlap with a low-complexity repeat (LCR) or not (No LCR). **c**, Heatmap showing, for each RBP present in the ENCODE eCLIP compendium (rows) and each RRI dataset, the $\log_2$(Odds Ratio) for RBP binding enrichment on RRI patches relative to matched control regions. Numbers inside cells report the corresponding $\log_2$(OR). Statistical significance was assessed with two-sided Fisher's exact test and Bonferroni correction; only cells with adjusted p-value $< 0.05$ are colored, with red indicating enrichment and blue indicating depletion. The left annotation marks RBP functional annotation. The right bar plot summarizes, for each RBP, the total number of RRI patches overlapping with its target sites across all evaluated datasets. **d**, Bar plot reporting the number of RBPs whose target sites significantly colocalize with RRI patches compared with matched control regions, grouped by RRI detection method (Psoralen-based and RIC-seq). Black dots denote the counts for individual datasets within each group. Bars represent mean values ± s.d.. **e**, Bar plot showing STAU1 expression quantified as FPKM values from RNA-seq (top panel) and Ribo-seq (bottom panel) data generated by Sugimoto et al. under STAU1 UT, KD, and rescue conditions. Bars represent mean expression, and individual data points for each replicate are overlaid. **f**, Box plots showing $\log_2$FC values calculated from RNA-seq data (left) and changes in translation efficience measured as $\Delta\log2$FC from Ribo-seq and RNA-seq data (right) for three transcript groups: transcripts not involved in RRI (No RRI), transcripts involved in RRI with no LCR (No LCR RRI) and transcripts involved in RRI with LCR (LCR RRI). Top panel displays KD vs UT comparison, while bottom panel displays rescue vs KD comparison. The dotted line at 0 marks no change. Boxes show the interquartile range (IQR), with the line indicating the median; whiskers extend to the most extreme data points within 1.5×IQR from the median; outliers were not included. Differences between distributions were assessed with two-tailed Mann-Whitney U-test, with p-values corrected using the Benjamini–Hochberg method. **g**, Pie chart summarizing 64 transcripts that are upregulated in KD vs UT comparison ($\log_2$FC $> 0$; adjusted p-value $< 0.05$) and are involved in LCR-bearing, STAU1-bound heteroduplexes. The darkest sector denotes genes coding for transcription factors, which are listed on the right. **h**, Bar plot showing POU3F2 expression quantified as FPKM values from RNA-seq (top panel) and Ribo-seq (bottom panel) data generated by Sugimoto et al. under STAU1 UT, KD, and rescue conditions. Bars represent mean expression, and individual data points for each replicate are overlaid. **i**, Schematic of the intermolecular pairing site between POU3F2 mRNA and AL022324.4 lncRNA. The reported transcript IDs refer to the longest functional isoform annotated Ensembl 99. The black box marks the identified duplex locus; tables report, for each RNA, the transcript coordinates of the RRI patch and of the overlapping LCR region.

### Supplementary Fig. 3

**a**

**Co-occurrence of LCRs and RRIs along the meta-transcript**

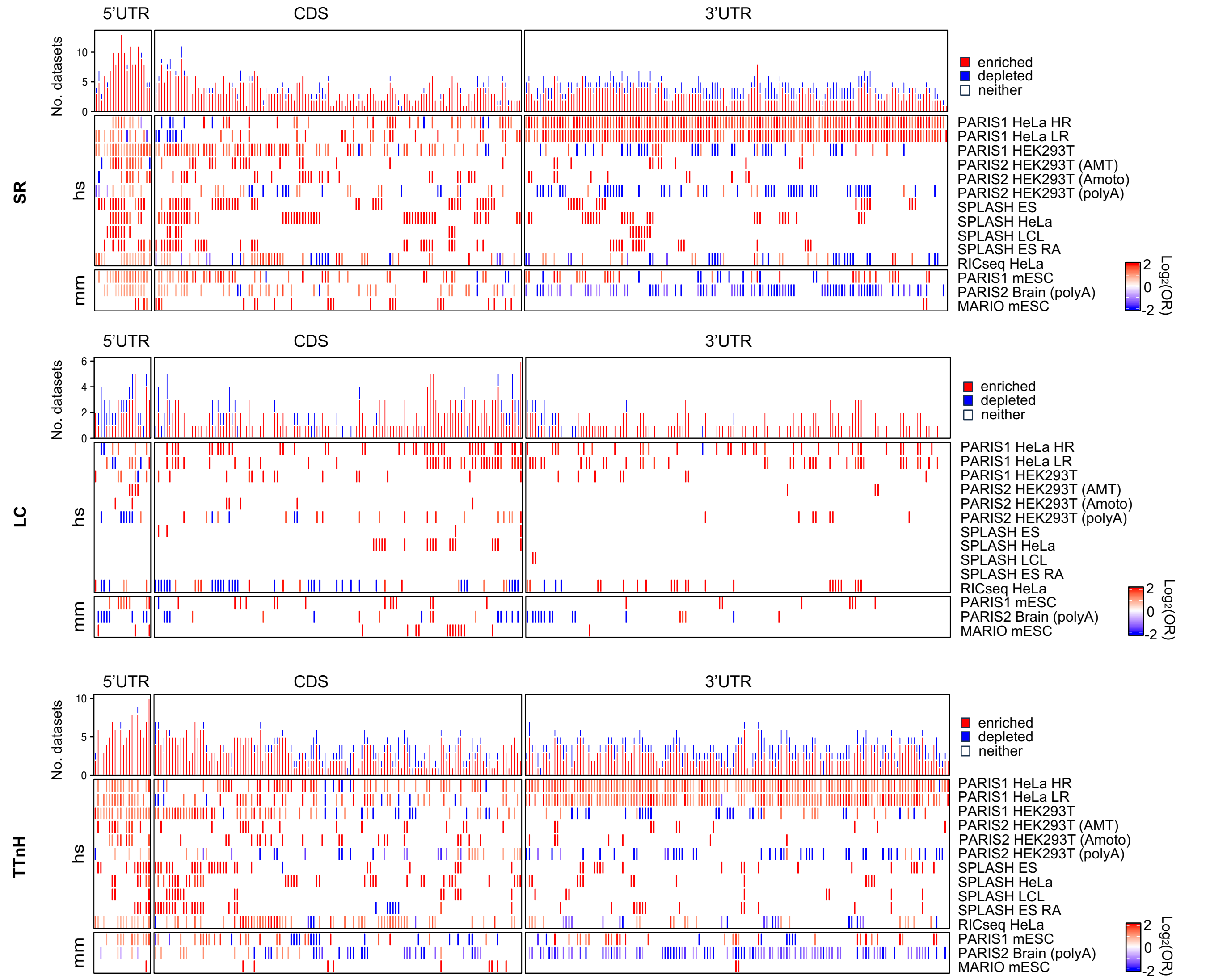


**b**

**Distance of LCR-bearing RRIs from the network center compared with that of control RRIs**

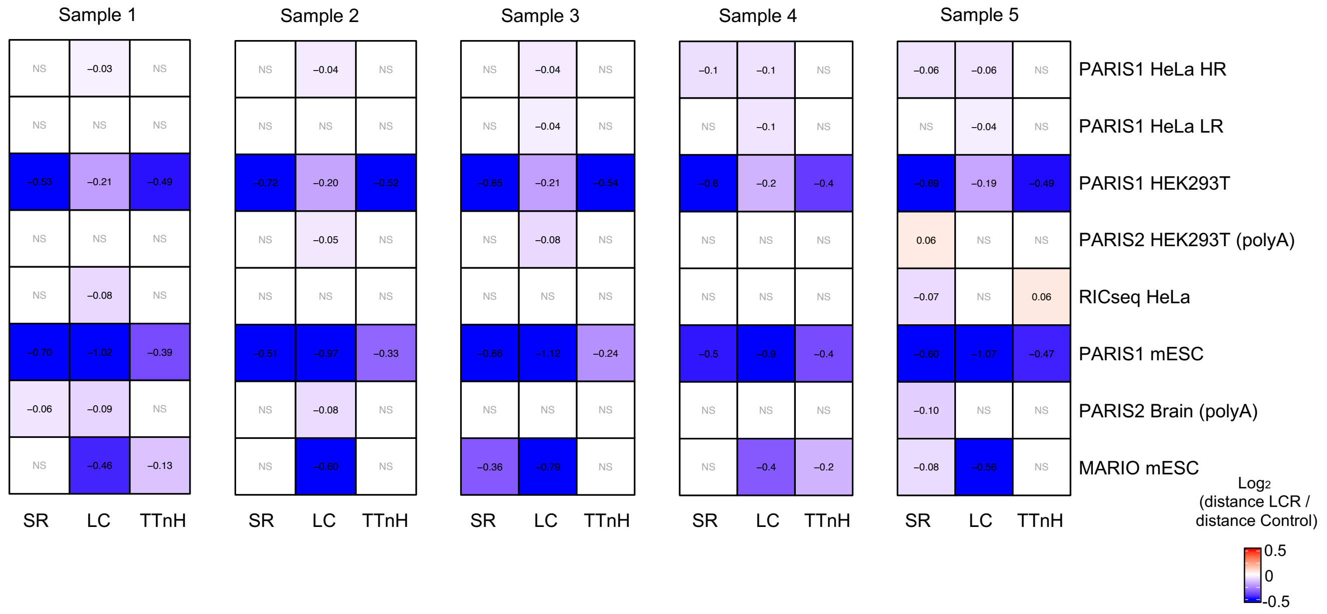

**c**

**Distance of LCR-bearing RRIs from the network center compared with that of control RRIs with matched expression**

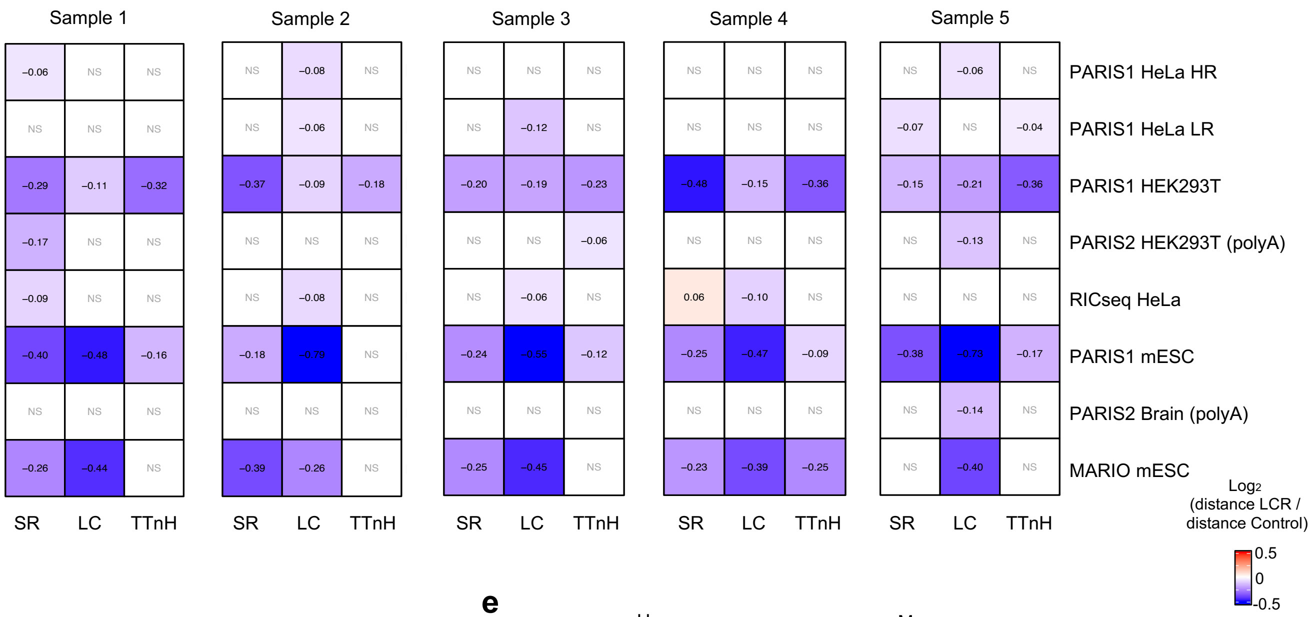


**d**

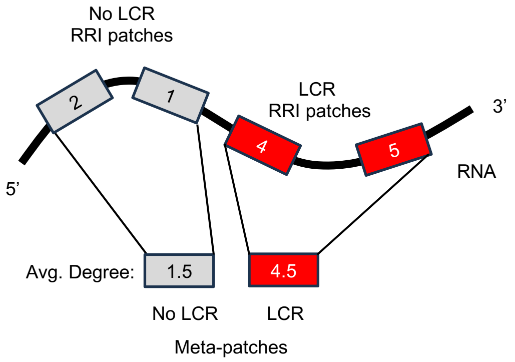


**e**

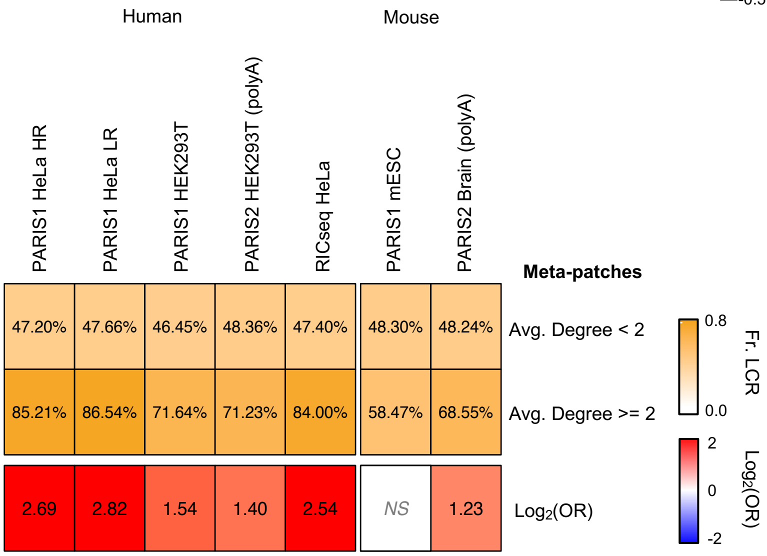

**Supplementary Fig. 3. RNA–RNA contacts involving LCRs exhibit region-specific enrichment patterns and high connectivity. a**, Heatmap depicting, for each transcriptome-wide experiment (rows), the $\log_2$(Odds Ratio), representing the nucleotide enrichment of SR (top), LC (middle) and TTnH (bottom) repeats in RRIs, calculated for each meta-transcript bin. Significant enrichment or depletion of repeats is indicated by red or blue colors, respectively; white denotes the absence of statistical significance, including cases with no nucleotide overlap. Statistical significance was assessed usingtwo-sided Fisher's exact test with Bonferroni correction for multiple testing. The bar plot above the heatmap shows, for each tested repeat family, the number of transcriptome-wide experiments where a statistically significant association between repeats and RRIs was found (red for enrichment, blue for depletion) or not (gray). **b**, Heatmaps depicting the distance from the RRI network center for interactions overlapping with SR, LC, or TTnH repeats, compared to a control RRI group. The analysis was conducted using five independent samplings of RNA–RNA interactions, presented sequentially from left to right, that were built as described in the **Methods** section. Each cell represents the $\log_2$ ratio of the median distance of LCR-containing RRIs to that of control RRIs. Colored cells with numbers indicate significant differences, while non-significant results are labeled as NS. The color gradient ranges from dark blue (shorter distances relative to controls) to light orange (longer distances). Statistical significance was determined using two-tailed Mann-Whitney U-test. **c**, Same as **b**, but with control RRIs chosen to match the expression of the RNAs involved in LCR-bearing interactions. **d**, Schematic representation of an RNA with both types of RRI patches: with or without LCR. Gray boxes mark patches without LCR, red boxes mark LCR-bearing patches. Numbers inside boxes denote the degree (number of RRI contacts overlapping with that patch). For each class (No LCR and LCR), a meta-patch score is computed as the mean degree across patches of that class on the same RNA. **e**, Heatmap summarizing, for each analyzed human RRI dataset, the enrichment of LCRs in meta-patches with average degree $\geq 2$ compared to those with average degree $< 2$. The top panel shows the fraction of LCR meta-patches in the two groups, while the bottom panel displays the $\log_2$(Odds Ratio). Red indicates significant enrichment of LCRs among the high-degree meta-patches, and NS denotes non-significant differences. Statistical significance was assessed using two-sided Fisher's exact test followed by Benjamini–Hochberg correction for multiple tests.

### Supplementary Fig. 4

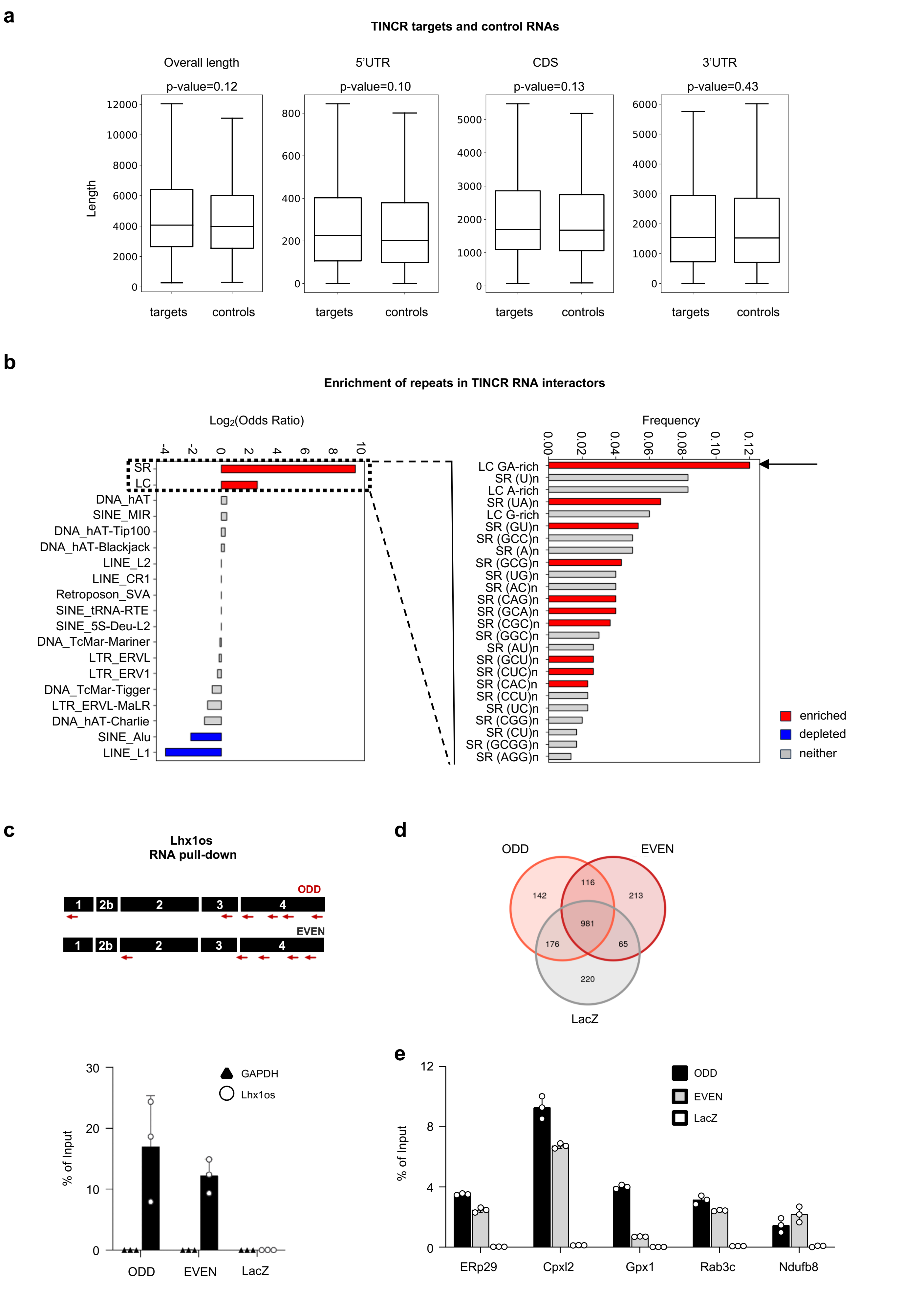

**f**

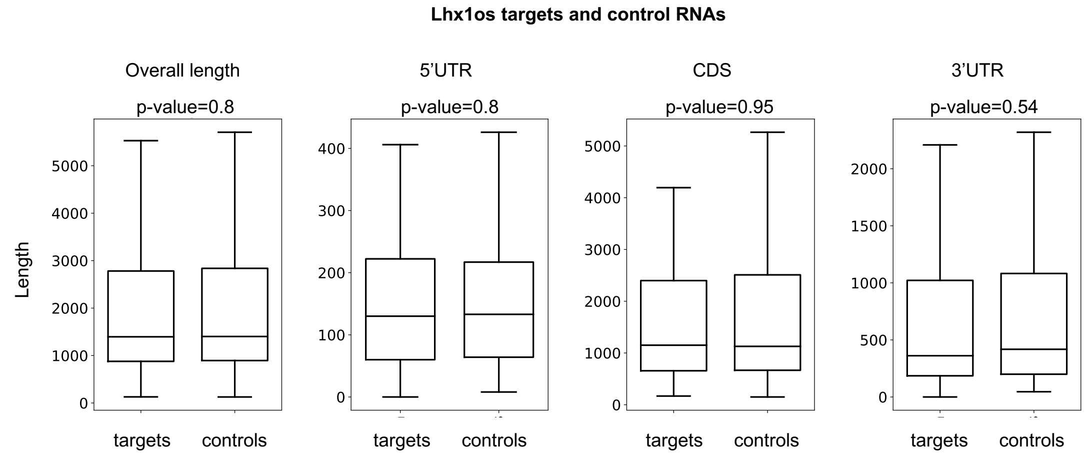
Lhx1os targets and control RNAs
Overall length
p-value=0.8
5'UTR
p-value=0.8
CDS
p-value=0.95
3'UTR
p-value=0.54
Length
0
1000
2000
3000
4000
5000
0
100
200
300
400
0
1000
2000
3000
4000
5000
0
500
1000
1500
2000
targets
controls
targets
controls
targets
controls
targets
controls

**g**

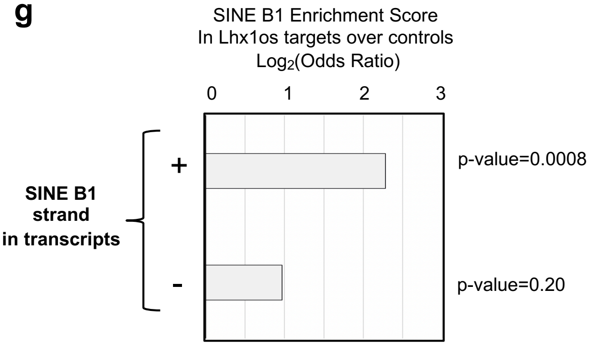
SINE B1 Enrichment Score
In Lhx1os targets over controls
Log2(Odds Ratio)
0
1
2
3
SINE B1
strand
in transcripts
+
-
p-value=0.0008
p-value=0.20

**h**

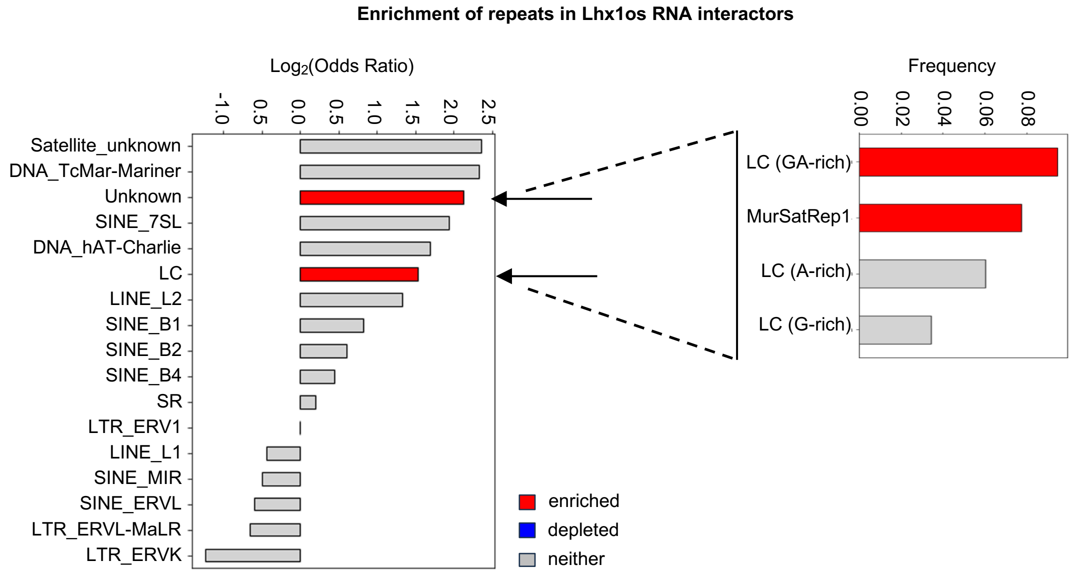
Enrichment of repeats in Lhx1os RNA interactors
Log2(Odds Ratio)
-1.0
0.5
0.0
0.5
1.0
1.5
2.0
2.5
Satellite_unknown
DNA_TcMar-Mariner
Unknown
SINE_7SL
DNA_hAT-Charlie
LC
LINE_L2
SINE_B1
SINE_B2
SINE_B4
SR
LTR_ERV1
LINE_L1
SINE_MIR
SINE_ERVL
LTR_ERVL-MaLR
LTR_ERVK
enriched
depleted
neither
Frequency
0.00
0.02
0.04
0.06
0.08
LC (GA-rich)
MurSatRep1
LC (A-rich)
LC (G-rich)

**i**

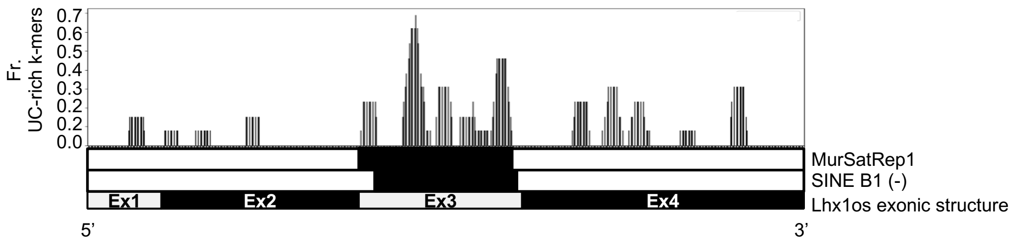
Fr.
UC-rich k-mers
0.7
0.6
0.5
0.4
0.3
0.2
0.2
0.0
MurSatRep1
SINE B1 (-)
Ex1
Ex2
Ex3
Ex4
Lhx1os exonic structure
5'
3'

**Supplementary Fig. 4. Complementary low-complexity repeats drive enrichment in lncRNA-target interactions. a**, Box plots showing the length distribution of entire transcripts and specific regions (5'UTR, CDS, and 3'UTR) for TINCR targets and control RNAs (see **Methods**). Boxes show the interquartile range (IQR), with the line indicating the median; whiskers extend to the most extreme data points within 1.5×IQR from the median; outliers were not included. Statistical significance was evaluated using two-tailed Mann-Whitney U-test. **b**, Bar plot depicting the enrichment of repeated elements in the TINCR targets. For each repeat family, the left panel reports the $log_2$(Odds Ratio) calculated based on the number of TINCR targets and controls with and without at least one repeat. The right panel indicates the frequency of targets hosting specific SRs and LCs. Only the repeated families and elements identified in at least one RNA of the analyzed groups were plotted. Red and blue bars indicate statistically significant enrichment and depletion of repeats, respectively. P-values were calculated using two-sided Fisher's exact test and corrected for multiple comparisons using the Benjamini-Hochberg method. **c**, Top panel: schematic representation of the Lhx1os regions targeted by the ODD and EVEN antisense oligonucleotide probe sets, with black boxes indicating distinct Lhx1os exons. Lower panel: bar plot illustrating qRT-PCR results for GAPDH and Lhx1os enrichment in RNA extracted from endogenous pull-down experiments (n=3) performed in DIV3 MNs (EB cells dissociated at day 6 and replated for additional 3 days) for each probe set used (ODD, EVEN, LacZ). Data are expressed in percentage of Input and presented as the mean ± s.d. Individual data points for each replicate are overlaid. **d**, Venn Diagram showing the intersection of RNAs enriched in Lhx1os pull-down RNA-seq experiment using EVEN, ODD or LacZ probe sets. **e**, Bar plot illustrating a representative result of the qRT-PCR analysis of the enrichment of the indicated RNAs in extract from endogenous pull-down experiments performed in DIV3 MNs (EB cells dissociated at day 6 and replated for additional 3 days) for each probe set used (ODD, EVEN, LacZ). Data are expressed in percentage of input and presented as the mean ± s.d of the technical replicates (n=3). Individual data points for each replicate are overlaid. **f**, Box plots showing the length distribution of entire transcripts and specific regions (5'UTR, CDS, and 3'UTR) for Lhx1os targets and control RNAs (see **Methods**). Boxes show the interquartile range (IQR), with the line indicating the median; whiskers extend to the most extreme data points within 1.5×IQR from the median; outliers were not included. Statistical significance was evaluated using two-tailed Mann-Whitney U-test. **g**, Bar plot representing the $log_2$(Odds Ratio) of sense (+) or antisense (-) SINE B1 enrichment in Lhx1os pull-down targets compared to controls. The analysis distinguishes between SINE B1 strand orientations (sense and antisense). Statistical significance was assessed using two-sided Fisher's exact test. **h**, Bar plot depicting the enrichment of repeated elements in the Lhx1os targets. For each repeat family, the left panel reports the $log_2$(Odds Ratio) calculated based on the number of Lhx1os targets and controls with and without at least one repeat. The right panel indicates the fraction of targets hosting specific repeated elements. Only the repeated families and elements identified in at least one RNA of the analyzed groups were plotted. Red and blue bars indicate statistically significant enrichment and depletion of repeats, respectively. P-values were calculated using two-sided Fisher's exact test and corrected for multiple comparisons using the Benjamini-Hochberg method. **i**, Bar plot displaying the frequency of UC-rich k-mers complementary to GA-rich low-complexity regions along the Lhx1os transcript, mapped from the 5' to the 3' end. Below the plot, the Lhx1os exonic structure is depicted, highlighting the positions of exons (Ex1 to Ex4). Overlaps with SINE B1 elements in the antisense orientation (-) and MurSatRep1 repeats are indicated, showcasing their co-localization with regions of elevated UC-rich k-mer frequencies, particularly within Exon 3.

### Supplementary Fig. 5

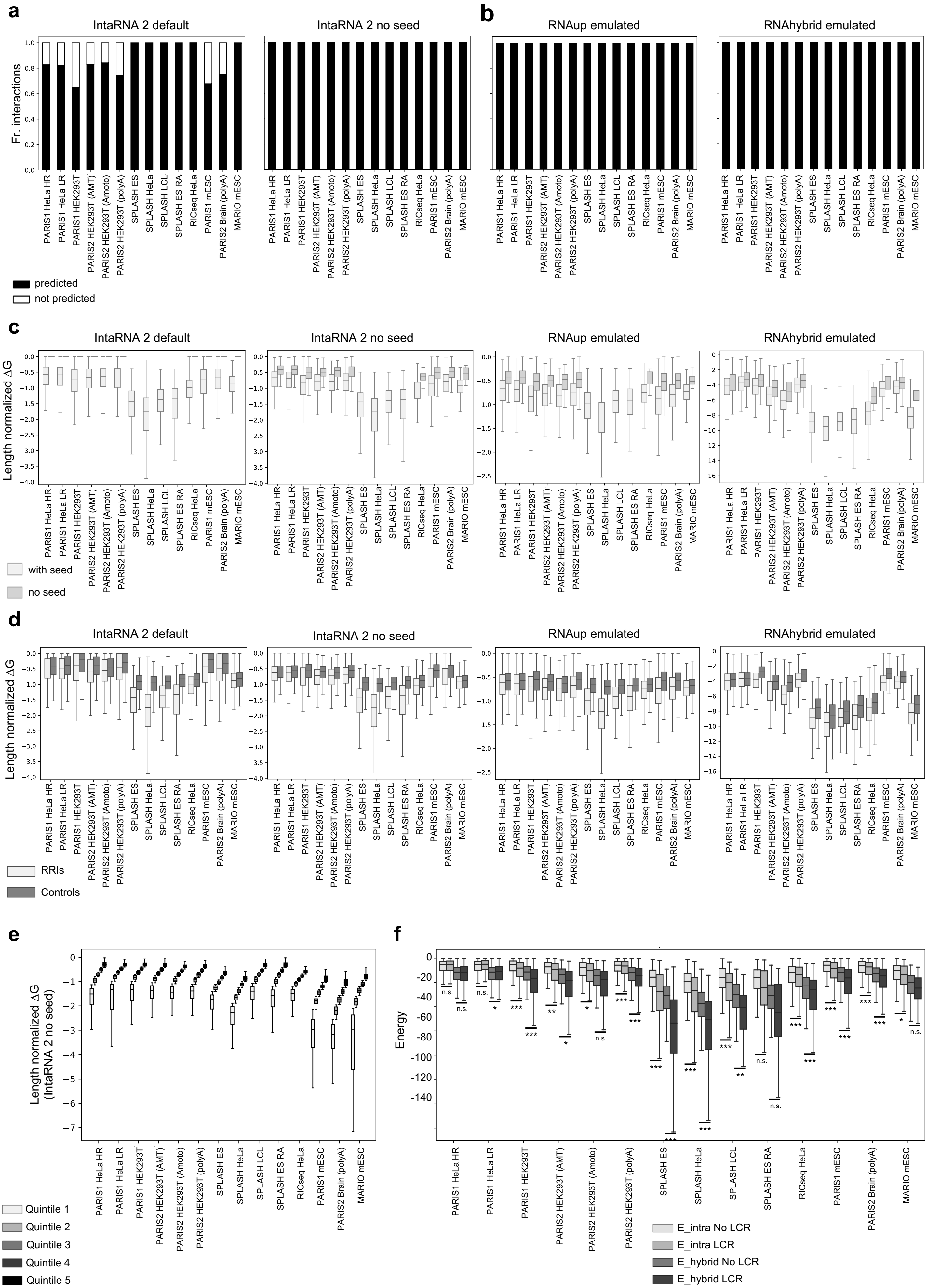

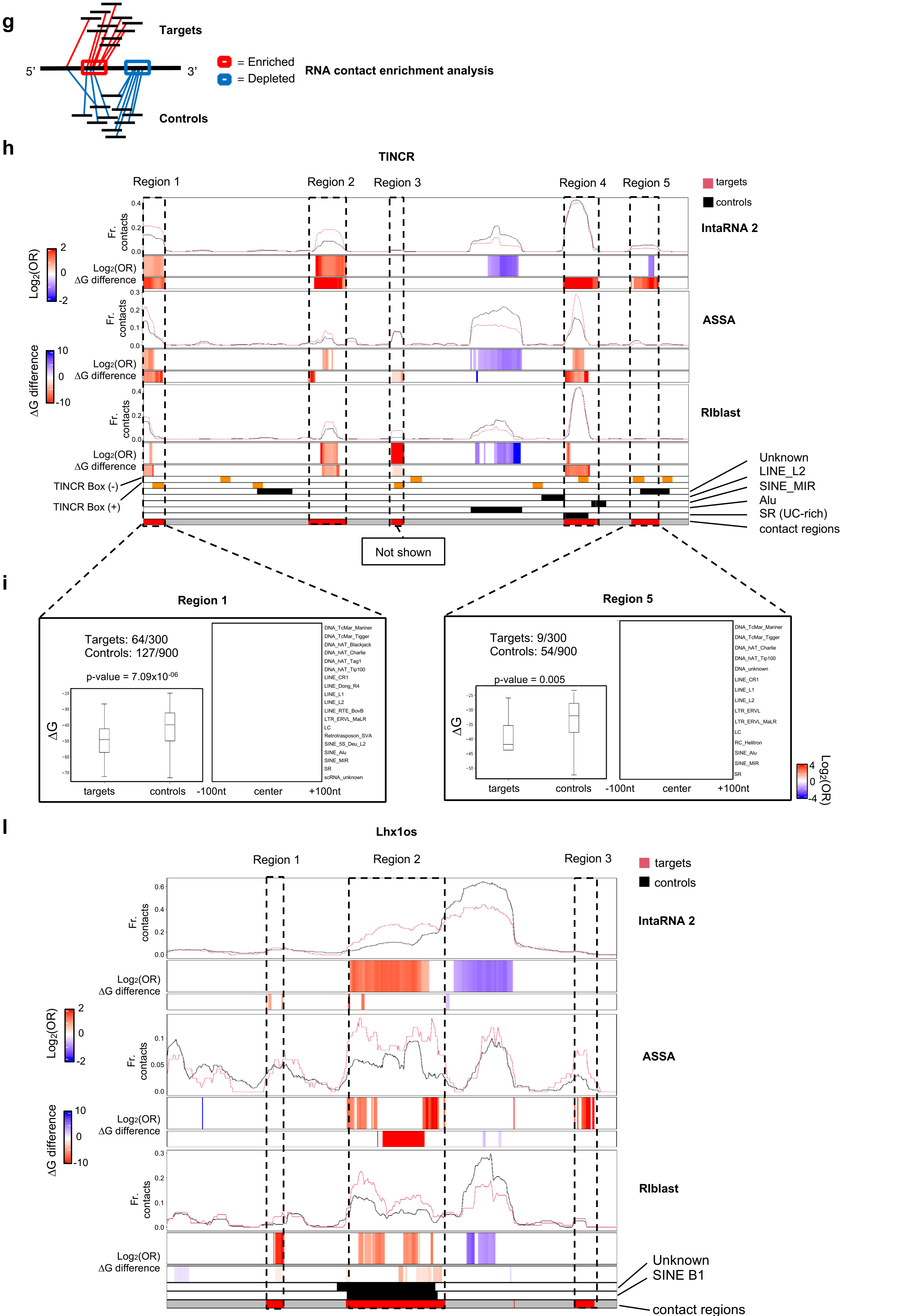
g
Targets
5'
3'
Controls
= Enriched
= Depleted
RNA contact enrichment analysis
h
TINCR
Region 1
Region 2
Region 3
Region 4
Region 5
targets
controls
IntaRNA 2
ASSA
RIblast
Fr. contacts
Log2(OR)
ΔG difference
Log2(OR)
ΔG difference
TINCR Box (-)
TINCR Box (+)
Unknown
LINE_L2
SINE_MIR
Alu
SR (UC-rich)
contact regions
Not shown
i
Region 1
Targets: 64/300
Controls: 127/900
p-value = 7.09x10-06
ΔG
targets
controls
-100nt
center
+100nt
DNA_TcMar_Mariner
DNA_TcMar_Tigger
DNA_hAT_Blackjack
DNA_hAT_Charlie
DNA_hAT_Tag1
DNA_hAT_Tip100
LINE_CR1
LINE_Dong_R4
LINE_L1
LINE_L2
LINE_RTE_BovB
LTR_ERVL_MaLR
LC
Retrotransposon_SVA
SINE_5S_Deu_L2
SINE_Alu
SINE_MIR
SR
scRNA_unknown
Region 5
Targets: 9/300
Controls: 54/900
p-value = 0.005
DNA_TcMar_Mariner
DNA_TcMar_Tigger
DNA_hAT_Charlie
DNA_hAT_Tip100
DNA_unknown
LINE_CR1
LINE_L1
LINE_L2
LTR_ERVL
LTR_ERVL_MaLR
LC
RC_Helitron
SINE_Alu
SINE_MIR
SR
Log2(OR)
l
Lhx1os
Region 1
Region 2
Region 3
targets
controls
IntaRNA 2
ASSA
RIblast
Unknown
SINE B1
contact regions

**Supplementary Fig. 5. Thermodynamically stable RNA–RNA interactions are associated with LCRs. a** and **b**, Stacked bar plots showing the fraction of interactions with successful or failed thermodynamic energy predictions across different transcriptome-wide experiments. Each bar represents the fraction of interactions for which thermodynamic energy predictions either failed (white) or succeeded (black) using IntaRNA 2 with various parameter configurations: default (**a**, left panel) and without seed constraint (**a**, right panel); emulating the behavior of RNAup (**b**, left panel) and RNAhybrid (**b**, right panel) algorithms. Noteworthy, the probability of identifying a seed between interacting regions is influenced by their length. **c**, Box plot displaying the distribution of length-normalized ΔG values (see **Methods**) for RNA–RNA interactions across different transcriptome-wide experiments, calculated using different IntaRNA 2 parameter configurations. Interactions containing a seed (successful IntaRNA 2 default predictions) are shown in light gray, while those without a seed (failed IntaRNA 2 default predictions) are represented in dark gray. A value of 0 was assigned to failed predictions. **d**, Box plot displaying the length-normalized ΔG values (y-axis) for RNA–RNA interactions and random controls across multiple transcriptome-wide experiments (x-axis). ΔG values were calculated using IntaRNA 2 without seed constraint. **e**, Box plot displaying the length-normalized ΔG values (y-axis) for RNA–RNA interactions across multiple transcriptome-wide experiments (x-axis). ΔG values were calculated using IntaRNA 2 without seed constraint. Each box corresponds to a quintile of the ΔG distribution, ranging from quintile 1 (light gray) to quintile 5 (black). **f**, Box plot displaying the energy of hybridization (E_hybrid) and the energy related to intramolecular interactions (E_intra), defined as the negative sum of the accessibility penalties for the query and target molecules (-[ED1+ ED2]), calculated for LCR-bearing (LCR) and non-LCR-bearing (No LCR) RRIs across multiple transcriptome-wide experiments (x-axis). Energy values were calculated using IntaRNA 2 without seed constraint. Statistical significance was assessed using two-tailed Mann-Whitney U tests with Bonferroni correction for multiple comparisons. n.s., not significant; *, $p < 0.05$; **, $p < 0.01$; ***, $p < 0.001$; ****, $p < 0.0001$. Boxes show the interquartile range (IQR), with the line indicating the median; whiskers extend to the most extreme data points within 1.5×IQR from the median; outliers were not included. **g**, Diagram illustrating the RNA contact enrichment analysis, which compares the predicted RNA–RNA interactions of a lncRNA (depicted as a horizontal black line spanning from 5' to 3') with two RNA groups: its RNA targets (red lines) and a control group (blue lines). Regions on the target RNA that preferentially interact with the RNA targets are highlighted with red boxes, while blue boxes indicate regions with reduced interactions compared to the control group. **h**, Heatmap displaying the results of the RNA contact enrichment analysis for the TINCR RNA sequence using three RNA–RNA interaction prediction tools: IntaRNA 2 (top), ASSA (middle) and RIblast (bottom). TINCR sequence is represented from the 5' end (left) to the 3' end (right) in segments of 50 nucleotides, with a 10-nucleotide sliding step. For each software, three consecutive panels resume the results of the analysis: the first panel (line plot) illustrates the frequency of RNAs from the target (red line) or the control (black line) group predicted by the software to be in contact with each segment. The second panel (heatmap) displays $\log_2$(Odds Ratio) values, which measure the enrichment of contacts with target RNAs compared to control RNAs. Significant enrichment or depletion of contacts is indicated by red or blue colors, respectively; white denotes the absence of statistical significance. Statistical significance was assessed using two-sided Fisher's exact test. The third panel (heatmap) reports the difference between the median ΔG values of contacts with the target and control groups. Significant negative or positive difference of ΔG medians is indicated by red or blue colors, respectively; white denotes the absence of statistical significance. Statistical significance was assessed using two-tailed Mann-whitney U-test. Orange boxes in the bottom annotation of the heatmap correspond to segments containing the TINCR box motifs in sense (+) or antisense (-) orientation. Black boxes correspond to bins with repeated elements from different families. Red boxes, also highlighted with dashed rectangles, indicate the 5 TINCR regions that displayed preferential binding with target RNAs according to at least one out of three prediction tools. **i**, Visual representation of the results of RNA contact enrichment analysis performed on TINCR region 1 (left panel) and 5 (right panel). The text in the top-left area reports the amount of RNAs from the target and control groups that are predicted to contact the TINCR region. The box plot in the bottom-left area reports the ΔG distribution of predicted interactions between the TINCR region and the two transcript sets, calculated using IntaRNA 2. Boxes show the interquartile range

(IQR), with the line indicating the median; whiskers extend to the most extreme data points within 1.5×IQR from the median; outliers were not included. Statistical significance was assessed using two-tailed Mann-whitney U-test. The heatmap in the right panel presents the results of the meta-region analysis of repeats. Each row represents a repeat family, and the columns correspond to nucleotide positions within a 200-nucleotide zone centered on regions from target or control RNAs that are predicted to contact the TINCR region. The enrichment or depletion of repeats in target RNA contact regions with respect to control RNAs is displayed in each cell of the heatmap. Significant enrichment or depletion of contacts is indicated by red or blue colors, respectively; white denotes the absence of statistical significance. Statistical significance was assessed using two-sided Fisher's exact test. **l**, Heatmap displaying the results of the RNA contact enrichment analysis for the Lhx1os RNA sequence using three RNA–RNA interaction prediction tools: IntaRNA 2 (top), ASSA (middle) and RIblast (bottom). Lhx1os sequence is represented from the 5' end (left) to the 3' end (right) in segments of 10 nucleotides, with a 1-nucleotide sliding step. For each software, three consecutive panels resume the results of the analysis: the first panel (line plot) illustrates the frequency of RNAs from the target (red line) or the control (black line) group predicted by the software to be in contact with each segment. The second panel (heatmap) displays $\log_2$(Odds Ratio) values, which measure the enrichment of contacts with target RNAs compared to control RNAs. Significant enrichment or depletion of contacts is indicated by red or blue colors, respectively; white denotes the absence of statistical significance. Statistical significance was assessed using two-sided Fisher's exact test. The third panel (heatmap) reports the difference between the median $\Delta G$ values of contacts with the target and control groups. Significant negative or positive difference of $\Delta G$ medians is indicated by red or blue colors, respectively; white denotes the absence of statistical significance. Statistical significance was assessed using two-sided Mann-whitney U-test. Black boxes correspond to bins with repeated elements from different families. Red boxes, also highlighted with dashed rectangles, indicate the 3 Lhx1os regions that displayed preferential binding with target RNAs according to at least one out of three prediction tools.

### Supplementary Fig. 6

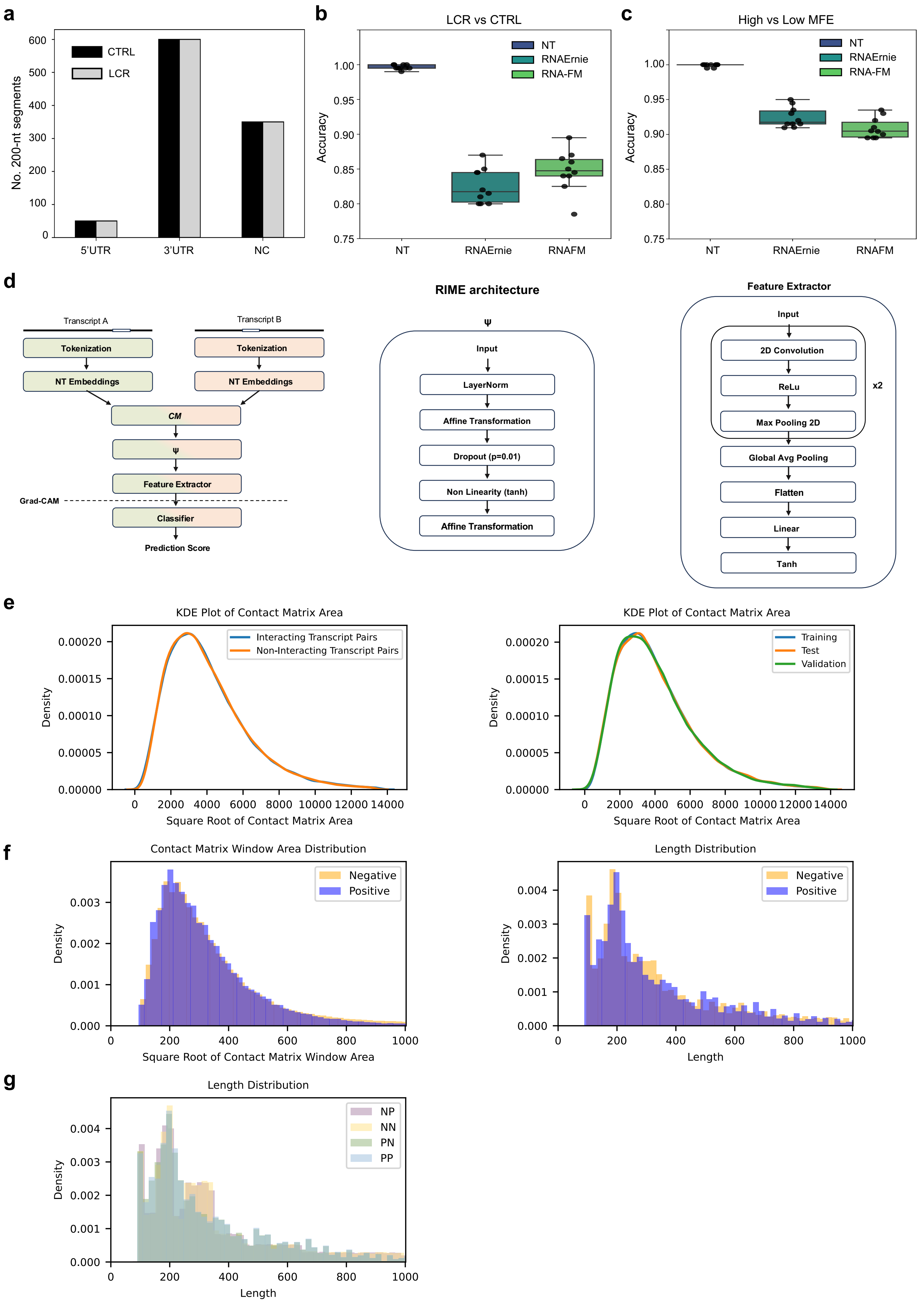

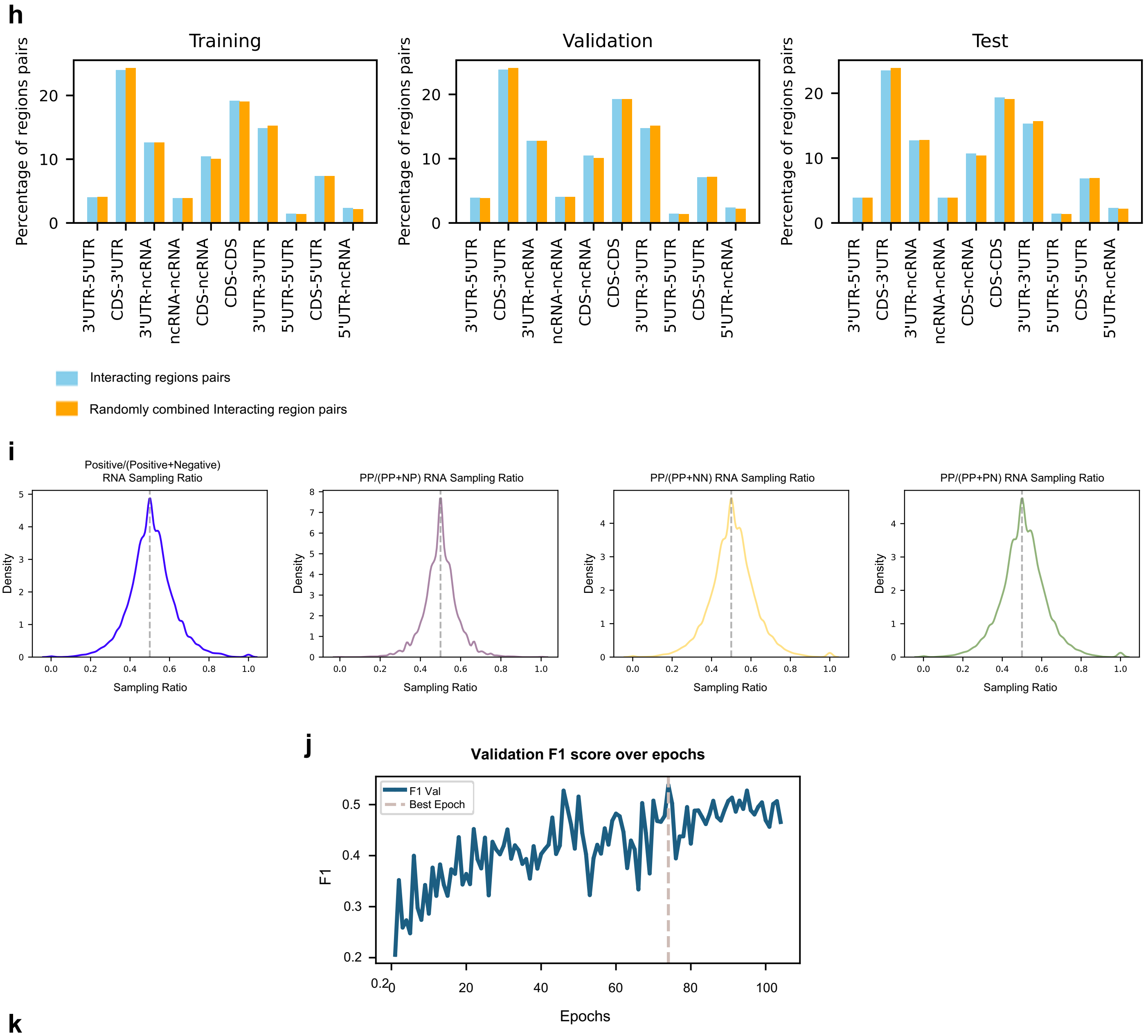


**k**

| Protein-based | RIC-seq (Human) | | | MARIO (Mouse) | | |
|---|---|---|---|---|---|---|
| | RRIs | Interacting Transcript Pairs | Non-Interacting Transcript Pairs | RRIs | Interacting Transcript Pairs | Non-Interacting Transcript Pairs |
| Test | 9,370 | 8,974 | 7,907 | 972 | 970 | 884 |

**Supplementary Fig. 6. Evaluation of nucleic acid language models, RIME architecture and training, validation, and test datasets composition. a,** Bar plot summarizing the distribution of the 1,000 200-nt LCR windows (gray) and CTRL windows (black; see **Methods**) across non-coding transcript regions (5′UTRs, 3′UTRs, and non-coding RNAs). These windows were used to evaluate the ability of nucleic acid language models to capture LCRs and secondary structure stability. **b**, Box plots showing the accuracy from 10-fold cross-validation of logistic regression classifiers trained to distinguish LCR from CTRL windows using three sequence embedding representations: NT, RNAErnie, and RNA-FM. **c**, Box plots showing the accuracy from 10-fold cross-validation of logistic regression classifiers trained to distinguish low MFE from high MFE windows using NT, RNAErnie, and RNA-FM embeddings. Boxes represent the interquartile range (IQR), with the median indicated by the horizontal line; whiskers extend to the most extreme values within 1.5×IQR, and dots denote individual folds. **d**, Schematic illustration of the RIME architecture (left panel). RIME utilizes Nucleotide Transformer (NT) embeddings to predict RNA–RNA interactions. Two RNA sequences, tokenized into 6-mers, are transformed into multidimensional embeddings by the NT model, capturing global sequence context. A Contact Matrix (CM) operation maps the embeddings into a matrix where each point represents a token pair combination. The feature space of the matrix, formed by concatenating token embeddings, undergoes dimensionality reduction through a transformation $\psi$ (central panel) tailored to the task. The contact matrix is processed using a CNN-based Feature Extractor block and a fully connected layer performs binary classification (right panel). Grad-CAM is applied after the last convolutional layer, generating spatial explanations to enhance interpretability of predictions. **e**, Kernel Density Estimate (KDE) plot showing the square root of the contact matrix area, calculated as the product of the lengths of the two full-length RNA sequences, for interacting transcript pairs and non-interacting transcript pairs of the the Psoralen-based dataset (left panel) and across the training, validation, and test sets (right panel). **f**, Distribution histograms showing the square root of the contact matrix windows area (left panel), and the corresponding transcript region lengths (right panel), as sampled from the training set by the data loader over 10 training epochs. Data is shown for both interacting and non-interacting windows. **g**, Histograms showing, for each contact matrix window category (PP, PN, NP, and NN), the distribution of the corresponding transcript region lengths sampled from the training set by the data loader during 10 epochs of training. **h**, Bar plots illustrating the composition of transcript region pairs categories across the training, validation, and test sets. The x-axis represents the 16 possible combinations between transcript regions (5'UTR, CDS, 3'UTR and ncRNA), while the y-axis indicates the percentage of each category. For each category, two bars are shown: one for true interacting transcript pairs (blue) and one for randomly combined pairs (orange). **i**, Density plots illustrating the representation of each RNA in the positive and negative sets during 10 training epochs. Each plot shows the ratio between the number of times each RNA was sampled in the positive set (PP) and the total number of times it was sampled, either against all negative classes combined (PN 50%, NN 25%, NP 25%; first panel) or separately against NP (second panel), NN (third panel), and PN (fourth panel). A value of 0.5 corresponds to equal sampling between positive and negative sets. **j**, Line plot showing the validation F1 score across training epochs. The vertical dashed line marks epoch 74, where RIME achieved its best performance. Training stopped after 30 additional epochs without improvement due to early stopping. **k**, Table reporting the amount of interacting region pairs (RRIs) and transcript pairs included in the independent protein-based RIC-seq and MARIO test sets.

### Supplementary Fig. 7

**a**

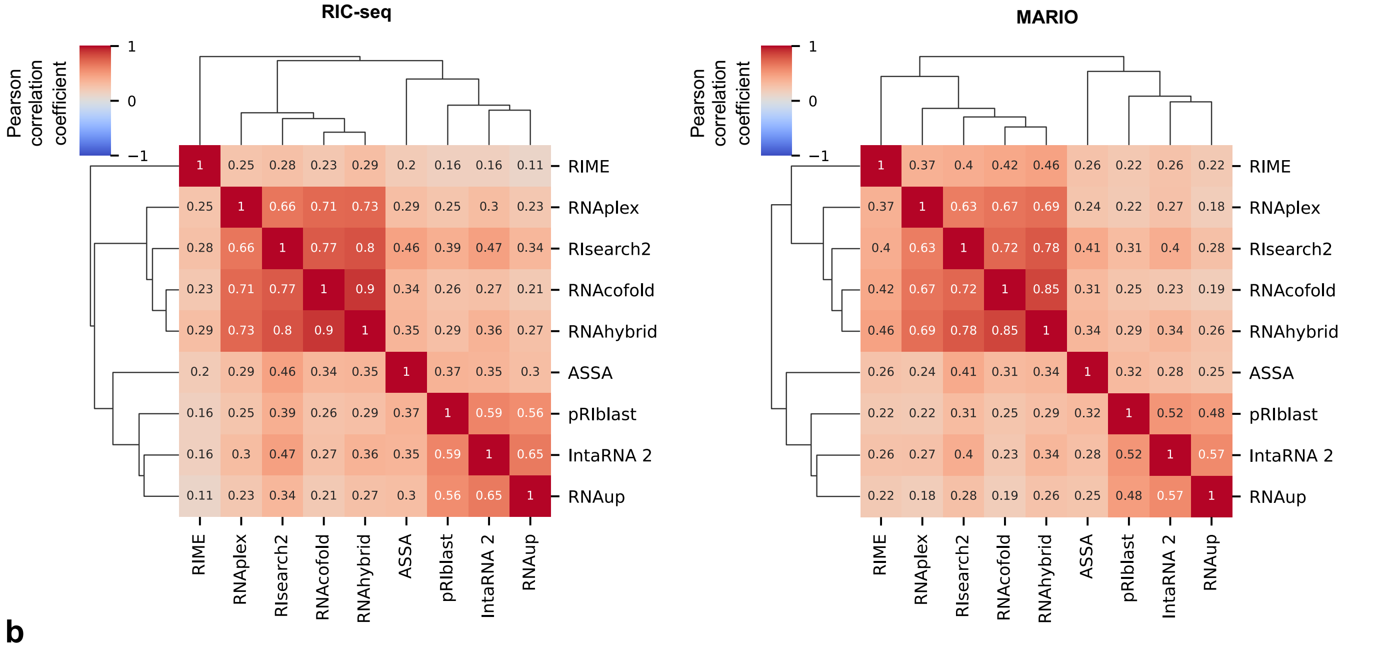


**b**

| Test set 200x200 | | | | |
|---|---|---|---|---|
| | **PP** | **NP** | **PN** | **NN** |
| Psoralen Human | 10,492 | 10,923 | 10,985 | 11,173 |
| Psoralen Mouse | 5,198 | 5,275 | 5,115 | 5,127 |
| RIC-seq | 8,978 | 9,844 | 8,002 | 8,190 |
| MARIO | 970 | 970 | 895 | 895 |

**c**

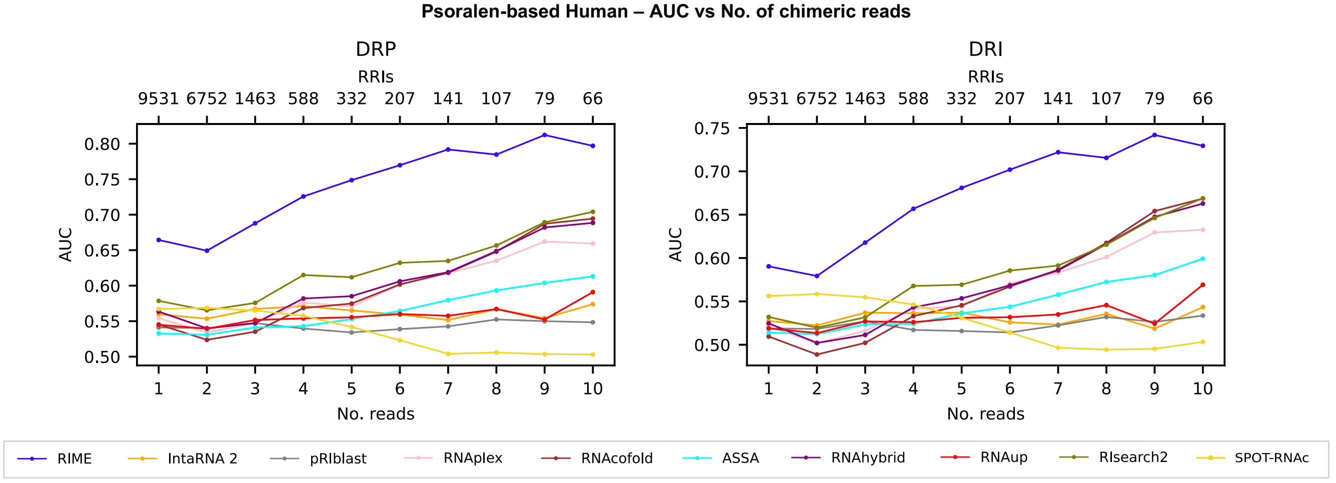


**d**

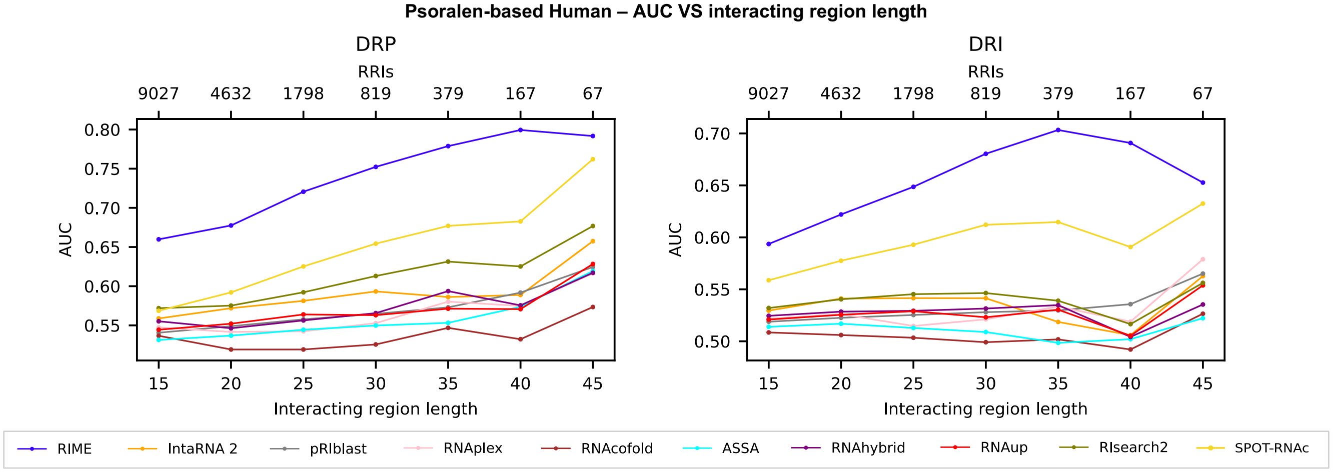

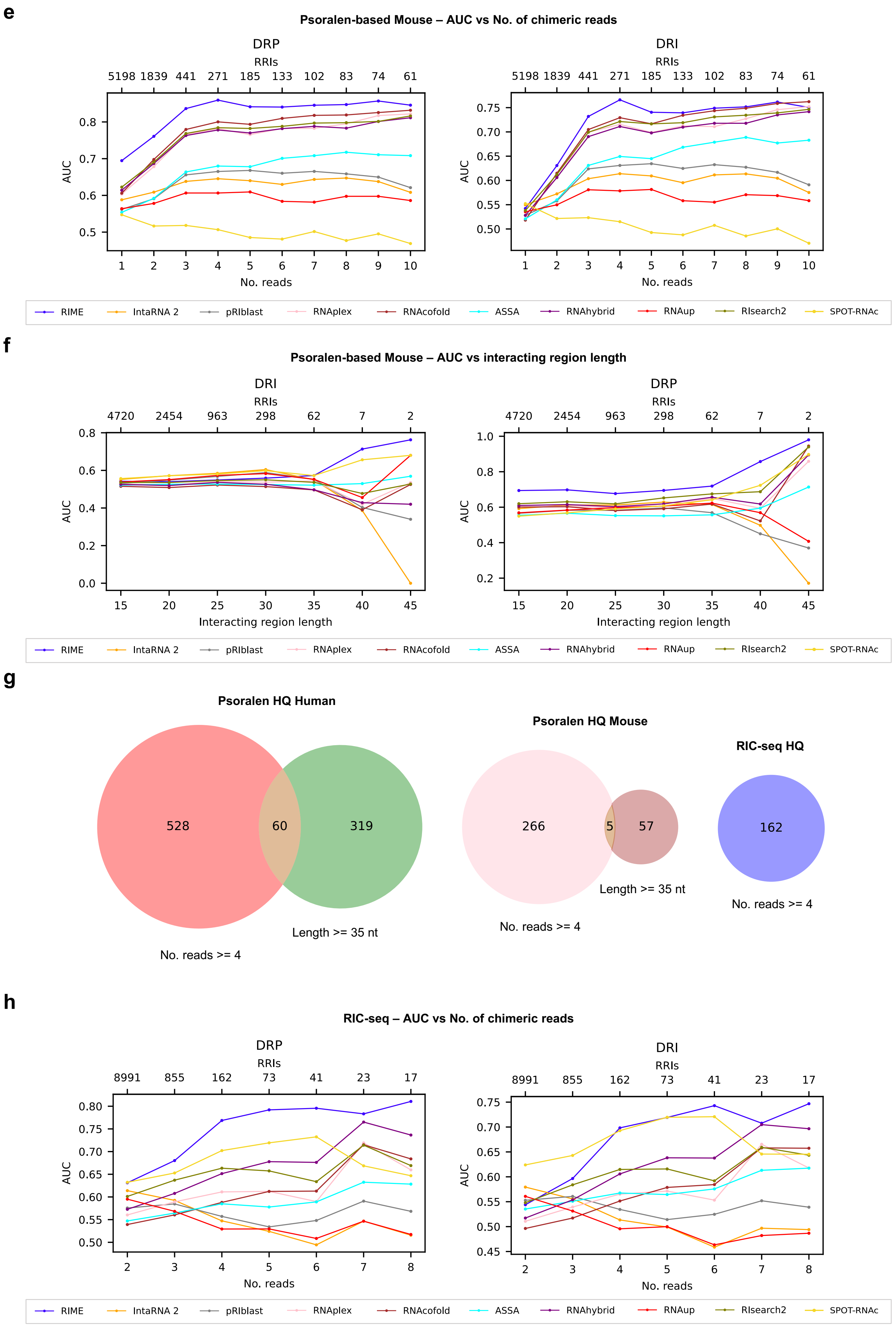

e
Psoralen-based Mouse – AUC vs No. of chimeric reads
DRP
RRIs
5198 1839 441 271 185 133 102 83 74 61
AUC
No. reads
DRI
RRIs
5198 1839 441 271 185 133 102 83 74 61
AUC
No. reads
RIME
IntaRNA 2
pRIblast
RNAplex
RNAcofold
ASSA
RNAhybrid
RNAup
RIsearch2
SPOT-RNAc
f
Psoralen-based Mouse – AUC vs interacting region length
DRI
RRIs
4720 2454 963 298 62 7 2
AUC
Interacting region length
DRP
RRIs
4720 2454 963 298 62 7 2
AUC
Interacting region length
RIME
IntaRNA 2
pRIblast
RNAplex
RNAcofold
ASSA
RNAhybrid
RNAup
RIsearch2
SPOT-RNAc
g
Psoralen HQ Human
528
60
319
No. reads >= 4
Length >= 35 nt
Psoralen HQ Mouse
266
5
57
No. reads >= 4
Length >= 35 nt
RIC-seq HQ
162
No. reads >= 4
h
RIC-seq – AUC vs No. of chimeric reads
DRP
RRIs
8991 855 162 73 41 23 17
AUC
No. reads
DRI
RRIs
8991 855 162 73 41 23 17
AUC
No. reads
RIME
IntaRNA 2
pRIblast
RNAplex
RNAcofold
ASSA
RNAhybrid
RNAup
RIsearch2
SPOT-RNAc

**Supplementary Fig. 7. Comparative evaluation of RIME, SPOT-RNAc and thermodynamics-based tools for RNA–RNA interaction prediction. a**, Heatmaps of Pearson correlation coefficients between RRI prediction tools, calculated using scores assigned to positive interactions in the RIC-seq (left panel) and MARIO (right panel) test sets. For comparability with RIME scores, ΔG values from thermodynamics-based tools were converted by inverting their sign. Tools with similar scoring patterns were grouped using average linkage hierarchical clustering based on Euclidean distances. **b**, Table summarizing the number of PP, NP, PN, and NN areas included in the Psoralen-based Human, Psoralen-based Mouse, RIC-seq and MARIO test sets. For each interacting transcript pair, we generated one PP and one NP. For each non-interacting pair, we generated one PN and one NN. The number of PPs is lower than the total number of RRIs due to filtering steps detailed in the **Methods** section. **c** and **d**, Line plots illustrating the performance (ROC-AUC) of RIME and other RRI prediction tools on Psoralen-based human test sets with progressively higher quality, measured for the DRP (left panel) and DRI (right panel) tasks. The bottom x-axis represents the minimum number of supporting reads (**c**) or the minimum interacting region length (**d**) thresholds. At each threshold value, the positive set was subsampled to match the quality criteria and, to address class imbalance, negative samples were randomly undersampled 100 times to match the number of positives. ROC-AUC scores were averaged across iterations. The number of interactions retained at different threshold values is reported in the top x-axis of each line plot. **e** and **f**, Line plots illustrating the performance (ROC-AUC) of RIME and other RRI prediction tools on Psoralen-based mouse test sets with progressively higher quality, measured for the DRP (left panel) and DRI (right panel) tasks. The bottom x-axis represents the minimum number of supporting reads (**e**) or the minimum interacting region length (**f**) thresholds. At each threshold value, the positive set was subsampled to match the quality criteria and, to address class imbalance, negative samples were randomly undersampled 100 times to match the number of positives. ROC-AUC scores were averaged across iterations. The number of interactions retained at different threshold values is reported in the top x-axis of each line plot. **g**, Venn diagrams illustrating the number of interactions in the Psoralen HQ Human and Psoralen HQ Mouse datasets (selected based on the minimum number of supporting reads or the minimum interacting region length; left and middle panel, respectively), and in the RIC-seq dataset (right panel). **h**, Line plots illustrating the performance (ROC-AUC) of RIME and other RRI prediction tools on RIC-seq test sets with progressively higher quality, measured for the DRP (left panel) and DRI (right panel) tasks. The bottom x-axis represents the minimum interacting region length thresholds. At each threshold value, the positive set was subsampled to match the quality criteria and, to address class imbalance, negative samples were randomly undersampled 100 times to match the number of positives. ROC-AUC scores were averaged across iterations. The number of interactions retained at different threshold values is reported in the top x-axis of each line plot.

### Supplementary Fig. 8

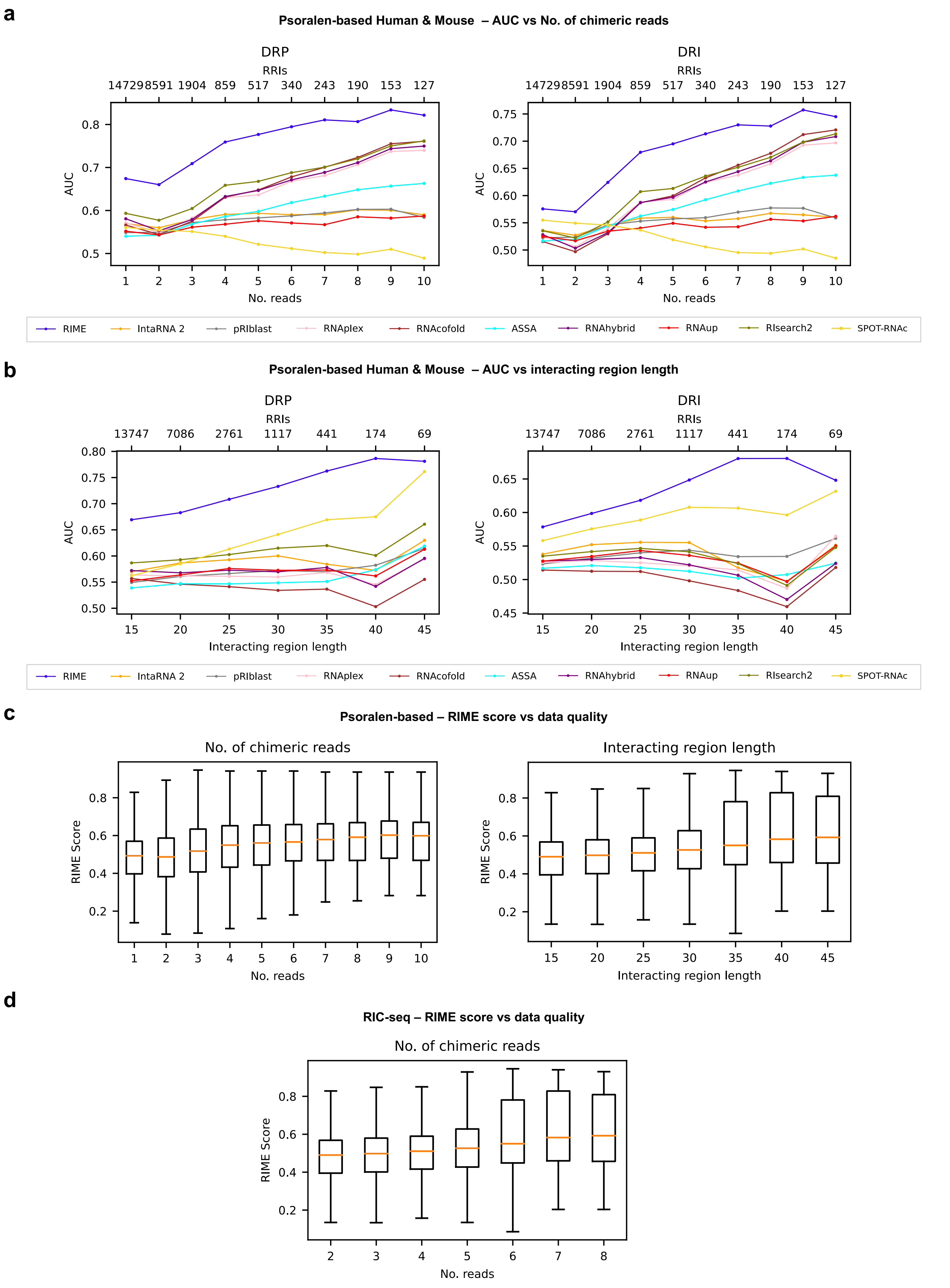

**e**

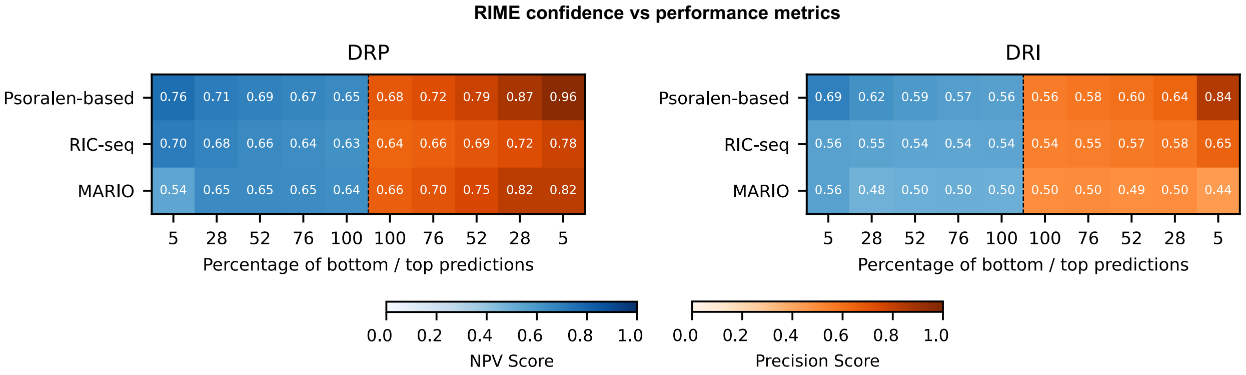
RIME confidence vs performance metrics
DRP
Psoralen-based 0.76 0.71 0.69 0.67 0.65 0.68 0.72 0.79 0.87 0.96
RIC-seq 0.70 0.68 0.66 0.64 0.63 0.64 0.66 0.69 0.72 0.78
MARIO 0.54 0.65 0.65 0.65 0.64 0.66 0.70 0.75 0.82 0.82
5 28 52 76 100 100 76 52 28 5
Percentage of bottom / top predictions
DRI
Psoralen-based 0.69 0.62 0.59 0.57 0.56 0.56 0.58 0.60 0.64 0.84
RIC-seq 0.56 0.55 0.54 0.54 0.54 0.54 0.55 0.57 0.58 0.65
MARIO 0.56 0.48 0.50 0.50 0.50 0.50 0.50 0.49 0.50 0.44
5 28 52 76 100 100 76 52 28 5
Percentage of bottom / top predictions
0.0 0.2 0.4 0.6 0.8 1.0
NPV Score
0.0 0.2 0.4 0.6 0.8 1.0
Precision Score

**f**

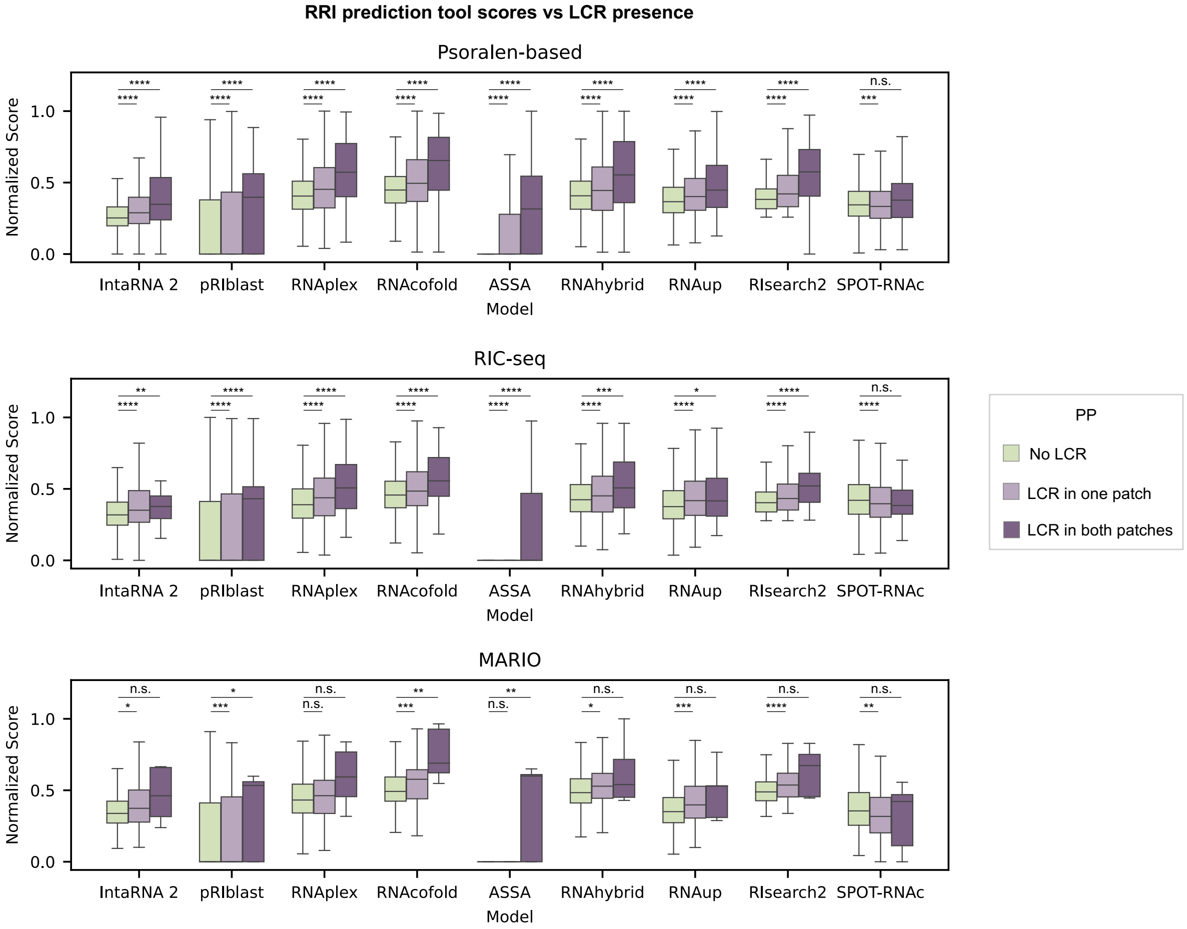
RRI prediction tool scores vs LCR presence
Psoralen-based
RIC-seq
MARIO
Normalized Score
0.0 0.5 1.0
IntaRNA 2 pRIblast RNAplex RNAcofold ASSA RNAhybrid RNAup RIsearch2 SPOT-RNAc
Model
n.s.
PP
No LCR
LCR in one patch
LCR in both patches

**g**

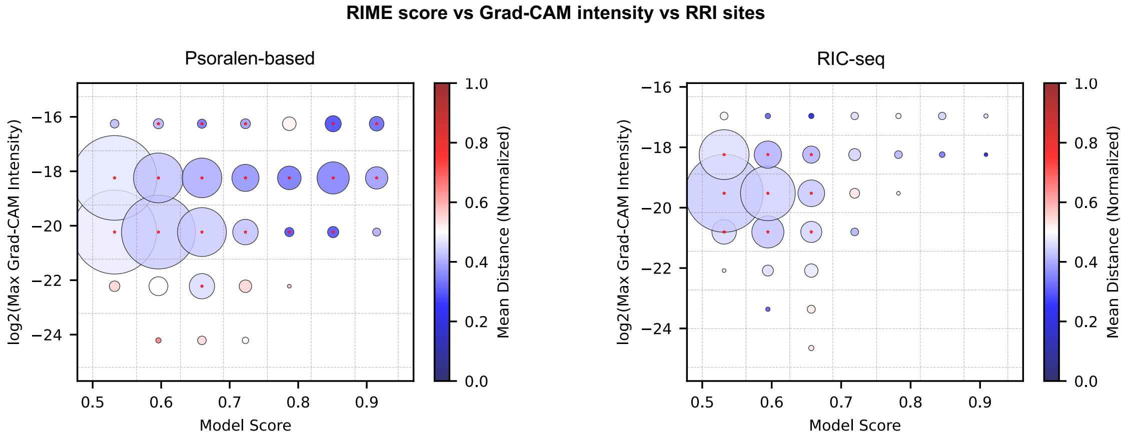
RIME score vs Grad-CAM intensity vs RRI sites
Psoralen-based
RIC-seq
log2(Max Grad-CAM Intensity)
Model Score
0.5 0.6 0.7 0.8 0.9
Mean Distance (Normalized)
0.0 0.2 0.4 0.6 0.8 1.0

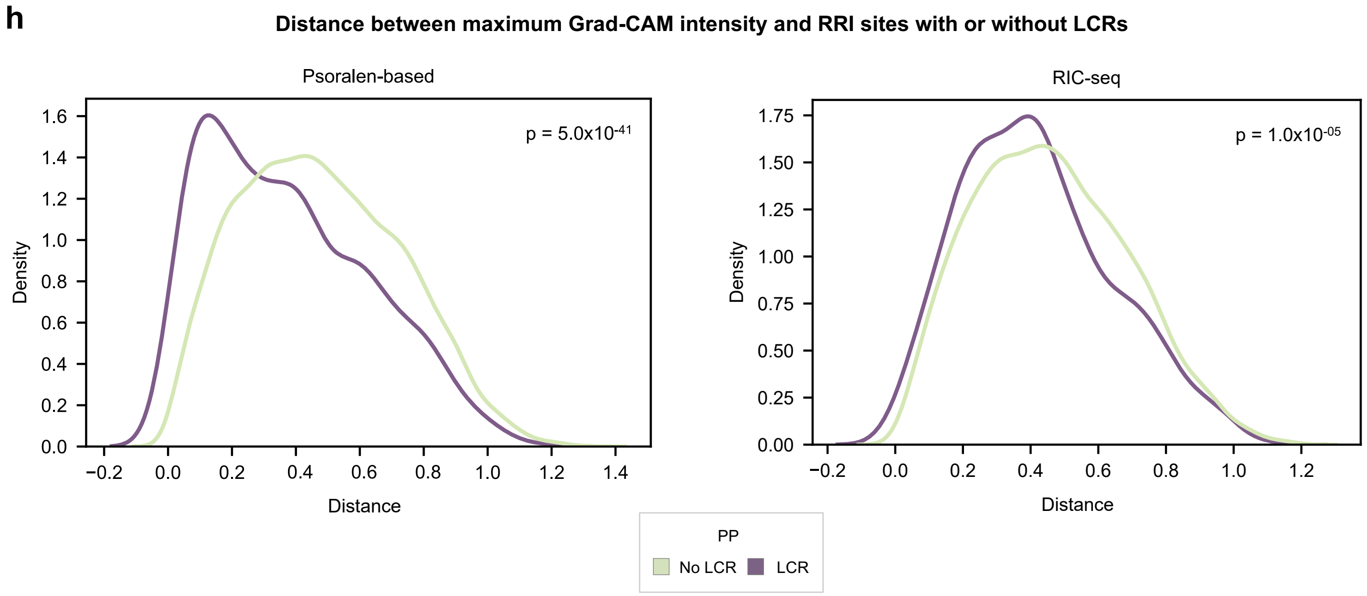
h
Distance between maximum Grad-CAM intensity and RRI sites with or without LCRs
Psoralen-based
p = 5.0x10^-41
Density
Distance
RIC-seq
p = 1.0x10^-05
Density
Distance
PP
No LCR
LCR

**Supplementary Fig. 8. Relationship between data quality, RIME's confidence, and RIME's performance. a** and **b**, Line plots illustrating the performance (ROC-AUC) of RIME and other RRI prediction tools on Psoralen-based test sets (both human and mouse) with progressively higher quality, measured for the DRP (left panel) and DRI (right panel) tasks. The bottom x-axis represents the minimum number of supporting reads (**a**) or the minimum interacting region length (**b**) thresholds. At each threshold value, the positive set was subsampled to match the quality criteria and, to address class imbalance, negative samples were randomly undersampled 100 times to match the number of positives. ROC-AUC scores were averaged across iterations. The number of interactions retained at different threshold values is reported in the top x-axis of each line plot. **c**, Box plots showing the distribution of RIME prediction scores calculated for Psoralen-based test sets with progressively higher quality. The x-axis represents the minimum number of supporting reads (left panel) or the minimum interacting region length (right panel) thresholds used to subset the test set. **d**, Box plot showing the distribution of RIME prediction scores calculated for RIC-seq test sets with progressively higher quality. The x-axis represents the minimum interacting region length (right panel) thresholds used to subset the test set. Boxes show the interquartile range (IQR), with the line indicating the median; whiskers extend to the most extreme data points within 1.5×IQR from the median; outliers were not included. **e**, Heatmaps illustrating the relationship between RIME model confidence and performance metrics, evaluated for the Psoralen-based, RIC-seq and MARIO test sets (rows). The data was divided into segments with progressively higher (top predictions) and progressively lower (bottom predictions) prediction scores. The horizontal axis represents the proportion of data (in percentage) corresponding to top or bottom predictions. Precision (orange) was evaluated for the top predictions, while Negative Predictive Value (NPV, blue) was assessed for the bottom predictions. To address the imbalance between the positive and the negative sets, the data was iteratively undersampled before ranking and selecting the top and bottom predictions. For each proportion of data considered, Precision and NPV scores were averaged over 50 iterative runs. These plots demonstrate how Precision and NPV evolve across different proportions of top and bottom predictions for the DRP (left panel) and DRI (right panel) tasks. **f**, Box plots of thermodynamics-based model scores across Psoralen-based (upper panel), RIC-seq (middle panel), and MARIO (lower panel) test sets. The scores are categorized based on the overlap of LCRs with the interacting patches: pairs where neither patch overlaps with LCRs, pairs where LCRs overlap with one patch, and pairs where both patches overlap with LCRs. ΔG scores were sign-flipped and normalized to the [0, 1] range for comparability, accounting for differences in scale across different tools. Boxes show the interquartile range (IQR), with the line indicating the median; whiskers extend to the most extreme data points within 1.5×IQR from the median; outliers were not included. Two-tailed Mann-Whitney U test was used to test for differences between distributions. P-values were corrected with the Benjamini-Hochberg procedure. *, $p < 0.05$; **, $p < 0.01$; ***, $p < 0.001$; ****, $p < 0.0001$. **g**, Bubble plot displaying the relationship between RIME scores and maximum Grad-CAM intensity values. The plot is divided into grid sections, with each section containing a set of points. The bubble size is proportional to the number of points within each grid section, while the bubble color represents the mean distance between the real centroid of the interaction's bounding box and the maximum Grad-CAM intensity value. A star is placed in the center of the bubble if the distance between the predicted and real centroids is significantly smaller (FDR-adjusted two-tailed Wilcoxon signed-rank test p-value < 0.05) than the theoretical distance of 0.5214, representing the expected distance between the real centroid and a random prediction. **h**, Density plots showing the distribution of distances between maximum Grad-CAM intensity values and the centroid of true interaction sites for RRIs overlapping with LCRs in the region of interaction (LCR) and RRIs without LCRs (No LCR), in Psoralen-based (left) and RIC-seq (right) test sets. P-values from two-tailed Mann-Whitney U-tests (LCR vs No LCR) are reported.

### Supplementary Fig. 9

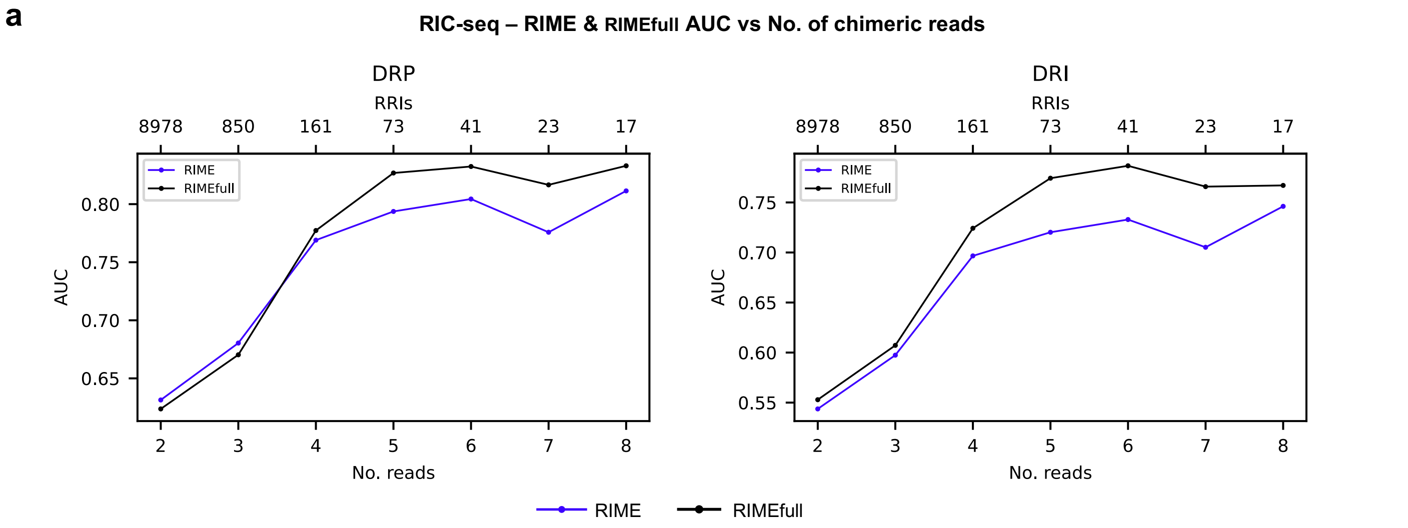


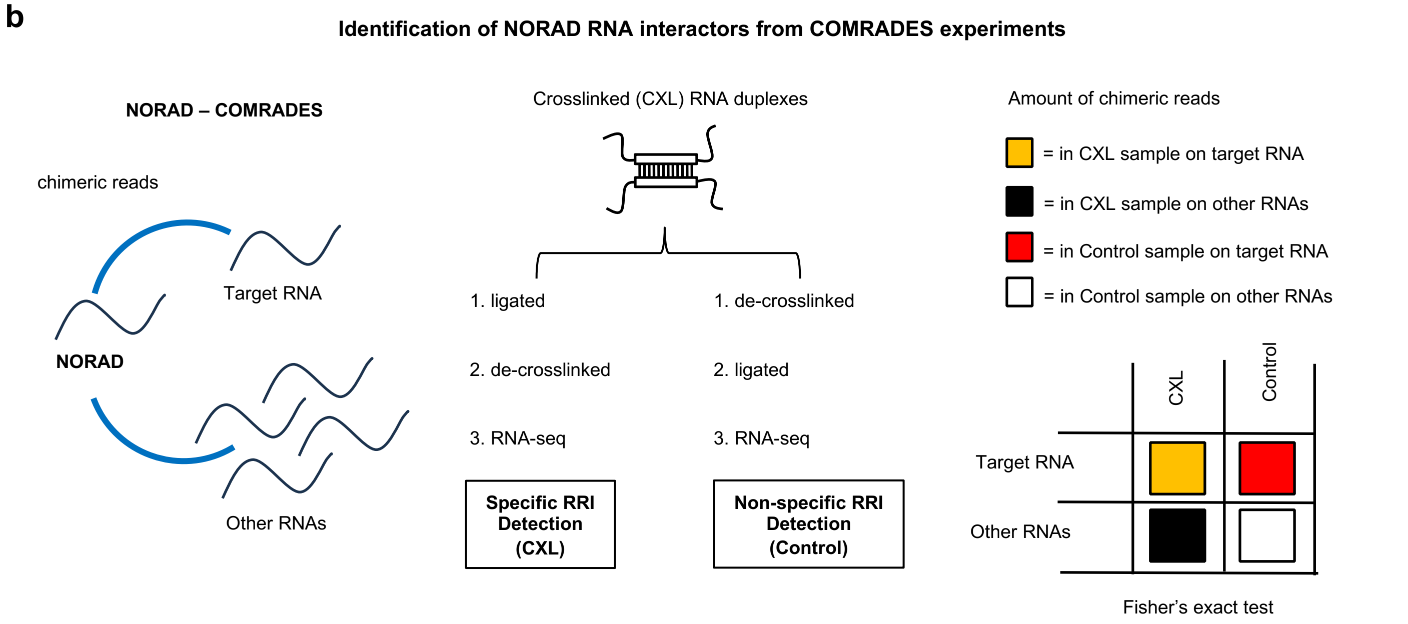


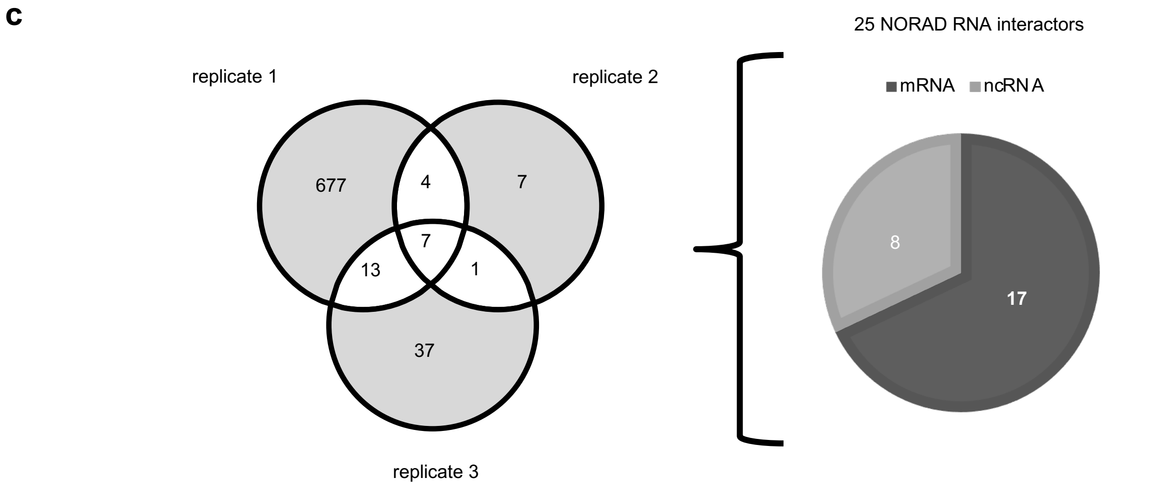

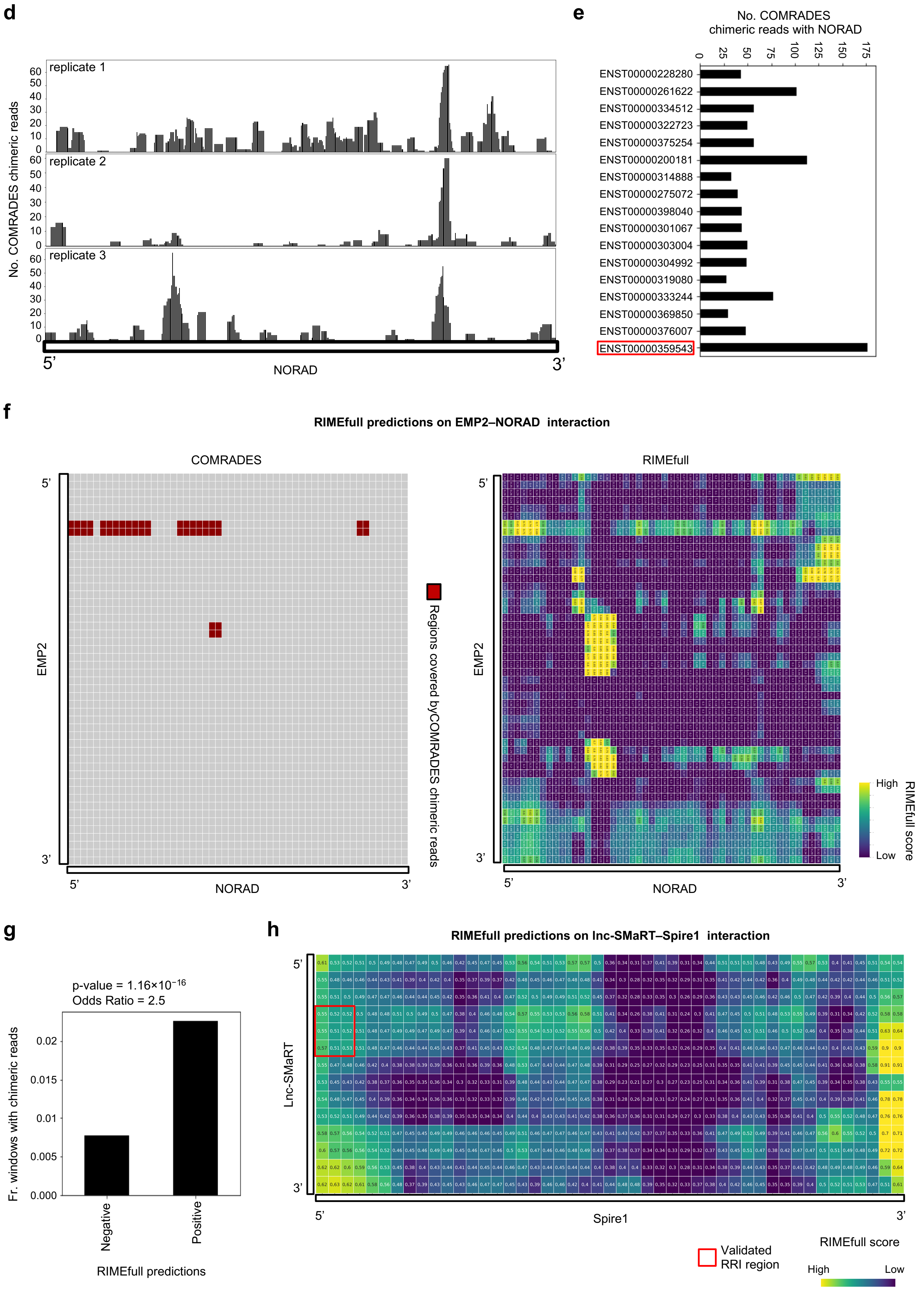
d
No. COMRADES chimeric reads
replicate 1
replicate 2
replicate 3
60
50
40
30
20
10
0
5'
3'
NORAD
e
No. COMRADES chimeric reads with NORAD
0
25
50
75
100
125
150
175
ENST00000228280
ENST00000261622
ENST00000334512
ENST00000322723
ENST00000375254
ENST00000200181
ENST00000314888
ENST00000275072
ENST00000398040
ENST00000301067
ENST00000303004
ENST00000304992
ENST00000319080
ENST00000333244
ENST00000369850
ENST00000376007
ENST00000359543
f
RIMEfull predictions on EMP2–NORAD interaction
COMRADES
EMP2
NORAD
Regions covered byCOMRADES chimeric reads
RIMEfull
High
Low
RIMEfull score
g
p-value = 1.16×10−16
Odds Ratio = 2.5
Fr. windows with chimeric reads
0.02
0.015
0.01
0.005
0.000
Negative
Positive
RIMEfull predictions
h
RIMEfull predictions on lnc-SMaRT–Spire1 interaction
Lnc-SMaRT
Spire1
Validated
RRI region
RIMEfull score
High
Low

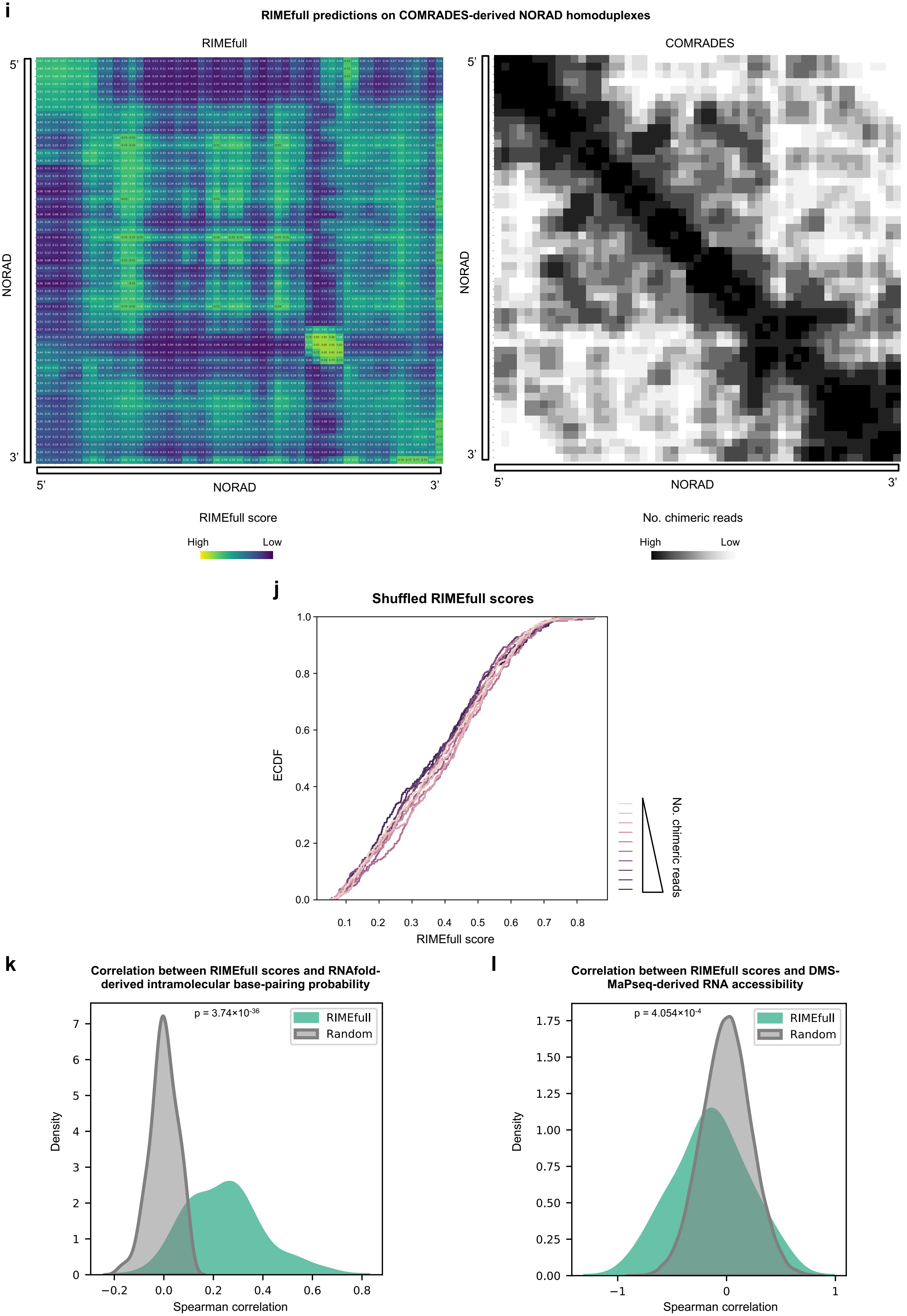
i
RIMEfull predictions on COMRADES-derived NORAD homoduplexes
RIMEfull
COMRADES
5'
3'
NORAD
RIMEfull score
High
Low
No. chimeric reads
High
Low
j
Shuffled RIMEfull scores
ECDF
RIMEfull score
No. chimeric reads
k
Correlation between RIMEfull scores and RNAfold-derived intramolecular base-pairing probability
p = 3.74×10-36
RIMEfull
Random
Density
Spearman correlation
l
Correlation between RIMEfull scores and DMS-MaPseq-derived RNA accessibility
p = 4.054×10-4
RIMEfull
Random
Density
Spearman correlation

**Supplementary Fig. 9. Evaluation of the RIMEfull model. a**, Line plots illustrating the performance (ROC-AUC) of RISE and RISEfull models on RIC-seq test sets with progressively higher quality, measured for the DRP (left panel) and DRI (right panel) tasks. The bottom x-axis represents the minimum interacting region length thresholds. At each threshold value, the positive set was subsampled to match the quality criteria and, to address class imbalance, negative samples were randomly undersampled 100 times to match the number of positives. ROC-AUC scores were averaged across iterations. The number of interactions retained at different threshold values is reported in the top x-axis of each line plot. **b**, Diagram illustrating the approach we adopted for detecting specific and non-specific RNAs interacting with NORAD based on the COMRADES experiment performed by Farberov and coworkers. Chimeric reads represent evidence of interactions between the targeted RNA (e.g., NORAD) and other RNAs. Crosslinked (CXL) RNA duplexes are processed under two conditions: for specific RRI detection (CXL), RNA duplexes are ligated, de-crosslinked, and sequenced, whereas for non-specific RRI detection (Control), duplexes are de-crosslinked first, then ligated and sequenced. Across the three replicates of the experiment, chimeric reads involving NORAD were classified into four groups based on their sample of origin (CXL or Control) and the target RNA to which they were mapped. For each target RNA, the enrichment of chimeric reads connecting to NORAD in the CXL sample compared to the Control sample was calculated. A two-sided Fisher's exact test was applied to determine the statistical significance of the observed enrichment, enabling the identification of specific RNA interactors associated with the NORAD lncRNA. **c**, The left panel shows a Venn diagram depicting the overlap of NORAD interactors identified across the three replicates of the COMRADES experiment. Interactors shared by at least two replicates were selected as *bona fide* specific targets, resulting in a total of 25 RNAs. These interactors were classified based on their biotypes, distinguishing between ncRNA and mRNA, as illustrated in the pie chart in the right panel. **d**, Bar plots reporting, for each NORAD sequence segment, the amount of COMRADES-derived chimeric reads that connect NORAD with its 17 target mRNAs in each experimental replicate. The sequence of the lncRNA is represented from the 5' end (left) to the 3' end (right) in segments of 50 nucleotides, with a 10-nucleotide sliding step. **e,** Bar plot depicting the amount of chimeric reads that connect each target RNA to NORAD lncRNA in all the COMRADES replicates. **f**, The left panel shows a heatmap indicating, for the NORAD–EMP2 transcript pair, the interacting RNA windows supported by COMRADES chimeric reads (red cells). The analyzed transcripts were segmented using a 200-nucleotide sliding window with a 100-nucleotide step. The same contact matrix is schematized by the heatmap in the right panel, where each cell reports the RIMEfull prediction score for the interaction between the corresponding RNA regions. **g**, Bar plot showing the proportion of RIME positive (score > 0.5) and negative (score < 0.5) predictions coinciding with windows supported by COMRADES chimeric reads. RIME predictions were conducted on all 200×200 nucleotide windows, with 100-nucleotide steps, derived from the contact matrices between NORAD and its 17 target mRNAs. Statistical significance was evaluated using Fisher's exact test. **h**, Heatmaps showing the RIMEfull predictions for the lnc-SMaRT–Spire1 transcript pair. Transcripts were segmented using a 200-nucleotide sliding window with a 100-nucleotide step. Each cell reports the RIMEfull prediction score for the interaction between the corresponding RNA windows. Previously validated interacting regions are highlighted with red boxes. **i**, The left heatmap shows the RIMEfull scores computed between pairs of 200-nucleotide NORAD segments. The lncRNA was segmented using a 200-nucleotide sliding window with a 100-nucleotide step. The same contact matrix is schematized by the heatmap in the right panel, where each cell reports the number of chimeric reads supporting the interaction between the corresponding regions, binned by deciles of the NORAD–NORAD chimeric read count distribution. **j**, Empirical cumulative distribution functions (ECDFs) of NORAD–NORAD RIMEfull scores after random shuffling across cells, shown for the 10 deciles defined by chimeric read counts. Each curve represents one decile (lighter colors indicate fewer reads; darker colors indicate more reads). **k**, Density plot of transcript-level Spearman correlations between intramolecular RIMEfull scores and average base-pairing probabilities predicted by RNAfold for 100 randomly selected human transcripts longer than 2,000 nucleotides (green). The null distribution (gray) was derived by randomly shuffling RIMEfull scores 1,000 times. Statistical significance of the difference between the two distributions was evaluated using two-sided Mann-Whitney U-test. **l**, Density plot of transcript-level Spearman correlations between DMS-Mapseq-derived accessibility scores of 200-nt RNA segments and RIMEfull scores calculated between such segments and their local context (green, see **Methods**). The null distribution (gray) was derived by randomly shuffling RIMEfull scores 1,000 times. Statistical significance of the difference between the two distributions was evaluated using two-sided Mann-Whitney U-test.
