## Supplementary material for "Decoding RNA–RNA Interactions: The Role of Low-Complexity Repeats and a Deep Learning Framework for Sequence-Based Prediction": Description of Supplementary Data

*File Name*: **Supplementary Data 1**
*Description***: Description of the RRI dataset derived from transcriptome-wide studies and its analysis.**

*Sheets*:

| RRI dataset | Description of the transcriptome-wide RNA duplex detection experiments analyzed in this study, including the number of RRIs retained at various data processing stages |
| --- | --- |
| RRI BEDPE | RNA-RNA interacting region pairs in BEDPE format |
| RRI patches | RRI patches in BED format, including the associated interactions |
| LCR RRI | Summary of the low-complexity repeats associated to RRIs and their classification |
| GO analysis | Summary of the GO term enrichment analysis of genes involved in RRI interactions mediated by specific low-complexity repeats |

*File Name*: **Supplementary Data 2**
*Description***: Description of the STAU1 hiCLIP-derived RNA heteroduplex dataset and its analysis. Results of RBP enrichment analyses.**

*Sheets*:

| RRI STAU1 hiCLIP BEDPE | RNA-RNA interacting region pairs derived from STAU1 hiCLIP in BEDPE format |
| --- | --- |
| STAU1 LCR RRI-NLEA | Results of the nucleotide-level repeat enrichment analysis on STAU1-bound heteroduplexes |
| RBP binding sites enrichment | Summary of RBP binding sites enrichment analysis on LCR-bearing RRI patches |
| RBP functional enrichment | Results of functional enrichment analysis on RBPs associated to LCR-bearing RRI patches |

*File Name*: **Supplementary Data 3**
*Description***: Information about the Lhx1os RNA pull-down experiments.**

*Sheets*:

| RNA pull-down probes | Describes the probes used in the RNA pull-down experiments |
| --- | --- |
| RNA-seq_preproc | Summarizes the number of reads at different stages of RNA-seq data preprocessing |
| Removed RNAs | Lists the RNAs removed in the cleaning step |
| RNA-seq_align_postproc | Summarizes the number of reads at in different stages of RNA-seq data alignment and post-processing |
| ODD vs Input | Describes the differential abundance analysis performed between Input samples and RNA pull-down samples obtained with ODD probe set |
| EVEN vs Input | Describes the differential abundance analysis performed between Input samples and RNA pull-down samples obtained with EVEN probe set |
| LacZ vs Input | Describes the differential abundance analysis performed between Input samples and RNA pull-down samples obtained with LacZ probe set |
| Primers | Describes the sequences of primers used to perform PCR analysis |

*File Name*: **Supplementary Data 4**
*Description***: RNA targets of TINCR and Lhx1os lncRNAs identified by pull-down experiments, with control transcripts.**

*Sheets*:

| TINCR pull-down RNA set | Describes the RNA targets and controls used for the analysis of TINCR interactors |
| --- | --- |
| Lhx1os pull-down RNA set | Describes the transcripts used for Lhx1os pull-down analysis |

*File Name*: **Supplementary Data 5**
*Description***: Performances of RIME, SPOT-RNAc and thermodynamics-based tools on different RRI sets.**

*Notes*:

To address class imbalance, negatives were randomly undersampled 100 times to match the number of positives, and all the metrics were averaged across iterations.

RIME DRP a: RIME model trained using only NN and NP samples as negatives (i.e., trained solely on the DRP task). Model selection was performed based on the F1 score computed over the RIME validation set.

RIME DRI a: RIME model trained using only PN samples as negatives (i.e., trained solely on the DRI task). Model selection was performed based on the F1 score computed over the RIME validation set.

RIME DRP b: RIME model trained using only NN and NP samples as negatives (i.e., trained solely on the DRP task). Model selection was performed based on the F1 score calculated over a validation set containing only NN and NP classes. Note: this model coincides with DRP a.

RIME DRI b: RIME model trained using only PN samples as negatives (i.e., trained solely on the DRI task). Model selection was performed based on the F1 score calculated over a validation set containing only PN classes.

*Sheets*:

| Psoralen Human AUC | AUC values calculated for the Psoralen Human test set, including the high-quality (HQ) subset |
| --- | --- |
| Psoralen Mouse AUC | AUC values calculated for the Psoralen Mouse test set, including the high-quality (HQ) subset |
| RIC-seq AUC | AUC values calculated for the RIC-seq test set, including the high-quality (HQ) subset |
| MARIO AUC | AUC values calculated for the MARIO test set |
| DRP RIME | RIME Accuracy, Precision, NPV, Recall, Specificity and F1 values calculated for the DRP task on different test sets |
| DRI RIME | RIME Accuracy, Precision, NPV, Recall, Specificity and F1 values calculated for the DRI task on different test sets |
